# Conserved early steps of stemmadenine biosynthesis

**DOI:** 10.1101/2025.11.23.690000

**Authors:** Mohamed O. Kamileen, Yoko Nakamura, Marlen Sigmund, Radhika Keshan, Veit Gabe, Sarah Heinicke, Maritta Kunert, Benke Hong, Ryan Alam, Gyumin Kang, Lorenzo Caputi, Sarah E. O’Connor

## Abstract

Stemmadenine acetate is a pivotal intermediate in the production of pharmacologically active monoterpene indole alkaloids. Here, we identify orthologs of stemmadenine acetate pathway genes (SGD, GS, GO, Redox1, Redox2, SAT) from *Tabernanthe iboga*. We characterize these enzymes in vitro, and additionally, we reconstitute stemmadenine acetate biosynthesis in *Nicotiana benthamiana*, comparing the formation of intermediates and shunt products that are produced when previously characterized orthologs from the related plant *Catharanthus roseus* are used. Ortholog pairs are catalytically indistinguishable, except in the case of GS. Surprisingly, the *T. iboga* ortholog catalyzes formation of an alternate stereoisomer 19*Z*-geissoschizine, seeding a low-flux Z-series in heterologous reconstitution systems in vitro and in planta. We additionally characterize the major shunt products that arose during reconstitution of stemmadenine acetate biosynthesis. We show that the substrate promiscuity of Redox2 results in formation of the shunt products 16(*R/S*)-isositsirikines, hampering pathway flux and yields. Additionally, we show that stemmadenine can be oxidized by endogenous *N. benthamiana* enzymes, leading to the shunt product condylocarpine.

Nevertheless, we could produce stemmadenine at a 6 mg yield from 19*E*-geissoschizine by heterologous expression in *N. benthamiana*. Overall, we highlight the prospects for milligram production of important biosynthetic intermediates in *N. benthamiana*.

## Introduction

Stemmadenine acetate is a pivotal intermediate in the biosynthesis of structurally diverse and bioactive monoterpene indole alkaloids (MIAs). Stemmadenine acetate is at a branch point that leads to numerous high-value alkaloids, including the anticancer agent vinblastine (1, 2) as well as the psychoactive ibogaine (3) (Fig. 1). The conversion of strictosidine, the universal MIA precursor, to stemmadenine acetate is well characterized in *Catharanthus roseus* (1, 2, 4) (Fig. 1). In contrast, the corresponding pathway in *Tabernanthe iboga*, a medicinal plant species producing a source of biologically active iboga-type alkaloids, has remained unexplored. In the canonical early pathway, strictosidine β-D-glucosidase (SGD) hydrolyzes strictosidine to the reactive aglycone (5), which is reduced by the medium-chain dehydrogenase/reductase (MDR) geissoschizine synthase (GS) to form 19*E*-geissoschizine (6, 7). The cytochrome P450 enzyme, geissoschizine oxidase (GO) converts this to a short-lived iminium intermediate (dehydroprekuammicine) (6, 8), which the MDR Redox1 subsequently reduces, and then the aldehyde is further reduced by the aldo-keto reductase (AKR) Redox2 to yield stemmadenine (2). The BAHD acetyl transferase, stemmadenine acetyltransferase (SAT), finally acetylates stemmadenine to produce stemmadenine acetate (Fig. 1) (1, 2).

**Figure 1.**
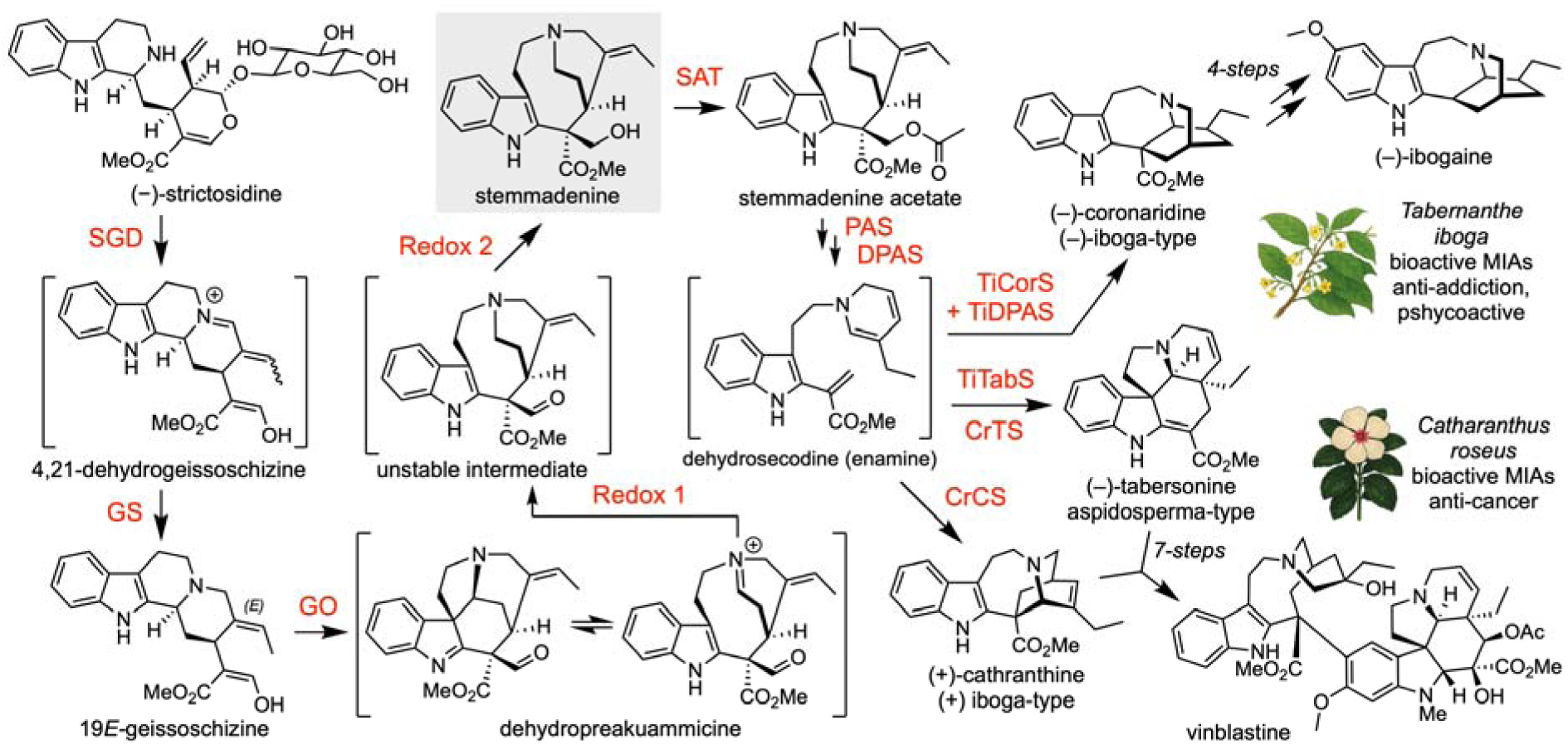
Biosynthesis of stemmadenine acetate. Overview of the early MIA pathway from strictosidine to stemmadenine acetate. Stemmadeine In *C. roseus*, stemmadenine is a key precursor en route to the anticancer agent vinblastine. The closely related medicinal plant *T. iboga* channels stemmadenine towards bioactive alkaloids, including the psychoactive, anti-addictive compound ibogaine.

Here we identify and functionally characterize the *T. iboga* orthologs SGD, GS, GO, Redox1, Redox2, and SAT, validating activities in vitro and in planta. We rebuild the route to stemmadenine from strictosidine and 19*E*-geissoschizine, benchmark *T. iboga* enzyme activities against *C. roseus* enzymes, and deploy multi-gene constructs to recapitulate stemmadenine biosynthesis in *N. benthamiana* for scalable bioproduction at a ca. 5 milligram scale. Alongside conserved pathway logic, we pinpoint steps at which the biochemical flux is diverted by pathway enzymes or endogenous host activities, yielding reductive and oxidative side products. Notably, *T. iboga* GS unexpectedly favors the formation of 19*Z*-geissoschizine over the on-pathway 19*E* isomer, seeding a parallel but low-flux Z-series of downstream intermediates in heterologous pathway reconstitution. Thus, even biosynthetic pathways with clear orthologous steps can yield unexpected outcomes.

## Results

### Identification of stemmadenine acetate pathway orthologs in *Tabernanthe iboga*

We performed LC-MS based untargeted metabolomics of *T. iboga* shoots (young leaf, mature leaf, and stem) and roots, and revealed that MIA intermediates, including (–)-strictosidine, (+)-stemmadenine acetate, (–)-pachysiphine, (–)-tabersonine, and (+)-pseudo-vincadifformine, accumulate predominantly in young leaves (Fig. 2A, S1). Guided by this tissue distribution, we searched for orthologs of the functionally characterized *C. roseus* SGD, GS, GO, Redox1, Redox2, and SAT genes in the *T. iboga* transcriptome. We similarly queried the sarpagan bridge enzyme (SBE) (8–10), which is a cytochrome P450 that, like GO, acts on 19*E*-geissoschizine. Candidates were prioritized by (i) transcript abundance in young leaves, where pathway intermediates accumulate (Fig. 2B), and (ii) Pearson correlation to known *T. iboga* biosynthetic genes for aspidosperma-, iboga-, and pseudo-aspidosperma-type scaffold biosynthesis (Fig. 2C, S2). Candidate identities were further supported by maximum-likelihood phylogenetic analysis (Fig. S3). This integrative strategy yielded a single high-confidence gene for each queried step.

**Figure 2.**
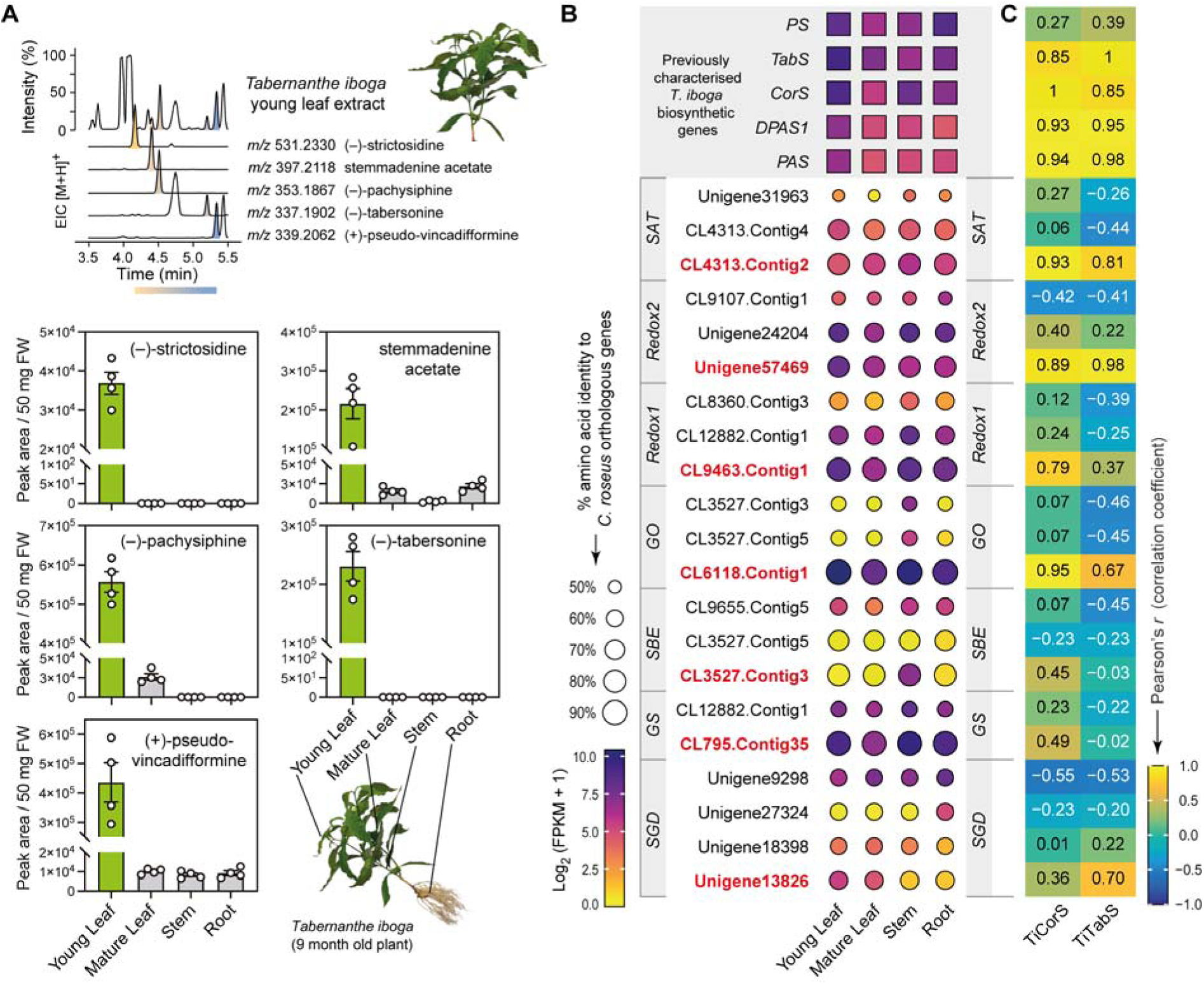
Metabolite analysis and candidate selection. (**A**) Metabolite analysis of *T. iboga* tissues. (**B**) Relative gene expression profiles and amino acid identity of *T. iboga* candidate genes orthologous to *C. roseus* early pathway enzymes. Genes selected for functional characterization are highlighted in red. (**C**) Pearson correlation of gene candidates with previously characterised MIA biosynthetic enzymes of *T. iboga*. Error bars represent mean ± standard deviation (n = 4 biological replicates).

### Characterization of *T. iboga* stemmadenine acetate biosynthesis from 19*E*-gesissoschizine

We cloned early-pathway gene orthogs from *T. iboga* (GO, Redox1, Redox2, SAT) into expression vectors and screened for enzymatic activity by *Agrobacterium*-mediated transient expression in *Nicotiana benthamiana*. To validate the gene function of these orthologs, we employed stepwise pathway reconstitution approaches, along with co-infiltration of the GO substrate, 19*E*-geissoschizine in *N. benthamiana* leaf disks. LC-MS analysis showed that previously characterized *C. roseus* enzymes and the *T. iboga* orthologs produced the same sequence of intermediates at each step leading to stemmadenine acetate, the product of SAT (Fig. 3A, S4). We also observed precondylocarpine acetate, the oxidation product of stemmadenine acetate; previous work has shown that an endogenous *N. benthamiana* enzyme or enzymes catalyzes this oxidation (1, 11, 12) (Fig. 3A, S4). Direct benchmarking revealed no significant differences in intermediate or final product levels between *C. roseus* and *T. iboga* ortholog sets (unpaired t-test, *P* > 0.05; *n* = 3; Fig. 3A, S4).

**Figure 3.**
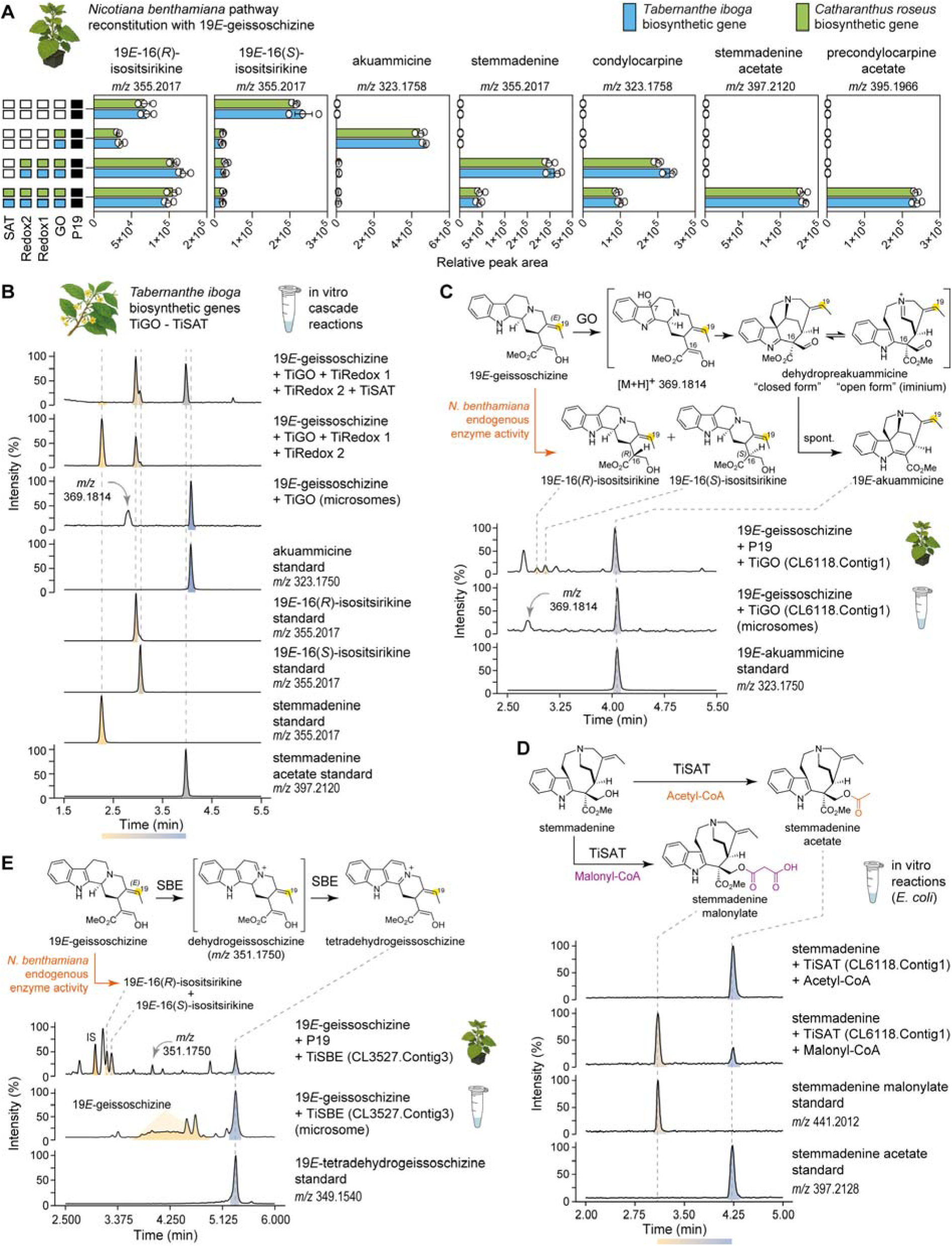
Biosynthesis of stemmadenine acetate from 19E-geissoschizine in *T. iboga*. **(A)** *N. benthamiana* pathway reconstitution of stemmadenine acetate from the substrate 19*E*-geissoschizine (unpaired t-test, *n =* 3 biological replicates, *P* > 0.05). **(B)** In vitro characterization of *T. iboga* early pathway enzymes by cascade reactions to stemmadenine acetate from 19*E*-geissoschizine substrate. **(C)** In vitro characterization of *T. iboga* geissoschizine oxidase (GO). **(D)** In vitro characterization of *T. iboga* stemmadenine acetyl transferase (SAT). **(E)** Characterization of *T. iboga* sarpagan bridge enzyme (SBE). Error bars represent mean ± standard deviation.

To corroborate these activities independently of the *N. benthamiana* host, we performed in vitro assays with isolated enzymes. TiGO was expressed in yeast, and the microsomal fraction was isolated. TiRedox1, TiRedox2, and TiSAT were produced recombinantly in *E. coli* and purified by nickel affinity chromatography. Using stepwise and coupled cascade reactions with 19*E*-geissoschizine as substrate, LC-MS analysis with authentic standards confirmed the expected series of intermediates, including stemmadenine and the downstream acetylated product, stemmadenine acetate (Fig. 3B, S5). Yeast microsomes harboring TiGO converted 19*E*-geissoschizine to akuammicine, and we also observed a compound with a mass consistent with the proposed short-lived oxyindole intermediate (*m/z* 369.1814) (2) (Fig. 3C, S6). In TiGO and TiRedox1 coupled reactions with 19*E*-geissoschizine in vitro, we observed the formation of two products at *m/z* 325.1910, consistent with 16(*R/S*)-deshydroxymethyl stemmadenine; spontaneous rearrangement products of the reduced dehydropreakuammicine iminium intermediate (Fig. S5). These products mirrored previously reported *C. roseus* Redox1 (2) in vitro and indicated that TiRedox1 performs the analogous MDR reduction in *T. iboga*. Coupled in vitro assays of purified *T. iboga* Redox2, GO, and Redox1 enzymes with 19*E*-geissoschizine generated stemmadenine, establishing TiRedox2 as the AKR that reduces the transient unstable aldehyde of TiRedox1 to the stable alcohol product, stemmadenine (Fig. 3A-B, S4, S5). Because the GO, GO + Redox1 intermediates are unstable and not isolable, TiRedox1 and TiRedox2 activities were necessarily inferred from coupled cascades, consistent with the *in planta* reconstitutions in *N. bethanamiana*.

Finally, recombinant *T. iboga* SAT converted stemmadenine to stemmadenine acetate in the presence of acetyl-CoA in vitro, with retention time and MS^2^ spectra identical to authentic standard (Fig. 3D, S7). When the CoA donor was switched to malonyl-CoA, we detected a product consistent with stemmadenine malonylate (*m/z* 441.2012) supported by MS^2^ spectral assignment (Fig. 3D, S7). We also tested the *T. iboga* SBE ortholog both in vitro using purified yeast microsomes and in vivo using TiSBE expressed in *N. benthamiana* with 19*E*-geissoschizine as the substrate. In each context, TiSBE catalyzed the formation of tetradehydrogeissoschizine (8), which matched an authentic standard by retention time and MS^2^ fragmentation (Fig. 3E, S8, S9).

### Multigene constructs do not alter the product profile

After validating the function of *T. iboga* genes GO, Redox1, Redox2, and SAT, we assembled two multigene constructs (pMGC) for transient expression in *N. benthamiana*. Construct 1 (pMGC1) comprised three transcriptional units encoding TiGO, TiRedox1, and TiRedox2, and construct 2 (pMGC2) comprised four transcriptional units encoding TiGO, TiRedox1, TiRedox2, and TiSAT (Fig. 4). Each pMGC was benchmarked against co-infiltration of single strains carrying the same genes. Leaf disk assays were performed with equal amounts of 19*E*-geissoschizine, and LC-MS analysis showed closely matching product profiles in both the multigene and co-infiltration formats. For pMGC1, both formats produced stemmadenine with no statistically significant difference in metabolite levels (Fig. 4A, S10; unpaired t-test, *n =* 3, *P* > 0.05). We quantified stemmadenine titers from leaves transiently expressing pMGC1 at *Agrobacterium* densities OD_600_ = 1.0 or 2.0, followed by exogenous feeding of 19*E*-geissoschizine at 0.05, 0.10, 0.25, or 0.50 mM. We could detect 150 ng/mg of product with 0.5 mM substrate (Fig. 4B, S11). Guided by these results, we executed a scale-up using pMGC1, in which two leaves of 50 *N. benthamiana* plants were infiltrated, and four days later, the same leaves were substrate-fed with 19*E*-geissoschizine, where each leaf received 1 ml at 0.5 mM. Leaves were harvested 14-16 hours after substrate delivery, and purification by preparative HPLC afforded 6 mg of stemmadenine, which was confirmed by NMR analysis (Fig. S12, Table NMR S1). Similarly, pMGC2 and its co-infiltration equivalent yielded stemmadenine acetate and precondylocarpine acetate at comparable levels (Fig. 4C, S13; unpaired t-test, *n =* 3, *P* > 0.05).

**Figure 4.**
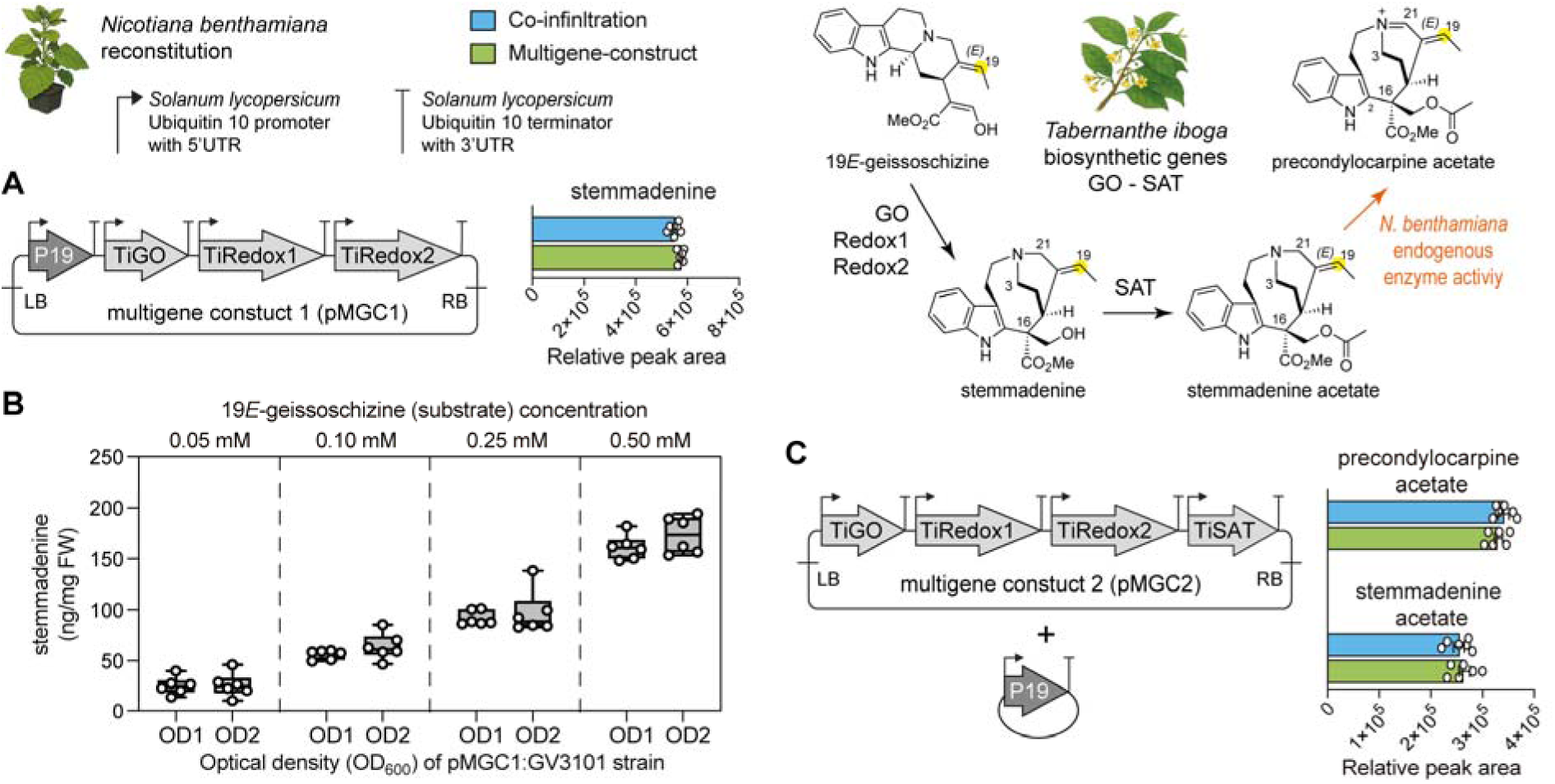
Multigene constructs for upscaling stemmadenine and stemmadenine acetate biosynthesis in *N. benthamian* from 19*E*-geissoschizine. (A) Multigene construct 1 (pMGC1) for stemmadenine biosynthesis. **(B)** Quantification of stemmadnine produced by pMGC1 at varying strain density and substrate concentrations. **(C)** Multigene construct 2 (pMGC2) for stemmadenine acetate biosynthesis. Error bars represent mean ± standard deviation (unpaired t-test, *n =* 6 biological replicates, *P* > 0.05).

### Pathway reconstitution yields undesired reductive and oxidative side products

During reconstitution of the stemmadenine acetate pathway in *N. benthamiana* and in coupled enzyme in vitro cascade reactions with 19*E*-geissoschizine, we consistently detected two side-products at *m/z* 355.2017, in addition to stemmadenine (*m/z* 355.2017) (Fig. S4, S5, S10, S13). These compounds were observed when 19*E*-geissoschizine (*m/z* 353.1862) was infiltrated into wild-type or P19-control plants, suggesting that 19*E*-geissoschizine was reduced by endogenous *N. benthamiana* enzymes (Fig. 5A, S14). However, these same compounds were also observed in in vitro assays only when 19*E*-geissoschizine was incubated with enzymes in combination with Redox2 (Fig. S5). To structurally characterize these products, we incubated 19*E*-geissoschizine with purified TiRedox2 from *E. coli* in the presence of NADPH and isolated the two products by preparative HPLC for NMR structure elucidation. The products were identified as 19*E*-16(*R*)-isositsirikine and 19*E*-16(*S*)-isositsirikine (Fig. 5A; Fig. S15, S16; Table NMR S2, S3). Chemical reduction of 19*E*-geissoschizine with NaBH_4_ produced the identical diastereomeric pairs with matching retention times and MS² spectra (Fig. 5B, S17).

**Figure 5.**
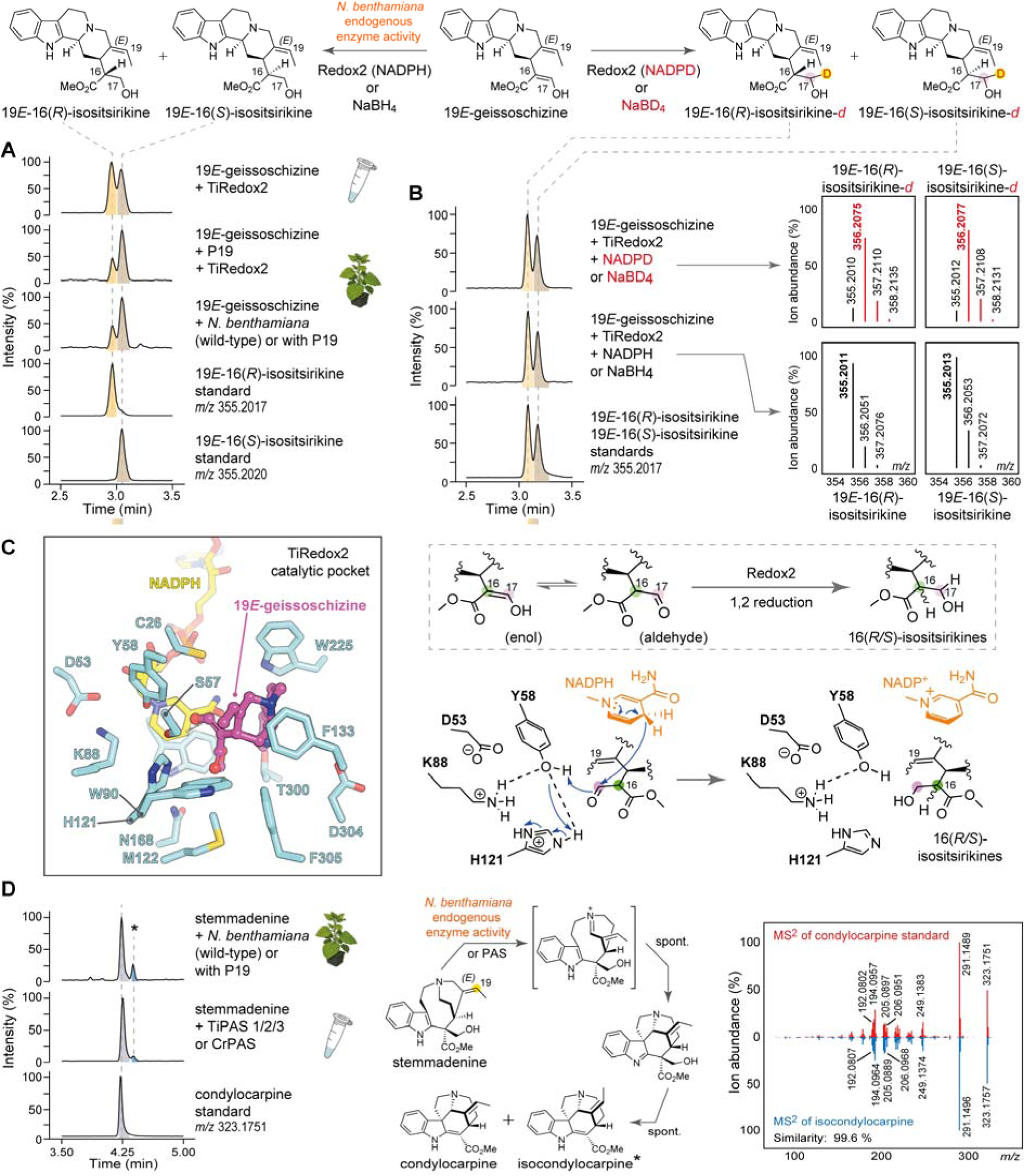
Pathway reconstitution results in undesired byproducts. **(A)** Formation of 19*E*-16(*R/S*) isositsirikine diastereomers by *N. benthamiana* endogenous enzymes and the action of aldo-keto reductase enzyme Redox 2. **(B)** Generation of deuterium labelled 19*E*-16(*R/S*) isositsirikines by enzymatic NADPD labelling and chemical reduction with NaBD_4_. **(C)** Catalytic pocket of AlphaFold v3 *T. iboga* Redox2 model with 19*E*-geissoschizine docked into the active site. Proposed catalytic mechanism of 19*E*-geissoschizine reduction by the enzyme Redox2, along with the catalytic tetrad. **(D)** Generation of condylocarpine and isocondylocarpine isomeric products from stemmadenine by *N. benthamiana* endogenous oxidative enzyme/s or downstream MIA biosynthetic enzyme PAS.

To probe the mechanism by which Redox2 can act to reduce 19*E*-geissoschizine, we performed Redox2 reactions with in situ–generated pro-(*R*)-NADPD. Both 19*E*-16(*R/S*)-isositsirikine products exhibited a +1 Da mass shift (*m/z* 356.2077), consistent with single deuterium incorporation (Fig. 5B, S17). Complementary NaBD_4_ reductions likewise yielded mono-deuterated diastereomers (Fig. 5B, S17). Following scale-up with pro-(*R*)-NADPD, isolation and NMR analysis established that the deuterium resides at C17 (Fig. S18, S19; Table NMR S4, S5). Guided by the phylogenetic placement of TiRedox2 (Fig. S2), we examined a multiple-sequence alignment of TiRedox2 alongside functionally characterized plant aldo-keto reductases (AKRs) *C. roseus* Redox2 (2) and *Alstonia scholaris* rhazimal reductase (RHR) (13) from MIA biosynthesis and *Papaver somniferum* codeinone reductase (COR) (14) from benzylisoquinoline alkaloid (BIAs) pathway (Fig. S20).

The alignment revealed the canonical AKR catalytic tetrad (Tyr, Lys, His, and Asp) (15). We generated an AlphaFold model of TiRedox2 with NADPH bound and docked 19*E*-geissoschizine into the active site (Fig. 5C, S21). The top-ranked docking poses place Y58 in close proximity (3 Å) to the carbonyl group of 19*E*-geissoschizine, forming a hydrogen bond to the aldehyde moiety and positioning the substrate for either a 1,2 or 1,4 reduction to generate 19*E*-16(*R/S*)-isositsirikine diasteriomers (Fig. 5C, S22).

In addition to the 19*E*-16(*R/S*)-isositsirikine diasteriomers, we detected an unknown product at *m/z* 323.1751 during pathway reconstitution to stemmadenine or stemmadenine acetate. This product was observed only during *N. benthamiana* reconstitution, and not in vitro coupled reactions (Fig. S4, S5). Therefore, we hypothesized that endogenous *N. benthamiana* enzymes catalyze formation of this product. Indeed, when stemmadenine was incubated with leaf disks from wild-type or P19-control plants, the same compound (*m/z* 323.1751) was observed (Fig. 5D, S23). We scaled the reaction by infiltrating stemmadenine into wild-type *N. benthamiana* leaves, isolated this product by preparative HPLC, and structurally characterized it by NMR. The structure was established to be condylocarpine (Fig. S24; Table NMR S6). Prior work has shown that stemmadenine acetate is oxidized to precondylocarpine acetate by precondylocarpine acetate synthase (PAS) (1, 3, 11), a flavin-containing berberine bridge-like (BBL) enzyme (16), and that native *N. benthamiana* BBLs can perform an analogous oxidation (1, 11). With this in mind, we assayed previously reported PAS enzyme from *T. iboga* or *C. roseus* with stemmadenine and observed a major product matching condylocarpine together with a minor isomer exhibiting an indistinguishable MS^2^ spectrum, here designated as isocondylocarpine (Fig. 5D, S23).

### *T. iboga* geissoschizine synthase (GS) redirects flux to an alternative geissoschizine isomer

Having reconstituted the stemmadenine acetate pathway from 19*E*-geissoschizine, we next set out to reconstitute the route from strictosidine, an earlier MIA precursor (17). We cloned and transiently expressed SGD and GS orthologs from *T. iboga*, together with the downstream genes *T. iboga* GO, Redox1, Redox2, and SAT in *N. benthamiana*. In parallel, we performed the same reconstitution with *C. roseus* as a positive control and to benchmark the levels of biosynthetic products and intermediates. Four days post-infiltration, strictosidine was supplied exogenously directly into the infiltrated leaves, and tissues were analyzed by LC-MS. Unexpectedly, tissues containing *T. iboga* enzymes yielded markedly 6× lower amounts of the final products stemmadenine acetate and precondylocarpine acetate compared to reconstitution with the *C. roseus* orthologs (Fig. 6A, S25, unpaired t-test, *n* = 3, *P* < 0.0001), motivating us to closely examine the steps upstream of 19*E*-geissoschzine, SGD, and GS.

**Figure 6.**
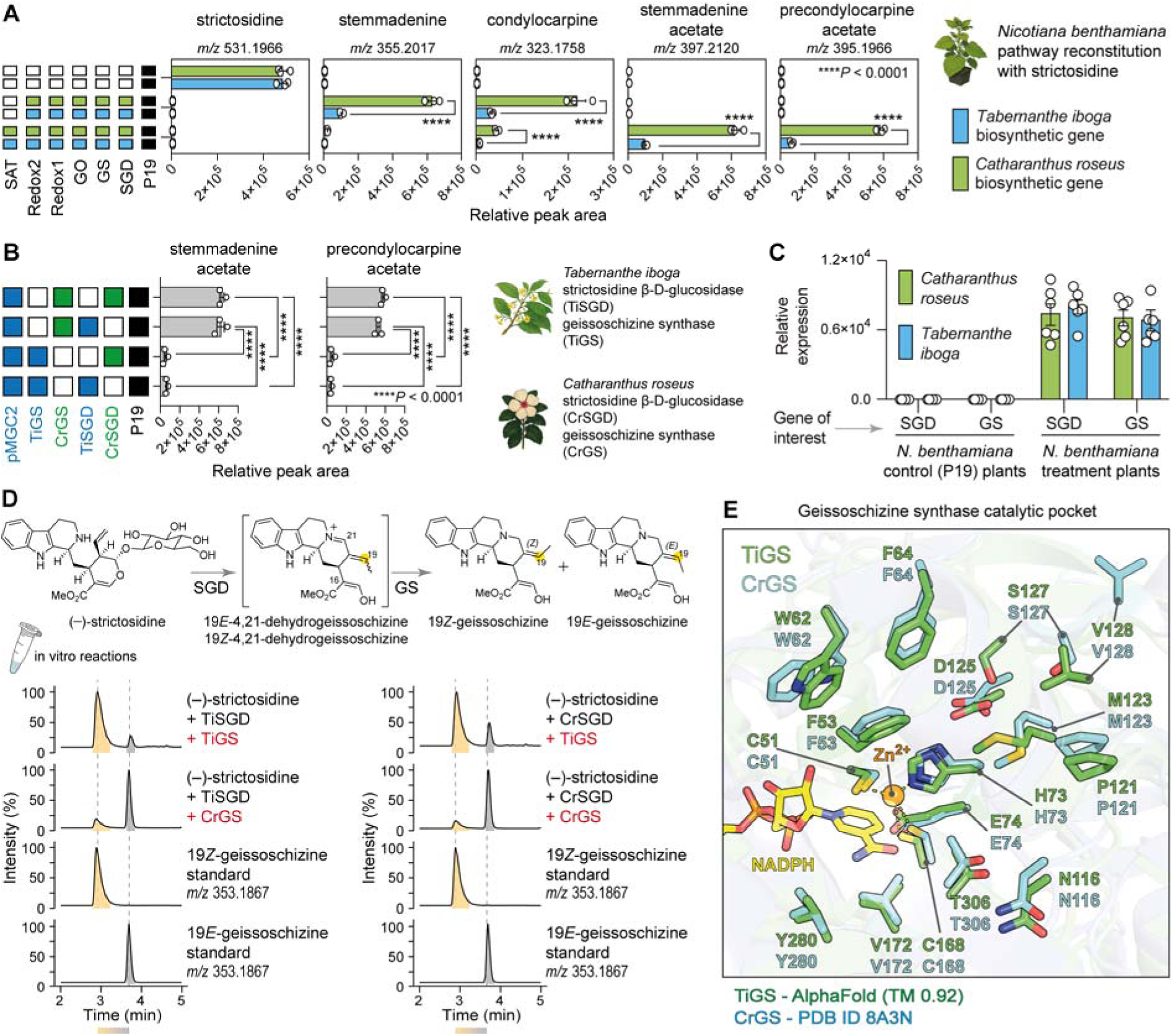
*T. iboga* stemmadenine acetate biosynthetic pathway reconstitution from strictosidine. **(A)** Pathway reconstitution of stemmadenine acetate by *T. iboga* enzymes benchmarked against *C. roseus* early-pathway enzymes (unpaired t-test, *n =* 3 biological replicates, \*\*\*\**P* < 0.0001). **(B)** Combinatorial biosynthetic pathway reconstitution of stemmadenine acetate using *C. roseus* SGD and GS enzymes (one-way ANOVA, *n =* 3 biological replicates, \*\*\*\**P* < 0.0001). **(C)** Relative expression levels of *T. iboga* and *C. roseus* SGD and GS genes transiently expressed in *N. benthamiana* (unpaired t-test, *n =* 6 biological replicates, *P* > 0.05). **(D)** In vitro coupled reactions of SGD and GS from *T. iboga* and *C. roseus*. **(E)** Catalytic pocket comparison between the crystal structure of *C. roseus* GS (PDB 8A3N) and the AlphaFold v3 model of *T. iboga* GS. Where appropriate, peak areas are calculated and presented relative to an internal standard. Error bars represent mean ± standard deviation.

We transfected *N. benthamiana* with the *T. iboga* multigene module pMGC2 (TiGO, TiRedox1, TiRedox2, and TiSAT), along with either SGD and GS from *T. iboga* or with the corresponding orthologs from *C. roseus*. We found that substituting CrGS for TiGS restored efficient production of stemmadenine acetae and precondylocarpine acetate (Fig. 6B, S26). We analyzed the relative gene expression levels of SGD and GS in *N. benathamiana* leaf tissues by qPCR. The results showed comparable expression of *T. iboga* and *C. roseus* genes, excluding expression differences as the cause of the difference in titer (Fig. 6C). Subcellular distribution of *T. iboga* early-pathway enzymes (TiSGD-TiSAT) in both *N. benthamiana* leaves and *C. roseus* flower petals showed localization patterns previously reported for *C. roseus* orthologs (5, 6, 18, 19), ruling out mislocalization as a cause for the low activity and product yields (Fig. S27, S28).

We then characterized the activity of *T. iboga* GS in vitro by purifying the enzymes from *E. coli* and conducting coupled enzyme reactions with strictosidine, mirroring the in planta combinations. Surprisingly, *T. iboga* GS catalyzed the formation of 19*Z*-geissoschizine with only minor levels of 19*E*-geissoschizine, the stereoisomer that is on-pathway (Fig. 6D, S29). In contrast, *C. roseus* GS favoured the production of 19*E*-geissoschizine with low levels of 19*Z*-geissoschizine (Fig. 6D, S29). Intrigued by this stereochemical divergence, we compared *T. iboga* GS to functionally characterized GS enzymes (6, 13, 20, 21) (Fig. S30). Multiple sequence alignments and AlphaFold-guided structural comparisons revealed no obvious differences within the active site (Fig. S31). We next asked whether *T. iboga* encodes an alternative 19*E*-geisoschizine-producing MDR. To investigate this, we screened 32 *T. iboga* MDRs identified from phylogenetic analysis, including previously characterized *T. iboga* dihydroprecondylocarpine acetate synthase (DPAS1 and 2) (3). These were tested in the *N. benthamiana* system coupled with TiSGD and pMGC1 (TiGO, TiRedox1, and TiRedox2) using strictosidine as the substrate. However, none of the tested MDR candidates substituted for *T. iboga* GS (CL795.Contig35) led to production of stemmadenine (Fig. S32).

### 19*Z*-geissoschizine seeds a low-flux Z-series of pathway intermediates

We reconstituted the pathway using chemically synthesized 19*Z*-geissoschizine (22) as the substrate in *N. benthamiana* with either of the aforementioned multigene modules, pMGC1 and pMGC2, in separate experiments, and compared the outcomes with those obtained with 19*E*-geissoschizine substrate under identical conditions. With 19*Z*-geissoschizine, we detected a family of products and intermediates at low yields, whose [M+H]^+^ exact masses matched those of the 19*E*-configured standards but eluted with systematic retention time offsets consistent with the *Z*-configuration (Fig. 7A, S33). MS^2^ spectra of these products closely mirrored those of the 19*E*-derived authentic standards (Fig. S34). To decouple host effects, we performed coupled in vitro reactions with purified *T. iboga* enzymes GO, Redox1, Redox2, and SAT enzymes using 19*Z*-geissoschizine. The product profile recapitulated the in planta *N. benthamiana* reconstitution results, including the low-level formation of putative 19*Z*-stemmadenine and 19*Z*-stemmadenine acetate, with MS^2^ features matching the 19*E*-configured standards (Fig. S35, S34).

**Figure 7.**
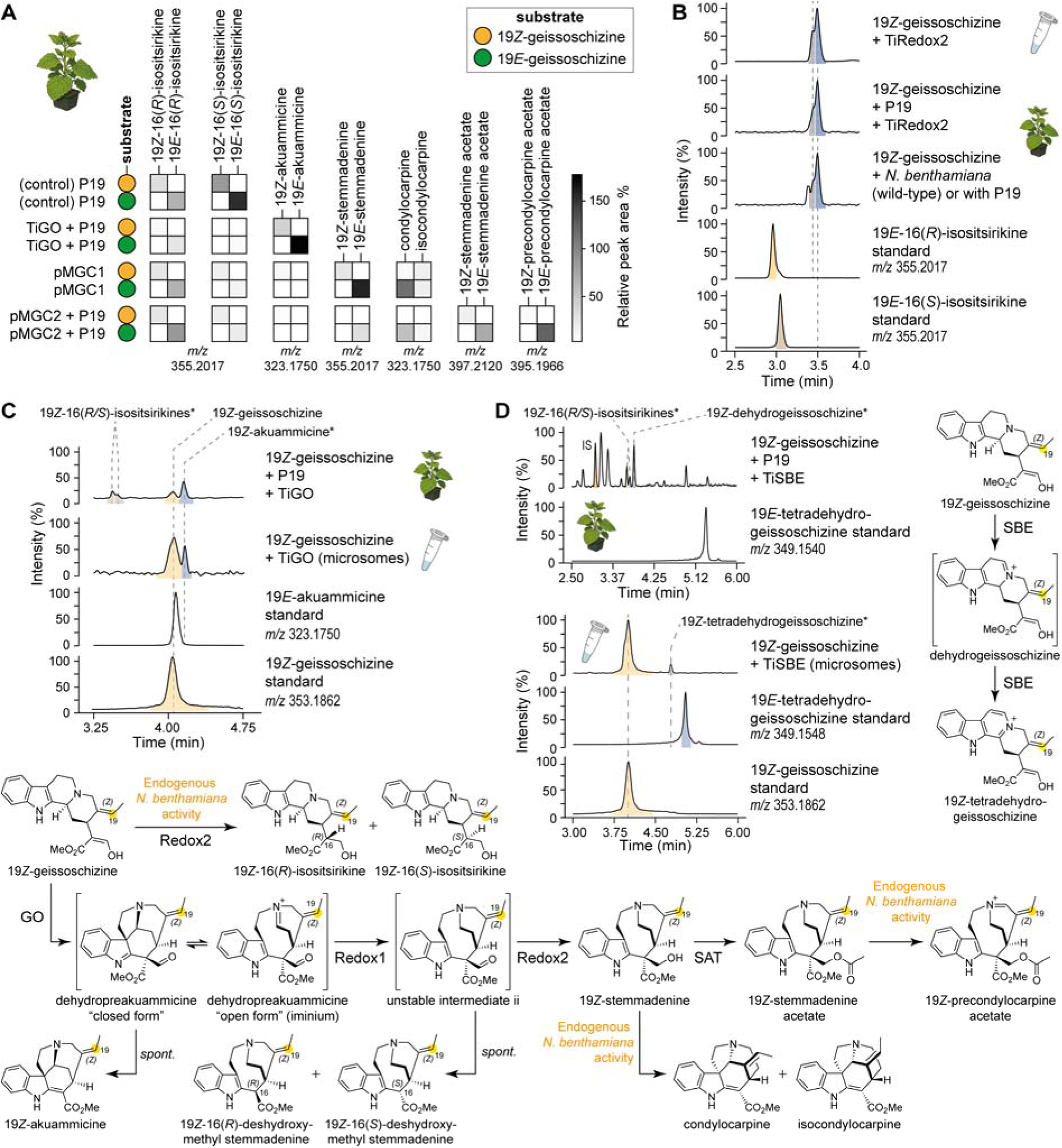
19*Z*-geissoschizine yields low levels of early pathway intermediates. (**A**) Heat map comparing abundances of 19*Z*-configured intermediates produced by *T. iboga* pathway reconstitution in *N. benthamian* when assayed with 19*Z*-geissoschizine substrate with those of the corresponding 19*E*-series assayed with 19*E*-geissoschizine under identical conditions. Abundances in the heat map are presented as peak area relative to the internal standard, averaged from three biological replicates. **(B)** Assignment of 19*Z*-16(*R/S*)-isositsirikine diasteriomers generated by endogenous *N. bethamiana* activity and by Redox2 in vitro from 19*Z*-geissoschizine. **(C)** Assignment of 19*Z*-akuammicine formed by GO in *N. bethamiana* and GO-expressed yeast microsomes from 19*Z*-geissoschizine. **(D)** Characterization of *T. iboga* sarpagan bridge (SBE) activity on 19Z-geissoschizine expressed in *N. benthamiana* and yeast microsomes.

Consistent with observations from the 19*E*-configured pathway, we also detected reductive and oxidative side products in the 19*Z*-series. Two LC-MS features with *m/z* 355.2017 were attributed to the compounds 19*Z*-16(*R*)-isositsirikine and 19*Z*-16(*S*)-isositsirikine (Fig. S33, S35). To verify these assignments, we assayed purified TiRedox2 enzyme with 19*Z*-geissoschizine in vitro, and with *N. benthamiana* leaf disks, both wild-type and those expressing the P19 suppressor. In both systems, the reactions yielded products with identical exact masses and MS² fragmentation spectra, and their retention times were shifted as expected, attributable to the *Z*-configuration (Fig. 7B, Fig. S36). Similarly, oxidation of 19*Z*-geissoschizine by TiGO produced a feature at *m/z* 323.1750, which is consistent with the formation of 19*Z*-akuammicine. This result was confirmed in two experimental settings, TiGO-expressing yeast microsomes and *N. benthamiana* leaf disks transiently expressing TiGO, both of which generated a product in which the exact mass and MS² spectrum were identical to that of authentic akuammicine, but with a retention time offset characteristic of the *Z*-isomer (Fig. 7C, Fig. S37). Finally, in vitro assays with TiSBE microsomes converted 19*Z*-geissoschizine to a product consistent with 19*Z*-tetradehydrogeissoschizine (*m/z* 349.1548), supported by MS² and a *Z*-shifted retention time (Fig. 7D, Fig. S38). In *N. benthamiana*, however, we did not detect the 19*Z*-tetradehydro species; instead, we observed accumulation of a compound consistent with 19*Z*-dehydrogeissoschizine (Fig. 7D, Fig. S39).

## Discussion

We identified and functionally validated *T. iboga* orthologs that convert strictosidine, the universal precursor to all MIAs, to stemmadenine acetate. The activites of these orthologs were benchmarked against previously characterized *C. roseus* enzymes (2, 4). In both *N. benthamiana* and in vitro assays, ortholog pairs of GO, Redox1, Redox2, and SAT produced the same products, indicating strong conservation of catalytic roles and a shared early-pathway logic (Fig. S4, S5). Interestingly, in addition to acetyl-CoA, TiSAT could accept malonyl-CoA in vitro to form stemmadenine malonylate, but *in planta*, only the acetylated product accumulated (Fig. 3D, S4, S7).

Upstream of 19*E*-geissoschizine, we observed a stereochemical fork that redirects flux. TiGS preferentially forms 19*Z*-geissoschizine instead of the expected 19*E* isomer (Fig. 5D, S29). Although the 19*Z* isomer can traverse through the early pathway enzymes to produce a parallel set of *Z*-configured intermediates, overall flux through this branch was low in both *N. benthamiana* reconstitutions and in vitro (Fig. 7A, S25, S33, S35). Consistent with this observation, *T. iboga* tissue extracts did not reveal detectable levels of these19*Z*-configured intermediates, suggesting that 19*Z*-geissoschizine is not physiologically relevant in planta.

Multiple-sequence alignment comparisons and modeling of the active-site revealed no obvious active-site substitutions to predetermine the *E*/*Z* outcome of geissoschizine (Fig. S30, S31). We could not identify an alternative 19*E*-producing MDR in *T. iboga* (Fig. S32). Together, these data explain the lower titers of stemmadenine obtained when starting from strictosidine with the *T. iboga* set (Fig. 6A, 6B, S25, S26). These results suggest an as-yet-undiscovered mechanism by which 19*E*-geissoschizine is produced in *T. iboga*.

Pathway reconstitution in *N. benthamiana* was also accompanied by the production of side products. The substrate, and pathway intermediate 19*E*-geissoschizine was reduced to 19*E*-16(*R/S*)-isosistsirikine diasteriomers (Fig. S4, S25), and the 19*Z*-geissoschizine isomer analogously yielded 19*Z*-16(*R/S*)-isositsirikines (Fig. S25, S33). The same 16(*R/S*)-isosistsirikine products arose in coupled in vitro cascade reactions containing TiRedox2 (Fig. S5, S35), demonstrating that TiRedox2 can directly reduce both 19*E* and 19*Z* geissoschizine (Fig. S14, S36) and suggesting that endogenous AKR enzymes in *N. benthamiana* can intercept MIA intermediates bearing carbonyl functional groups such as aldehydes and ketones. Deuterium labeling with pro-(*R*)-NADPD placed the isotopic label at C17 (Fig. S17-19), consistent with an AKR “push-pull” mechanism (15, 23) and favouring 1,2-reduction of the substrate aldehyde to generate the alcohol products (Fig. 5C, S22). We also observed endogenous oxidation of stemmadenine to condylocarpine and isocondylocarpine in the *N. benthamiana* system (Fig. 5D, S23). Incubation with the downstream MIA biosynthetic enzyme PAS (1, 3) reproduced this transformation in vitro, implicating BBL oxidases in the undesired oxidation of stemmadenine in *N. benthamiana* (Fig. S23). Practically, harvest timing proved critical: sampling within 24 h of substrate feeding minimized background conversions, whereas extended incubations increased losses and produced complex profiles. With these optimizations, despite the formation of these side products, we enabled the milligram-scale bioproduction of stemmadenine; a valuable MIA precursor that lacks a direct synthetic route; from an accessible substrate in *N. benthamiana* chassis highlighting the potential for larger-scale production of complex alkaloids.

### Experimental procedures Chemicals and reagents

All chemicals, solvents, and molecular biology reagents and kits were obtained from commercial suppliers. Strictosidine was prepared from tryptamine (Sigma-Aldrich, Cat. No. 193747) and secologanin (Biosynth, Cat. No. FS65469) using *Catharanthus roseus* strictosidine synthase (CrSTR) as previously described (24). 19*E*-geissoschizine and 19*Z*-geissoschizine were synthesized according to established protocols (22, 25). Akuammicine was enzymatically generated as previously reported (6). Tetradehydrogeissoschizine was synthesised following a previously reported method (8). Stemmadenine was enzymatically produced in *N. benthamiana* as described in this study. Stemmadenine acetate and precondylocarpine acetate were prepared from stemmadenine as previously described (1).

### Plant material and growth conditions

*Tabernanthe iboga* plants were maintained as previously described (3). *Catharanthus roseus* (L.) G. Don. (var. Little Bright Eyes, LBE) seeds were germinated in standard potting soil and grown in a climate-controlled chamber under a 16=h:=8=h, light:=dark photoperiod at 23 °C and 60% relative humidity. Flower petals from 6- to 8-month-old *C. roseus* LBE plants were used for transient transformation experiments.

*Nicotiana benthamiana* plants used for *Agrobacterium*-mediated transient expression were cultivated in a glasshouse on a low-nutrient F1 compost soil mix under a 16=h:=8=h, light:=dark cycle at 22 °C and 55% relative humidity. Plants were used for infiltration 3-4 weeks after germination.

### Candidate selection

Candidate genes were mined from a previously generated *T. iboga* transcriptome (3). Full-length open reading frames were retrieved by BLAST searches using functionally characterized *C. roseus* early-pathway enzymes (SGD-SAT) as queries. Orthologs were prioritized by co-expression analysis with previously characterized *T. iboga* genes downstream of stemmadenine acetate (3, 12, 26), and by tissue-resolved expression to favor candidates enriched in tissues where early pathway MIA intermediates accumulate.

Protein sequences of shortlisted hits were aligned with their characterized counterparts using MUSCLE (default settings) (27). Maximum-likelihood phylogenies were inferred with IQ-TREE (default model selection; 1,000 bootstrap replicates), and branch support is reported as bootstrap percentages (28). Trees were visualized and annotated in iTOL (29).

### Gene cloning methods for heterologous expression

Candidate genes (Table S1) were PCR-amplified from first-strand cDNA synthesized from total RNA of young leaves of *Tabernanthe iboga* and *Catharanthus roseus*. RNA from *T. iboga* and *C. roseus* was extracted as previously described (12). cDNA was prepared using SuperScript IV VILO Master Mix (Thermo Fisher Scientific) per the manufacturer’s instructions. PCRs used Q5 Hot Start High-Fidelity 2X Master Mix (New England Biolabs) with primers bearing overhangs compatible with In-Fusion HD Cloning (Takara Bio), and the primer sequences are listed in Table S2. Amplicons were purified using the DNA Clean and Concentrator-5 kit (Zymo Research) and cloned with In-Fusion HD Cloning kit (Takara Bio) into pre-digested vectors as follows: plant expression, 3Ω1 (BsaI-HF, spectinomycin resistance) for *Nicotiana benthamiana* and *C. roseus*; bacterial expression, pOPINF (HindIII-HF/KpnI-HF, carbenicillin), Addgene #26042) for N-terminal Hisx6 tags and pET28a(+) (NcoI-HF/XhoI-HF, kanamycin), Novagen #69864) for C-terminal Hisx6 tags; yeast expression, pESC-HIS (SalI-HF/BamHI-HF, carbenicillin), Agilent technologies #217451). Assembled plasmids were transformed into *E. coli* TOP 10 cells (Thermo Fisher Scientific) by heat shock (42 °C) and plated on Luria-Bertani (LB) agar containing the appropriate antibiotics. After overnight incubation at 37 °C, and single colonies were cultured in selective LB medium (5 ml) at 37 °C, 200 rpm. Plasmid DNA was extracted using the Wizard Plus SV Minipreps DNA Purification System (Promega) and verified by Sanger sequencing using the primers listed in Table S2. Precondylocarpine acetate synthase (PAS) from *C. roseus* and *T. iboga* with a C-terminal Hisx6 tag for *N. benthamiana* expression was cloned into 3Ω1 as previously described (11).

Multigene constructs for plant transient expression were designed and assembled using the GoldenBraid 2.0 toolkit (30, 31). In silico assemblies were built in Geneious Prime version. 2023.1.2. Genes were screened for internal BsaI and BsmbI/Esp3I sites. Sequences containing sites were domesticated by PCR using primers generated from the GoldenBraid Domesticator web tool (Table S3). Domesticated ORFs were PCR-amplified with Level-1 overhangs primers containing overhangs (Table S3) and assembled with the *Solanum lycopersicum* Ubiquitin10 promoter and terminator (SlUbq10p/t; Table S1) into 3α1/3α2 (kanamycin) using BsaI-HF and T4 DNA ligase (New England Biolabs) by Golden Gate PCR cycling (37 °C/16 °C, 5 min each, 50 cycles; 65 °C, 10 min). Level 1 transcriptional units were sequence-verified by Sanger sequencing using gene-specific primers (Table S3). Level 2 assemblies were generated using Level 1 units in 3Ω1/3Ω2 (spectinomycin) using Esp3I-HF and T4 DNA ligase (New England Bioscience) and verified by Sanger sequencing using primers listed in Table S3. Final Level 3 assemblies were built using Level 2 constructs into the 3α1 using BsaI-HF and validated by whole-plasmid sequencing (Plasmidsaurus). Golden Gate cloning products were transformed into *E. coli* TOP10 cells by heat shock (42 °C), plated on selective LB agar, and blue-white screening was used to identify positive transformants, and plasmids were purified as described above.

For subcellular localization, biosynthetic genes were PCR-amplified with a C-terminal GSGSS-linker (primers in Table S4) and assembled as C-terminal fusions to eYFP (GoldenBraid part GB0024) under SlUbq10p/t in 3α1 by Golden Gate PCR-cycling with BsaI-HF and T4 DNA ligase using the conditions described above.

Assembled constructs were transformed into *E. coli* TOP10 cells by heat shock (42 °C) and selected on LB agar (kanamycin, 50 μg/ml), screened on blue-white selection, and verified by Sanger sequencing using primers listed in Table S4. Plant organelle marker constructs fused to red fluorescent protein (mCherry) for subcellular localization were obtained from Addgene (32).

Transformation into expression hosts was conducted as follows. Sequence verified plant transient expression constructs (3Ω1 and 3α1) were electroporated into *Agrobacterium tumefaciens* GV3101 cells (Goldbio; Gentamycin, Rifampicin). The transformed cells were recovered in 1=ml of LB medium, incubated at 28 °C and 200 rpm for 4=h, then plated on selective LB agar plates and incubated at 28 °C for 48=h.

Single colonies were confirmed by colony PCR and grown overnight in selective LB medium at 28 °C and 200=rpm. Overnight cultures were used to prepare 25% glycerol stocks for storage at-80 °C till further use. Sequence-verified bacterial expression constructs (pOPINF and pET28a(+)) were transformed into *E. coli* BL21 (DE3) cells by heat shock (42 °C), outgrown in 1 ml LB at 37 °C, 200 rpm for 1 hour, and plated on selective LB agar and incubated at 37 °C for 24 h. Single colonies were PCR-verified, grown in selective LB medium (37 °C, 200=rpm, overnight), and stored as 25% glycerol stocks at-80 °C till further use. Sequence-verified yeast expression constructs (pESC-HIS) were transformed into the *Saccharomyces cerevisiae*

WAT11 strain (ade2; contains the *Arabidopsis thaliana* cytochrome P450 reductase I gene, ATR1) using the Frozen-EZ Yeast Transformation II Kit (Zymo Research). Transformants were selected on agar containing SD-His medium (6.7 g/l yeast nitrogen base without amino acids, 2 g/l drop-out mix without histidine, 74 mg/l adenine hemisulfate) supplemented with 2% glucose (w/v) and grown at 30 °C for 48 h. Positive transformants were confirmed by colony PCR, grown in SD-His + 2% glucose medium (10 ml, 30 °C, 220 rpm, 48 h). These cultures were then used to prepare 25% glycerol stocks, which were stored at-80 °C until further use.

### qPCR analysis

Total RNA was extracted from *Nicotiana benthamiana* leaf tissues using the RNeasy Mini Kit (Qiagen) according to the manufacturer’s protocol. Genomic DNA was digested with DNase I (Zymo Research), and RNA was further purified using the RNA Clean & Concentrator-25 (Zymo Research) according to the manufacturer’s protocol. First-strand cDNA synthesis was performed with 1000 ng of RNA using the SuperScript-IV VILO master mix (Thermo Fisher Scientific) following the manufacturer’s instructions.

Quantitative real-time PCR (qPCR) was conducted on a QuantStudio 1 Real-Time PCR System (Applied Biosystems) using Fast SYBR™ Green Master Mix (Applied Biosystems). Reactions were performed in fast mode, with cDNA diluted 1:10 before amplification. Primers used for qPCR amplification are presented in Table S5. The *N.□benthamiana* housekeeping gene Elongation factor 1 alpha (EF1α, 8965.1) was used as the internal reference for normalization (33). Each treatment included six biological replicates (*n* = 6), and all reactions were performed in technical triplicate. Relative transcript levels were calculated using the 2^−ΔΔCT^ method, and data analysis was performed using the QuantStudio Analysis Software. Statistical analysis and data visualization were performed using GraphPad Prism 10 (version 10.4.2).

### Agrobacterium-mediated transient expression in Nicotiana benthamiana and Catharanthus roseus

Pathway reconstitution, in vivo assays, heterologous protein expression, and subcellular enzyme localization studies were performed via *Agrobacterium tumefaciens* (GV3101) mediated transient expression in *N. benthamiana* leaves and *C. roseus* flower petals as previously described (11, 12). *Agrobacterium* strains harboring the gene constructs of interest were grown overnight at 28 °C and 220rpm. Cells were harvested by centrifugation at 4000 ×=g for 10=min, the supernatant was discarded, and the pellet was resuspended in infiltration buffer (10 mM MES, pH 5.6, 10 mM magnesium chloride, 200 μM acetosyringone). Following a second centrifugation and discard of supernatant, cells were resuspended in 10 ml of fresh infiltration buffer and incubated in the dark at room temperature for 2 h with gentle rocking. For co-infiltration of multiple constructs, strains were combined in equal volumes at an OD_600_ of 0.4 per strain. For multigene constructs, cultures were adjusted to a final OD_600_ of 0.8. For subcellular localization experiments, *Agrobacterium* strains carrying P19-TBSV, organelle marker genes, and mCherry-fusion constructs were adjusted to a final OD_600_ of 0.3 (0.1 per strain). Infiltration of the *Agrobacterium* inoculum into *N. benthamiana* leaves were performed on the abaxial side of 3-week-old plants using a 1 ml needleless syringe. For *C. roseus* flower petal infiltration, a single puncture was made on the petal with a sterile needle, and the *Agrobacterium* suspension was infiltrated at the site. Leaf discs from *N. benthamiana* were harvested 3 days post-infiltration for pathway reconstitution and in vivo assays, while leaf and petal discs were collected 2 days post-infiltration for localization experiments. For protein expression and purification of precondylocarpine acetate synthase (PAS), C-terminal 6×His-tagged TiPAS1–3 and CrPAS were transiently expressed in *N. benthamiana* leaves as described previously (11). All experiments included at least three biological replicates, using different leaves from three independent plants to minimize batch effects. Appropriate negative and experimental controls: wild-type, P19-TBSV, and GFP were included, with and without substrate addition.

### Heterologous protein expression in *Escherichia coli* and His tagged protein purification

Recombinant proteins were expressed in *Escherichia coli* BL21 (DE3) cells. Starter cultures (10 ml) were prepared from glycerol stocks (see section cloning methods) in LB medium supplemented with 100 µg/ml carbenicillin for pOPINF-based constructs and 50 µg/ml kanamycin for pET28a(+)-based constructs and incubated at 37 °C and 200 rpm for 16 h. The starter cultures (2 ml) were used to inoculate 100 ml of auto-induction medium (Formedium, cat no. AIM2YT0205) supplemented with the appropriate antibiotic, and the cultures were grown to an OD_600_ of 0.8 at 37°C, 200 rpm, then incubated at 18 °C, 200 rpm for overnight protein expression. Cells were harvested by centrifugation at 3,000 g, 4 °C, for 10 min, and resuspended in 10 ml of pre-chilled lysis buffer (buffer A: 50 mM Tris-HCl, pH 7.4, 50 mM glycine, 500 mM NaCl, 5% glycerol, 20 mM imidazole) supplemented with 10 mg/ml lysozyme and EDTA-free protease inhibitor (Roche Diagnostics). All subsequent steps were performed on ice or at 4 °C. Cells were lysed by sonication (5 min total, 2 sec on/ 3 sec off cycles) and the lysate was clarified by centrifugation at 35,000 × g, 4 °C, for 20 min, followed by filtration through a 0.45 µm glass filter. His_6_-tagged proteins were purified using an ÄKTA Pure 25 fast protein liquid chromatography (FPLC) system (Cytiva) on a 1 ml HisTrap HP column (Cytiva) pre-equilibrated with buffer A. The lysate was loaded at 0.5 ml/min, and bound proteins were eluted in 0.5 ml fractions using buffer B (50 mM Tris-HCl, pH 7.4, 50 mM glycine, 500 mM NaCl, 5% glycerol, 500 mM imidazole). Elution fractions were analyzed by SDS-PAGE with coomassie blue staining. Fractions containing the protein of interest were pooled and concentrated using Amicon Ultra centrifugal filters (10 kDa MWCO, Millipore) by centrifugation according to the manufacturer’s instructions. The protein was dialyzed in the same filter unit using buffer C (20 mM HEPES, pH 7.4, 150 mM NaCl), then concentrated to a final volume of 0.2 ml. Protein concentrations were determined using a NanoPhotometer (Implen) by measuring absorbance at 280 nm and applying theoretical extinction coefficients. Proteins were aliquoted, snap-frozen in liquid nitrogen, and stored at −80 °C until use in enzymatic assays. For large-scale protein purification, 1 l *E. coli* cultures were used, and the proteins were expressed and purified under the same conditions as described above using a 5 ml HisTrap HP column (Cytiva). C-terminal His_6_-tagged TiPAS1-3 and CrPAS was transiently expressed in *Nicotiana benthamiana* and purified using the same buffers on a 5 ml HisTrap HP column (Cytiva) as previously described (11).

### Yeast protein expression and microsome preparation

*S. cerevisiae* WAT11 cells transformed with a pESC-HIS expression construct encoding P450 enzyme (TiGO or TiSBE) or with an empty cassette (EV) control, were streaked from glycerol stocks onto SD-His medium (6.7 g/l yeast nitrogen base without amino acids, 2 g/l drop-out mix without histidine, 74 mg/l adenine hemisulfate) containing agar (20 g/l) plates supplemented with 2% glucose (w/v) and incubated at 30 °C for 48 h. Individual colonies were inoculated into 10 ml of SD-His + 2% glucose (w/v) medium and cultured overnight at 30 °C, 200 rpm. The following day, cultures were diluted to an OD_600_ of 1.0 into 100 ml of SD-His + 2% glucose (w/v) medium and incubated for 28-34 h at 30 °C, 200 rpm. Cells were harvested by centrifugation at 4000 × g for 5 min at room temperature, resuspended in 100 ml of SD-His medium containing 1.8% galactose (w/v) and 0.2% glucose (w/v), and incubated for a further 18-24 h at 30 °C, 200 rpm for induction of protein expression. All culturing and handling steps were performed under aseptic conditions. For microsome preparation, cells were harvested by centrifugation at 4000 × g, 4 °C, 10 min, resuspended in 10 ml (2 ml/g wet weight) of TEK buffer (50 mM Tris-HCl, 1 mM EDTA, 100 mM KCl, pH 7.4), and incubated for 5 min at room temperature. Cells were pelleted again, and resuspended in 2 ml of ice-cold TES buffer (50 mM Tris-HCl, 1 mM EDTA, 600 mM sorbitol, 10 g/l BSA, pH 7.4), and lyzed with equal volume of 0.5 mm glass beads using a Bead Genie homogenizer (Scientific Industries) for eight cycles (1 min on/1 min off at 5000 rpm) at 4 °C. The lysate was supplemented with 5 ml of ice-cold TES buffer, mixed thoroughly, and the supernatant was collected into pre-chilled tubes. Beads were washed three times with TES buffer, and all supernatants were pooled and centrifuged at 8000 × g, 4 °C, for 10 min to remove cell debris. The clarified supernatant was subjected to ultracentrifugation at 100,000 × g, 4 °C, for 90 min to pellet microsomal membranes. The resulting translucent pellet was washed once with 1 ml ice-cold TEG buffer (50 mM Tris-HCl, 1 mM EDTA, 20% glycerol, pH 7.4) and then resuspended with 1 ml of TEG buffer using a Dounce homogeniser. Microsomal protein preparations were aliquoted and stored at –80 °C until use in enzyme assays.

### In vivo enzyme assays in *Nicotiana benthamiana*

In vivo enzymatic assays were performed using *N. benthamiana* leaf disks following *Agrobacterium*-mediated transient gene expression, as previously described (12). Three days post-infiltration (see section *Agrobacterium*-mediated transient expression in *Nicotiana benthamiana*), three 10 mm leaf disks were excised from each infiltrated leaf using a cork-borer (Ø 1 cm) and transferred into individual wells of a 48-well plate (10 mm diameter wells). Each well contained 250 µl of 50 mM HEPES buffer (pH 7.5), and an individual leaf disk was placed abaxial or adaxial side down and immediately submerged in the buffer.

Substrates, 19*E/Z*-geissoschizine or stemmadenine, were added to the wells at a final concentration of 50 µM. For strictosidine, the substrate was directly infiltrated into the transiently expressing *N. benthamiana* leaves by first piercing the abaxial side of the tissue with a needle and then infiltrating 50 µl of 200 µM strictosidine (aq) solution using a needleless syringe. Substrate infiltrated plants were incubated under the same growth conditions for 24 h prior to harvesting for metabolite extraction. Wild-type and/or P19-TBSV infiltrated plants incubated with the same substrates were included as controls in all experiments. Plates were sealed with lids and parafilm to prevent buffer evaporation and incubated in a growth chamber at 22 °C, 60% relative humidity, under a 16 h light/8 h dark photoperiod for 14 h (overnight). After incubation, the leaf disks were briefly blotted on tissue paper to remove excess buffer, transferred to 2 ml safe-lock tube containing clean, two 3 mm tungsten carbide beads (Qiagen), and flash-frozen in liquid nitrogen. The leaf disks were ground using a TissueLyser II homogenizer (Qiagen) at 25 Hz for 1 min. Metabolites were extracted by adding 300 µl of 70% methanol (aq) supplemented with 0.1% formic acid to the powdered tissue, vortexing, and sonicating for 10 min at room temperature. For strictosidine infiltrated plants, three disks were harvested 24 h post-infiltration from the treated area and subjected to the same metabolite extraction procedure. Extracts were clarified by centrifugation at 20,000 × g for 10 min, and the supernatant was filtered through 0.22 µm PTFE syringe filters before LC-MS analysis. All in vitro assays were performed with a minimum of three biological replicates conducted on independent occasions.

### In vitro enzyme assays

All analytical-scale in vitro reactions using purified recombinant proteins or microsomal preparations were carried out in 50=mM HEPES buffer (pH=7.5) in a final volume of 100=μl. Reactions contained of 2 μM purified enzyme prepared as 50 μM stock solutions in protein buffer C and/or 50=μg of total microsomal protein (0.50=μg/μl microsomal protein) with 25 μM substrate. The substrates, strictosidine, 19*E/Z*-geissoschizine, stemmadenine were prepared at 2 mM stock concentrations in methanol. Where required, cofactors prepared as 10 mM stock solution in water were added to a final concentration of 1 mM for NADPH (Sigma Aldrich, cat. no 10107824001), 0.5 mM for acetyl Co-A (Sigma Aldrich, cat. no. A2181), and 0.5 mM for malonyl Co-A (Sigma Aldrich, cat. no M4263). Negative control reactions consisted of boiled protein or empty vector (EV) microsomal fractions. All reactions were carried out in biological triplicates from different enzyme preparations. For assays involving deuterium-labeled cofactor, NADPH was replaced with NADPD generated in situ as previously described (11). In 100 μl reactions, 0.5 mM NADP (Sigma-Aldrich, cat. no. 10128031001), 5 mM isopropanol-*d*_8_ (Sigma-Aldrich, cat. no. 175897), and 1% (v/v) alcohol dehydrogenase (Sigma-Aldrich, cat. no. 49641) were added to regenerate Pro-(*R*)-NADPD. Reactions were initiated by the addition of substrate and incubated at 30 °C with shaking at 600 rpm for 60 min in a thermomixer (Eppendorf). Reactions were quenched by the addition of 100 μl of 70% methanol (aq) supplemented with 0.1% formic acid, centrifuged at 20000=× g for 10 min at room temperature, and the supernatant was filtered through a 0.22 μm PTFE syringe filter into LC-MS vials for analysis.

### Large-scale enzyme assays for product isolation

Enzymatic reactions were scaled to a final volume of 10 ml to generate 19*E*-16(*R/S*)-isositsirikine and their deuterium-labeled analogs (19*E*-16(*R/S*)-isositsirikine-*d*) from 19*E*-geissoschizine. Reactions were assembled in 50 mM HEPES buffer (pH 7.5) containing 2 mg of 19*E*-geissoschizine, a NADPH generation system (1.0=mM NADP^+^; 2.0=U/ml glucose-6-phosphate dehydrogenase (G6PDH); 3.0 mM glucose-6-phosphate (G6P); 0.2 mM NADPH), and TiRedox2 enzyme at a final concentration of 100 μM. Reactions were incubated at 30 °C for 14 h. For the preparation of deuterium-labeled analogs (19*E*-16(*R/S*)-isositsirikine-*d*), the NADPH generation system was substituted with a deuterated (NADPD) system containing 1.0 mM NADP^+^, 5 mM isopropanol-*d*_8_ (Sigma-Aldrich, 175897), 1% (v/v) alcohol dehydrogenase (Sigma-Aldrich, 49641). All reaction components and conditions were kept the same. Following incubation, the reaction mixtures were passed through preconditioned CHROMABOND HR-X solid-phase extraction (SPE) cartridges (85 µm, 3 ml/60 mg; Macherey-Nagel). The SPE cartridges were first conditioned with three column volumes each of 100% methanol and 10% methanol (aq). The 10 ml reaction mixtures were loaded onto the SPE cartridges, washed with three column volumes of 20% methanol (aq), and dried under vacuum. Elution was performed with 3 ml of methanol, and the eluates were evaporated to dryness. The dried reaction products were reconstituted with 5 ml of 50% methanol (aq) and filtered through 0.22 µm PTFE membranes before preparative HPLC purification.

### Extraction of stemmadenine and condylocarpine from *Nicotiana benthamiana*

A total of 50 *N. benthamiana* plants (two leaves per plant were infiltrated) were infiltrated with *Agrobacterium* harbouring the multigene construct pMGC01, containing the stemmadenine biosynthetic genes. Infiltration was performed as described in the transient overexpression protocol at an OD_600_ of 1.0. Four days post-infiltration, 1 ml of 0.5 mM 19*E*-geissoschizine was infiltrated into the same two leaves per plant. Leaves were harvested 14-16 h after the substrate was infiltrated. The stems were removed, and the leaf tissue was flash-frozen in liquid nitrogen, then lyophilized in a freeze dryer (Labconco). The dehydrated leaf material was ground into a fine powder and extracted twice with 500 ml of 70% methanol (aq) at room temperature for 3 h under constant stirring. The extract was filtered under vacuum using MN 615 filter paper (Macherey-Nagel), and the filtrate was evaporated to dryness using a rotary evaporator. The dried extract was reconstituted in 20 ml of 50% methanol (aq), filtered through a 0.22 µm PTFE membrane, and prepared for purification by preparative HPLC.

### Purification of compounds by preparative HPLC

Target compounds were isolated and purified using an Agilent 1260 Infinity II Preparative High-Performance Liquid Chromatography (HPLC) system equipped with a 2 ml sample injection loop, a multiple-wavelength detector, and a fraction collector (Agilent Technologies). Chromatographic separation was carried out at room temperature (25 °C) using a Kinetex XB-C18 (250 x 10 mm, 5 µm; 100 Å, Phenomenex) column. For isolation of stemmadenine and condylocarpine from *Nicotiana benthamiana*, the mobile phases were water with 0.1% formic acid (A) and acetonitrile (B). The elution gradient was 10–30% B over 15=min, 90% B for 7 min, followed by re-equilibration at 10% B for 5=min with a flow rate of 7.0 ml/min. Manual injections of 1 ml were performed per run, and the separation for fractionation was monitored at wavelengths of 222, 254, and 328 nm. For the isolation of enzymatic products 19*E*-16(*R/S*)-isositsirikine and their deuterium-labeled analogs (19*E*-16(*R/S*)-isositsirikine-*d*), the mobile phases were 5 mM ammonium acetate in water (A) and 5 mM ammonium acetate in methanol (B). Separation was performed using a gradient of 15–25% B over 15 min, 90% B for 7 min, and re-equilibration at 10% B for 5 min with a flow rate of 7.0 ml/min. Manual injections of 0.25 ml were made for each run, and the detection for separation and fractionation was monitored at wavelengths of 220 and 254 nm. Collected fractions were analyzed by LC-MS, and those containing the desired products were pooled, dried using a rotary evaporator, and analyzed by NMR.

### LC-MS analysis and metabolomics

LC-MS analysis was performed using a Thermo Scientific UltiMate 3000 ultra-high performance liquid chromatography (UHPLC) system coupled to a Bruker Impact II ultra-high-resolution quadrupole-time-of-flight (UHR-Q-ToF) mass spectrometer (Bruker Daltonics). Two liquid chromatography (LC) methods were used. All MIAs except for 19*E/Z*-geissoschizine were analyzed using a Kinetex XB-C18 (Phenomenex) column as described below, while the separation of 19*E/Z*-geissoschizine was achieved using a BEH C18 (Waters) column with modified mobile phases and gradients adapted from previously published methods (6, 13).

For general MIA analysis, metabolites were separated by reversed-phase liquid chromatography using a Kinetex XB-C18 (100 x 2.1 mm, 2.6 µm; 100 Å, Phenomenex) column at 40 °C. Mobile phase A consisted of water with 0.1% formic acid, and mobile phase B was acetonitrile. The flow rate was 0.6 ml/min, and 2 µl of each sample was injected. The chromatographic gradient was programmed as follows: 10% B for 1 min, then a linear increase to 30% B over 5 min, followed by 90% B for 1.5 min, and re-equilibration at 10% B for 2.5 min (total run time: 10 min). Authentic standards (20 µM solutions in methanol) were analyzed under the same conditions. Mass spectrometry (MS) acquisition was performed in positive electrospray ionization (+ESI) mode with the following settings: capillary voltage, 3500 V; end plate offset, 500 V; nebulizer pressure, 2.5 bar; and drying gas nitrogen (N_2_), 11 l/min at 250 °C. MS data were acquired in the *m/z* range 80-1000 at 12 Hz. Tandem mass spectra (MS^2^) were acquired in a data-dependent mode, triggered at an absolute intensity threshold of 400 counts, with a 0.5 sec cycle time and collision energy stepping from 20 to 50 eV. An active exclusion window of 0.2 min was applied. Before each run, the instrument was externally calibrated with a sodium formate-isopropanol solution at a rate of 0.18 ml/h using a syringe pump. To prevent salt contamination, the first minute of each chromatographic run was diverted to waste. For quantification of stemmadenine, the MS was set to full-scan mode to maximize the number of data points across the chromatographic peak for analyte quantitation. For this purpose, the spectral rate was set to 2 Hz, and a mass range 80-1000 *m/z* was selected.

For the separation of 19*E/Z*-geissoschizine, a modified LC program from previously reported methods was applied (6). Mobile phase (A) was water with 0.05% ammonium hydroxide (pH 10.0) (Merk Millipore, 5.33003.0050), mobile phase (B) was acetonitrile. The flow rate was set to 0.4 ml/min, and 2 µl of the sample was injected. The gradient was set to 10% B for 1 min, then linearly increased to 90% B over 8 min, followed by a 1.5 min wash at 90% B and a 2.5 min re-equilibration at 10% B (total run time: 14 min). Separation was performed on an Acquity UPLC BEH-C18 column (2.1 x 50 mm; 1.7 µm; 130 Å, Waters) maintained at 50 °C. Authentic standards (20 µM solutions in methanol) were analyzed under the same conditions. The MS acquisition parameters remained unchanged.

LC-MS data were visualized using Bruker Compass Data Analysis (version 5.3). For relative quantification and metabolomics, data were analyzed using default pipelines embedded in Bruker Compass MetaboScape 2024b (version 7.0.1). The non-targeted metabolomics workflow was used to extract features based on accurate mass (±5 ppm) and retention time, with automated peak integration of extracted ion chromatograms. Where appropriate, the internal standard (ajmaline) was used to calculate relative intensities and peak areas of analytes for comparisons across treatments. MS^2^ spectral similarity was calculated using tidyMass (34). A standard curve for stemmadenine with 9 calibrants ranging from 0.5 µM to 10 µM was used to quantify stemmadenine produced in *N. benthamiana*. Statistical comparisons were performed using GraphPad Prism 10 (version 10.4.2).

### NMR analysis

NMR spectra were acquired on a Bruker Avance III HD 700 MHz spectrometer (Bruker Biospin GmbH, Rheinstetten, Germany) equipped with a TCI cryoprobe. Standard pulse sequences were used as implemented in Bruker TopSpin (version 3.6.1). Chemical shifts were referenced to the residual solvent signals of MeOH-*d*_3_ (δ_H_ 3.31/δ_C_ 49.0), CDCl_3_ (δ_H_ 7.24/δ_C_ 77.23). All spectra were recorded at 298 K. Data processing and visualization were performed using Bruker TopSpin (version 3.6.1).

### Confocal laser scanning microscopy

Micrographs of freshly punched *Nicotiana benthamiana* leaf discs and *Catharanthus roseus* petal discs were acquired using a Zeiss cLSM 880 Axio Imager 2 confocal laser scanning microscope (Zeiss, Oberkochen, Germany) equipped with a C-Apochromat 40x/1.20 water immersion objective. The sample disks were water-mounted in custom-designed 3D-printed object slides with circular wells, 400 µm deep for leaf disks and 200 µm deep for petal disks and covered with standard 170 µm-thick cover glasses. Imaging was performed using two frame-sequential tracks, each comprising a single detection channel. In the first track, eYFP was excited with a 514 nm Argon laser (3–5% transmission) using a main beam splitter (MBS) with a 458/514 nm bandpass filter. Emission was detected between 517–568 nm (detector gain 550) with a pinhole set to 1 Airy Unit. This track also included a transmitted light T-PMT channel (gain 250) in brightfield mode.

The second track was dedicated to mCherry detection. Excitation was performed with a 543=nm Argon laser (15–25% transmission) using an MBS 458/543 splitter. Emission was detected in the range of 590–735 nm (detector gain 725). Images were acquired primarily using unidirectional scanning with an averaging of 8 and a pixel dwell time of approximately 1 µs. The resolution and zoom settings were optimized for each sample to ensure high-quality imaging.

### Protein structure prediction and molecular docking

Protein structure predictions for *Tabernanthe iboga* geissoschizine synthase (TiGS) and Redox2 (TiRedox2) were generated using AlphaFold v3 (35). Co-factor NADPH was docked into the predicted structures of TiGS and TiRedox2 using AlphaFold3-integrated ligand modelling tools. Molecular docking of the TiRedox2 substrates 19*E/Z*-geissoschizine (PubChem CIDs: 10948159 and 10893550) was performed using AutoDock Vina (version 1.1.2) (36). For each substrate, the docking pose with the lowest predicted binding energy was selected for structural interpretation. Protein structures and docking conformations were visualized and analyzed using PyMol (version 2.5.5; Schrödinger, LLC).

## Data availability

Genes characterized in this study are deposited in NCBI GenBank with the following accession numbers: TiSGD (xxxxxxxx), TiGS (xxxxxxxx), TiGO (xxxxxxxx), TiSBE (xxxxxxxx), TiRedox1 (xxxxxxxx), TiRedox2 (xxxxxxxx), TiSAT (xxxxxxxx).

## Supporting Information

This article contains supporting information.

## Acknowledgments

We thank Dr. Samuel Carr, Dr. Chloe Langley, and Dr. Moonyoung Kang for valuable discussions. We acknowledge Dr. Prashant Sonowanne for providing the modified 3Ω1 vector and advice on plant binary vector constructions for *N. benthamiana* expression. We thank the greenhouse team for the horticulture services provided for *T. iboga* and *N. benthamiana*, with special thanks to Eva Rothe and Franz Kaltofen. This work was supported by the Max Planck Society.

## Author contributions

S.O.C., M.O.K., and L.C. Conceptualization; M.O.K., Y.N., V.G., B.H., and G.K. Data curation; M.O.K., S.O.C., Y.N., V.G., B.H., R.A., and G.K. Formal analysis; S.O.C. Funding acquisition; M.O.K., Y.N., M.S., R.K., V.G., S.H., M.K., B.H., R.A., and G.K. Investigation; M.O.K., Y.K., V.G., S.H., B.H., R.A., and G.K. Methodology; S.O.C., and M.O.K. Project administration; S.O.C. Resources; S.O.C., M.O.K., L.C. Supervision; M.O.K., S.O.C., Y.N., V.G., S.H., and M.K. Validation; M.O.K., V.G., and Y.N. Visualization; M.O.K., and S.O.C. Writing - original draft; S.O.C., M.O.K., Y.N., S.H., and M.K. Writing - review & editing.

## Funding and additional information

We gratefully acknowledge the Max Planck Society.

## Abbreviation and nomenclature

MIA: monoterpene indole alkaloid
Ti: *Tabernanthe iboga*
Cr: *Catharanthus roseus*
SGD: strictosidine β-D-glucosidase
GS: geissoschizine synthase
GO: geissoschizine oxidase
SBE: sarpagan bridge enzyme
SAT: stemmadenine acetyltransferase
PAS: precondylocarpine acetate synthase
MDR: medium-chain dehydrogenase/reductase
AKR: aldo-keto reductase
BBL: berberine bridge-like
pMGC: plasmid multigene construct.

