## Supporting Information for "Conserved early steps of stemmadenine biosynthesis"

**Contents**


- Supporting Table S1-S5 pages 2-7
- Supporting NMR Table S1-S6 pages 8-10
- Supporting Figure S1-S39 pages 11-51
- References page 52

### Supporting Tables

**Table S1**. Nucleotide sequences of biosynthetic genes and transcriptional elements described in this study.

| Gene Name | Nucleotide sequence |
|---|---|
| P19<br>(GoldenBraid 2.0 part ID GB0038) | ATGGAACGAGCTATACAAGGAAACGACGCTAGGGAACAAGCTAACAGTGAACGTTGGGATGGAGGATCAGGAGGTACCACTTCT<br>CCCTTCAAACTTCCTGACGAAAGTCCGAGTTGGACTGAGTGGCGGCTACATAACGATGAGACTAATTCGAATCAAGATAATCCC<br>CTTGGTTTCAAGGAAAGCTGGGGTTTCGGGAAAGTTGTATTTAAGAGATATCTCAGATACGACAGGACGGAAGCTTCACTGCAC<br>AGAGTCCTTGGATCTTGGACGGGAGATTCGGTTAACTATGCAGCATCTCGATTTTTCGGTTTCGACCAGATCGGATGTACCTAT<br>AGTATTCGGTTTCGAGGAGTTAGTATCACCGTTTCTGGAGGCTCTCGAACTCTTCAGCATCTCTGTGAGATGGCAATTCGGTCT<br>AAGCAAGAACTGCTACAGCTTGCCCCAATCGAAGTGGAAAGTAATGTATCAAGAGGATGCCCTGAAGGTACTGAAACCTTCGAA<br>AAAGAAAGCGAGTGA |
| GFP<br>(Green fluorescent protein) | ATGGTGAGCAAGGGCGAGGAGCTGTTCACCGGGGTGGTGCCCATCCTGGTCGAGCTGGACGGCGACGTAAACGGCCACAAGTTC<br>AGCGTGTCCGGCGAGGGCGAGGGCGACGCCACCTACGGCAAGCTGACCCTGAAGTTCATCTGCACCACCGGCAAGCTGCCCGTG<br>CCCTGGCCCACCCTCGTGACCACCTTCAGCTACGGCGTGCAGTGCTTCAGCCGCTACCCCGACCACATGAAGCAGCACGACTTC<br>TTCAAGTCCGCCATGCCCGAAGGCTACGTCCAGGAGCGCACCATCTTCTTCAAGGACGACGGCAACTACAAGACCCGCGCCGAG<br>GTGAAGTTCGAGGGCGACACCCTGGTGAACCGCATCGAGCTGAAGGGCATCGACTTCAAGGAGGACGGCAACATCCTGGGGCAC<br>AAGCTGGAGTACAACTACAACAGCCACAACGTCTATATCATGGCCGACAAGCAGAAGAACGGCATCAAGGTGAACTTCAAGATC<br>CGCCACAACATCGAGGACGGCAGCGTGCAGCTCGCCGACCACTACCAGCAGAACACCCCCATCGGCGACGGCCCCGTGCTGCTG<br>CCCGACAACCACTACCTGAGCACCCAGTCCGCCCTGAGCAAAGACCCCAACGAGAAGCGCGATCACATGGTCCTGCTGGAGTTC<br>GTGACCGCCGCCGGGATCACTCACGGCATGGACGAGCTGTACAAGTGA |
| SlUbq10p<br>(*Solanum lycopersicum* Ubiquitin 10 promoter) | GGAGGTCAACTACCCCAATTTAAATTTTATTTGATTAAGATATTTTTATGGACCTACTTTATAATTAAAAATATTTTCTATTTG<br>AAAAGGAAGGACAAAAATCATACAATTTTGGTCCAACTACTCCTCTCTTTTTTTTTTTGGCTTTATAAAAAAGGAAAGTGATTA<br>GTAATAAATAATTAAATAATGAAAAAAGGAGGAAATAAAATTTTCGAATTAAAATGTAAAAGAGAAAAAGGAGAGGGAGTAATC<br>ATTGTTTAACTTTATCTAAAGTACCCCAATTCGATTTTACATGTATATCAAATTATACAAATATTTTATTAAAATATAGATATT<br>GAATAATTTTATTATTCTTGAACATGTAAATAAAAATTATCTATTATTTCAATTTTTATATAAACTATTATTTGAAATCTCAAT<br>TATGATTTTTTAATATCACTTTCTATCCATGATAATTTCAGCTTAAAAAGTTTTGTCAATAATTACATTAATTTTGTTGATGAG<br>GATGACAAGATTTCGGTCATCAATTACATATACACAAATTGAAATAGTAAGCAACTTGATTTTTTTTCTCATAATGATAATGAC<br>AAAGACACGAAAAGACAATTCAATATTCACATTGATTTATTTTTATATGATAATAATTACAATAATAATATTCTTATAAAGAAA<br>GAGATCAATTTTGACTGATCCAAAAATTTATTTATTTTTACTATACCAACGTCACTAATTATATCTAATAATGTAAAACAATTC<br>AATCTTACTTAAATATTAATTTGAAATAAACTATTTTTATAACGAAATTACTAAATTTATCCAATAACAAAAAGGTCTTAAGAA<br>GACATAAATTCTTTTTTTGTAATGCTCAAATAAATTTGAGTAAAAAAGAATGAAATTGAGTGATTTTTTTTTAATCATAAGAAA<br>ATAAATAATTAATTTCAATATAATAAAACAGTAATATAATTTCATAAATGGAATTCAATACTTACCTCTTAGATATAAAAAATA<br>AATATAAAAATAAAGTGTTTCTAATAAACCCGCAATTTAAATAAAATATTTAATATTTTCAATCAAATTTAAATAATTATATTA<br>AAATATCGTAGAAAAAGAGCAATATATAATACAAGAAAGAAGATTTAAGTACAATTATCAACTATTATTATACTCTAATTTTGT<br>TATATTTAATTTCTTACGGTTAAGGTCATGTTCACGATAAACTCAAAATACGCTGTATGAGGACATATTTTAAATTTTAACCAA<br>TAATAAAACTAAGTTATTTTTAGTATATTTTTTTGTTTAACGTGACTTAATTTTTCTTTTCTAGAGGAGCGTGTAAGTGTCAAC<br>CTCATTCTCCTAATTTTCCCAACCACATAAAAAAAAAAATAAAGGTAGCTTTTGCGTGTTGATTTGGTACACTACACGTCATTAT<br>TACACGTGTTTTCGTATGATTGGTTAATCCATGAGGCGGTTTCCTCTAGAGTCGGCCATACCATCTATAAAATAAAGCTTTCTG<br>CAGCTCATTTTTTCATCTTCTATCTGATTTCTATTATAATTTCTCTGAATTGCCTTCAAATTTCTCTTTCAAGGTTAGAATTTT<br>TCTCTATTTTTTGGTTTTTGTTTGTTTAGATTCTGAGTTTAGTTAATCAGGTGCTGTTAAAGCCCTAAATTTTGAGTTTTTTTC<br>GGTTGTTTTGATGGAAAATACCTAACAATTGAGTTTTTTCATGTTGTTTTGTCGGAGAATGCCTACAATTGGAGTTCCTTTCGT<br>TGTTTTGATGAGAAAGCCCCTAATTTGAGTGTTTTTCCGTCGATTTGATTTTAAAGGTTTATATTCGAGTTTTTTTCGTCGGTT<br>TAATGAGAAGGCCTAAAATAGGAGTTTTTCTGGTTGATTTGACTAAAAAAGCCATGGAATTTTGTGTTTTTGATGTCGCTTTGG<br>TTCTCAAGGCCTAAGATCTGAGTTTCTCCGGTTGTTTTGATGAAAAAGCCCTAAAATTGGAGTTTTTATCTTGTGTTTTAGGTT<br>GTTTTAATCCTTATAATTTGAGTTTTTTCGTTGTTCTGATTGTTGTTTTTATGAATTTTGCAG |
| SlUbq10t (*Solanum lycopersicum* Ubiquitin 10 terminator) | GCTTGTTGTGGTTGTCTGGTTGCGTCTGTTGCCCGTTGTCTGTTGCCCATTGTGGTGGTTGTGTTTGTATGATGGTCGTTAAGG<br>ATCATCAATGTGTTTTCGCTTTTTGTTCCATTCTGTTTCTCATTTGTGAATAATAATGGTATCTTTATGAATATGCAGTTTGTG<br>GTTTCTTTTCTGATTGCAGTTCTGAGCATTTTGTTTTTGCTTCCGTTTACTATACCACTTACAGTTTGCACTAATTTAGTTGAT<br>ATGCGAGCCATCTGATGTTTGATGATTCAAATGGCGTTTATGTAACTCGTACCCGAGTGGATGGAGAAGAGCTCCATTGCCGGT<br>TTGTTTCATGGGTGGCGGAGGGCAACTCCTGGGAAGGAACAAAAGAAAAACCGTGATACGAGTTCATGGGTGAGAGCTCCAGCT<br>TGATCCCTTCTCTGTCGATCAAATTTGAATTTTTGGATCACGGCAGGCTCACAAGATAATCCAAAGTAAAACATAATGAATAGT<br>ACTTCTCAATGATCACTTATTTTTAGCAAATCAGCAATTGTGCATGTCAAATGATTTCGGTGTAAGAGAAAGAGTTGATGAATC<br>AAAATATCTGTAGCTGGATCAAGAATCTGAGGCAGTTGTATGTATCAATGATCTTTCCGCTACAATGATGTTAGCTATCCGAGT<br>CAAATTGTTGTAGAATTGCATACTTCGGCATCACATTCTGGATGACATAATAAATAGGAAGTCTTCAGATCCCTAAAAAATTGA<br>GAGCTAATAACATTAGTCCTAGATGTAACTGGGTGACAACCAAGAAAGAGACATGCAAATACTACTTTTGTTTGAAGGAGCATC<br>CCTGGTTTGACATATTTTTTCTGAATATCAAACTTTGAAACTCTACCTAGTCTAATGTCTAACGACAGATCTTACTGGTTTAAC<br>TGCAGTGATATCTACTATCTTTTGGAATGTTTTCTCCTTCAGTTATACATCAAGTTCCAAGATGCAGGTGTGCTTGATTGATGT<br>ACATGGCTGTGAGAAGTGCATCCTGATGTTCAGATGATGGTTCATTCTAATGTCTTTTCCTTCAATCAGTTTTCTCAGTCTGAC<br>TTAGCTTGTTTCATCTGCATGTTTGAATGTTCGTTTACTCATAGTAATTGCATTTTTGTAGCAGAACATATCATTGGTCATGGT<br>TTCAACTGTGCGCGAGTCTTATGCTTATTCAAACTAGGAAAGCCTCCGTCTAGAGGGTACACGAGTTGTTGCTCTGTGTGCGTC<br>AGTCCATAGTATTAATCTTGCTAGTTGTAGTATATTGTTTATGTGGACTCGGAATTCATCATATGCTCCTTCTTTGCATCAAGT<br>AAGGCAAGGTAATGTATAGAAGCTTTTTAACTCTTTCATGGAAGCTGGCCTTTGCCAGCATACCATCCAGAAGATATCAACCCT<br>GCATCTTGGCTGCCG |
| TiSGD<br>*T. iboga*<br>Strictosidine β-d-glucosidase<br>(This study) | ATGCATTCGTATTCAACAACTATGGAAACTACTCATAGCCCTCTTGTTGCTGCTATTGCTCCAAGATCAACTGCAGTAGCAGAC<br>ACGAAGAATTCTAACGGCACTAGACCAGCGAGCAAGGTCGTCCATCGTCGAGAATTCCCTGAAGATTTTATCTTTGGTGCCGGA<br>GGATCCGCTTATCAGTGTGAGGGAGCAGCTAATGAAGGTAATCGAGGCCCAAGTATATGGGATACTTTCACTCAGCAAACACCA<br>GGCAAAATAGCTGACAGGTCAAATGGAGACAAAGCCATCAATTCTTACCATATGTACAAGGAAGATGTTAAGATTATGAAGCAG<br>ACAGGCCTGGAAGCATATAGATTCTCAATTTCCTGGTCCAGAGTATTGCCAGGTGGAAGATTATCAGCTGGAGTTAATAAGGAG<br>GGAGTCAAGTTCTACCATGACTTTATAGATGAGCTTCTAGCCAATGGCATCAAACCCTTTGCAACTCTCTTCCACTGGGATGCT<br>CCCCAAGCCCTTGAAGACGAGTATGGTGGCTTCTTGAGTTCCCGGATTGTGGACGATTTTTGTGAGTATGCAGAATTTTGTTTT<br>TGGGAATTTGGTGACAAAGTGAAAAATTGGACAACATTCAATGAGCCTCACACCTATAGTGTCAACGGCTATACCATTGGTGAA<br>TTTGCACCAGGGAGAGGTGGATATGACAAAGGTGATCCAGGCACAGAACCCTATTTAGTGTCACACAATATCCTTCTTGCTCAC<br>AAAGCAGCTGTGGAAATATATAGGAAGAAATTTCAGGAATGTCAAGAAGGTGAAATTGGATTTGTAGTTAATGCAACTTGGATG<br>GAGCCTCTCAATCACAACAAGGCTGACATTGAAGCTCAGAAAAGAGCTCTTGATTTTATGCTTGGATGGTTTATGGAGCCATTA<br>ACAACCGGTGACTACCCAAAATCCATGAGAAACCTAGTTGGAGGACGTCTTCCAACATTTTCACCTGAAGAATCCGAAGGATTA<br>AAAGGCTGCTATGATTTTATTGGAATCAATTATTATACTGCTACTTACGTGACTGATGCAGTTAAATCTACCAATGAACGTTTA<br>GATTACAATACTGATGGTCAATATGTCACTACTTTTGACCGGGATAATGTACCCATCGGTCCAGTGTTGTATGGAGGGTGGCAG<br>CATGTTGTTCCAGTTGGACTCCACAAACTTTTGGTTTACACAAAGGACACATACCATGTTCCAGTCCTTTACGTCACGGAAAAT<br>GGTATGGTTGAAGAAAACAAAACCAGCATGCTGCTTCCAGAAGCTCGTCACGACACCAATAGGGTAGATTTTCACCGAGAGCAT<br>ATTTCAAGCGTGCGAGATGCTATTGATGATGGTGTGAATGTAAAAGGTTACTTTGCCTGGTCATTCTTCGACAACTTTGAGTGG<br>AATTTGGGTTTTACATGTCGTTATGGAATTATTCATGTTGATTTCGAAAGTTTTGCAAGATATCCGAAGGATTCAGCCATATGG<br>TACAAGAATTTCATTTTTGGAAAATCTCACACTTTGCCGGTTAAAAGACCTCGTGACGAAGATCGAGAAGCTGAGTTAGTCAAA<br>AGGCAAAAATCCCAAGCAGGCACAAATAATATGGTCTTTTGA |

| | |
|---|---|
| TiGS<br>*T. iboga*<br>Geissoschizine synthase<br>(This study) | ATGGCTGCAGAAACAGCCCAACTCGACCTCTCAGTGAAAGCTATTGGATGGGGTGCTACTGATTCATCTGGGGTTCTTCAGCCC<br>ATCAAGTTCTGCAGAAGGACCCCGGGTGAGCGCGATGTGAAGATAAGAGTGTTGTATTGTGGTCTCTGCAATTTCGATATGGAA<br>ATGGTCAGGAACAAATGGGGTTTCACTAGATATCCTTACGTATTTGGACATGAGACTGCGGGTGAGGTGGTAGAAGTAGGCAGC<br>AAGGTGCAGAGATTCAAGGTTGGGGACAAAGTGGCTGTGGGGTGCATGGTTGGATCTTGCGGCAAATGTTACAATTGTGAAAAT<br>GGTATGGAAAATTACTGCCCAGAGCCCAACATGGCTGATGGCTCTGTTTACCGTGAACAAGGAGAGCGTTCATATGGTGGTTGT<br>TCCAATGTCATGGTTGTTGATGAGAAATTCGTTCTCCGTTGGCCCGAGAACTTCCCACAGGACAAAGGAGTTCCTCTCCTGTGT<br>GCTGGGGTCGTTGTTTATAGTCCATTAAAACATTTTGGACTCGACAAACCTGGAATGCACATTGGTGTTTTTGGCCTTGGAGGT<br>CTTGGCTCTGTTGCTGTCAAGTTTATTAAGGCTTTTGGTTCTAAGGCAACTGTGATCAGTACATCAAGGCGTAAAGAGAAGGAA<br>GCCATCGAAGAGCATGGTGCAGATGCATTTGTTGTAAACAATGAGACTGAGCAATTGGAGGCCTTAGCAGGTACAATGGATGCC<br>GTTGTTGACACTACCCCTGGTGGACTCACTCCCATGTCACTCATGCTCAATTTGCTAAAGTTTGACGGAACAGTGATTATGGTT<br>GGTGCACCAGAAAGAGTATTTGACCTCCCAGTGCCACCGCTCATTCTTGGAAGGAAAAAGGTAGTTGGGAGCTCCACTGGAGGT<br>CTGAGGGAGTACCAAGAGGTGCTTGATTTTTCAGCCAAACACAACATTCTGTGTGATACTGAGGTCATCTCGATTGACTATCTC<br>AGCACTGCCATGGAGCGCGTCAGGAATCTGGATGTCAAATACCGATTCTCAATTGACATTGGCAATACATTGAAATATGATGAA<br>TGA |
| TiSBE<br>*T. iboga*<br>Sarpagan bridge enzyme<br>(This study) | ATGGAGATGATGAACTTCTCTTTCACCTCCCCAGTTTTCCTTCTCCTCTCCTTCTTTCTTCTCATGTTACTCAAACAACTAAGA<br>GGAAACAAAGGTGACAAGGGCAAAAAGTTGCCACCAGGTCCAAAAAAACTTCCCATTGTCGGAAATTTGCACCAGATGCTTGGA<br>TCACTGCCTCATCGTGTTCTCAAGAAATTGGCAGATCAATATGGACCGGTTATGCACTTACAGATCGGTGAACTTTCAGCAGTA<br>ATCATATCATCTGCACAAAAAGCAAAACAAGTCTTACAAACTCATGGCATCGCCGTTGCAGATCGGCCCCAAACTACTGTTGCT<br>AAGATCATGTTATACAATAGTCTGGGTGCCACTTTCGCTTCATATGGCGATTATTTGAAGCAATTGCGTCAAATTTATGCATTG<br>GAACTTCTAAGCCCCAAGAGTGTTCGATCGTTCTGGACTATCATGGAAGATGAGCTTTCCATTATGGTTACATCAATCAAGTCT<br>GACGTTGGAGAACCAGTGATTTTGCATGATAAAATGATGACTTATTTGTACGCCACACTTTGCAGAGCTACTGTTGGTAGCGTA<br>TGCAATGGACGAGAGACGTTGATAATGGCTGCAAGAGAAACTTCAGCGCTTTCTGCGGCCATTAGGATTGAAGATCTCTTCCCT<br>TCAGTAAAAATACTTCCTGTTGTCAGTGGACTAAAGTCAAGATTAACCAAACTGATGAAACAACTCGATACTGTTCTAGAGAAT<br>ATAATCAGTGAGCGGGAGAAGAACTTATTCTCATCCAACAACAACCAGCCATTGTCGGATGAAGAAGATATGCTGGGCGTTCTC<br>CTGATGTACAAGAACGGAAAGGGAAAAGACTCTAAATTCCGAATCACCAACAATGACATCAAGGCTATCATTTTTGAATTGATC<br>TTGGCTGGAACTCTAAGTTCAGCAGCTATAGTTGAGTGGTGCATGTCTGAGATGATGAAAAACCCTAGAGTCCTGAAGAAGGCA<br>CAGGATGAAGTGAGGCAAGTTCTCAACGGTAAGAAAACAATCAATGGAGATGATGTTACCAAAATGGATTATGTGAACATGGTA<br>GTCAAGGAATCTGTGAGATTACACCCTCCTGCTCCCCTATTGTTTCCAAGAGAATGCAGGGAAGAATTTGAGATTGACGGAATG<br>ACCATACCCAAAAAATCTTGGGTGATTATAAATTATTGGGCAGTGGGAAGAGACCCTGAACTTTGGGACGATGCCAACGAGTTT<br>AAGCCAGAGAGATTCAGCAACAAACCAATTGATTTCTATGGAAGCCATTTTGAGTTGATACCATTTGGGGCAGGAAGGAGGGTT<br>TGTCCAGGCATATTATTTGGAGTGACTAATGTTGAACTTTTGCTTGCAGCGCTTCTTTATCATTTTGACTGGAAACTCCCTGGA<br>AGCATGAAACCTGAAGAATTAGATATGACTGAGTTATTTGGTGCTGGTTGCATAAGGGAAAACCCATTGTGCCTAATTCCCAAC<br>ATCATTTCAGGAAATTAA |
| TiGO<br>*T. iboga*<br>Geissoschizine oxidase<br>(This study) | ATGGAGCTCTCTTTCTCCTCCCCGGCTCTCTACATCCTCTACTTCATGCTTTTCTTTCTTGTAAAGCAACTGTTAAAGCCCAAA<br>AGTCACAAGAAATTGCCACCAGGTCCTAGGACGCTACCCATAATCGGAAACTTGCACCAGCTTATGGGACCATTGCCTCATCGC<br>ACCCTCAAGGAGTTGTCCGATAAACATGGACCTCTCATGCACTTGAAGATGGGTGAACGTTCAGCAATTATCGTATCAGATGCA<br>AGAATGGCAAAAATAGTCCTGCACAATCATGGGCTGGCAGTGGCGGACCGGTCGGTGAATACTGTTGCTAGCATCATGACCTAC<br>AACAGTTTGGGGGTTACATTTGCTCAATACGGCGATTACTTGACAAAGCTGCGTCAGATTTACACTTTGGAACTCCTTAGTCCC<br>AAAAAAGTTCGATCATTTTACAGTTGTTTTGAAGACGAGCTTGATGGTTTTGTGAAATCAATCAAGTCTCAAGTCGGACAACCC<br>ATGGTTTTGTACGAGAAAGCTTCTACTTACTTGTATGCCACAATTTGCAGAACTATTTTTGGAAGCATATGCAAGGAGAGGGAG<br>AAGATGATCAAGATAGTGAAAAGAACATCGTTACTCTCTGGAACCCCTCTCAGGTTGGAGGATCTGTTCCCTTCAATGAGCGTG<br>TTTTGTAGGTTCAGCAAAACACTGAACCAATTGAGGGATCTGTTACAGGAAATGGATGATATTCTCGAGGATATTATCATTGAG<br>CGTGAAAAGGCCGCTGAAATCTCGACAGAGGCCAAGGACGACGAAGATATGCTCAGCGTTCTCTTGAGGCATAAATGGCACAAT<br>CCCAGCGGCGCAAAGTTCAGAATCACAAACGCAGACATCAAAGCCATCATTTTTGAGTTGATCTTAGCTGCAACTCTAAGCGTA<br>GCAGATGTTACAGAATGGGCTATGGTTGAAATACTGAGGGATCCCAAAAGTTTGAAGAAGGTATACGACGAAGTCAGAGAAGTC<br>TGCAAGGAGAAGAAAAGAGTCACCGGATATGACGTTGAGAAATTAGAATATATGCACCTTTGTTTTAAGGAATCTACAAGAATA<br>CACCCTGCTGCTCCTCTCTTAGTCCCCAGAGAATGCAGAGAAGACTTCGAGGTCGACGGATACACTGTTCCAAAAGGAGCATGG<br>GTGTTGACAAACTGCTGGGCTGTCCAGATGGATCCCAAGATCTGGCCGGAGCCTGAGAAGTTTGATCCAGAGCGATACATCCGT<br>GATCCCATGGATTTCTACGGAAGCAACTTTGAGTTGATACCCTTTGGTACGGGAAGGAGAGGATGTCCAGGAATCTTGTTTGGA<br>GTGACCAATGCTGAGCTTCTCTTAGCTGCAATGTTCTATCATTTTGATTGGGAAATTGCAGACGGGAAAAAACCAGAAGAAATA<br>GATCTTACCGAGGACTTTGGTGCTGGTTGCATTATGAAATATCCTTTGAAACTTGTTCCACATCTTGCCAACGAATAG |
| TiRedox1<br>*T. iboga*<br>Redox enzyme 1<br>(This study) | ATGGCTGATCGCGTCAAGACCATCGGATGGGCAGCTCATGACAGTTCCGGGATCCTCTCTCCCTTCGAATTCACCCGAAGGGCA<br>ACTGGTGATGAAGATGTAAGGTTTAAGGTGCTATATTGTGGTATCTGTCACTCCGACCTTCACAATGTCAAGAACGAAATGGGC<br>TTCACTTCATACCCTTGTGTCCCAGGGCATGAAGTTGTAGGTGAAGTGACGGAGGTTGGAAGCAAAGTCAAGAAATTCAAAGCT<br>GGAGACAAAGTTGGGGTAGGATTATTCGTCGATTCGTGTCGCGAATGCGAGCAGTGTAAAAATGATCTAGAGCCCTACTGCCCC<br>AAGTTGAAAATGGCCTATCTGTCTCCTGATGATGATGGAACCATTATTCAGGGAGGATACTCTAATGAGATGGTAGTCAAGGAA<br>CACTATGTACTTCGGTGGCCGGAAACTCTTCCTCTCGATGCTGGTGCACCGCTAGTCGGCGCCGGTAGCACTGTCTATAGTCCA<br>ATGAAGTATTATGGACTTGATAAGCCAGGACAACATTTGGGAGTTGTTGGCCTCGGTGGACTCGGCCACTTAGCTGTGAAATTC<br>GCCAAGGCTTTTGGACTCAAAGTCACCGTGATCAGTACTTCCCCTGGGAAAAAGGAAGAAGCCATCACTCACCTAGGAGCTGAC<br>TCATTTTTGGTCAGCACTGATCAAGAGCAAATGCAGAAAGCAACGAGCACCTTAGACGGTATCATTGACACAGTATCAGCACCT<br>CATGCAGTGATGCCATTGTTTTTCTTATTGAAACCTCATGGGAAGCTTATTGTAGTCGGGGCACCAAATAAGCCACTTGAAATA<br>GATGTTCCATTTCTTCTTATGGGAAGAAAAATGCTGGGAGCTTCCGCTGTCGGGGGAATGAAGGAGACTCAAGAAATGCTTGAT<br>TTTGCAGCACAACACAATATCTCTGCAGATGTTGAAGTTGTGCCAATGGACTACGTGAACAAAGCTATTGAGCGCCTTGAGAAA<br>GGCGATGTTAGGTATCGATTTGTTCTTGATATTGGGAATACAATGGGCGCTGCCTAG |
| TiRedox2<br>*T. iboga*<br>Redox enzyme 1<br>(This study) | ATGGAAAGGGAAGTTCAAATCCCAGAAATCGAGTTGAACTCCGGGCACAAGATGCCACTTGTGGGCTTCGGGACATGTGTGCCA<br>GATCCCATTCCGCCATTGGAGGAACTTGTAGCAAGCTTCTTGGAGGCTATCAAAGTTGGGTATCGCCACTTTGACACCGCATCA<br>AGCTATGGCACAGAGGAGGCCCTTGGTAAAGCTGTGGCTCAAGCAATAGAGAGTGGCTTGGTTAAGGGCAGAGAGGAATTCTTC<br>ATCAGTGATAAATTGTGGTGTGAAGATGCTGATCATGATCTTATCTTGCCAGCCCTCAAGAAAACACTAGGGAATCTGGGGCTT<br>GATTATTTGGATCTTTATATGATACATATGCCAGTAAGAGTGAGGCAGGGTGCTCCAATGTTCAATTATTCAAAAGAAGATTTA<br>CTCCCTTTTGATATACAAGGGACATGGAAGGCCATGGAAGAATGCAGCAAATTGGGCTTGGTCAAGTCTATTGGTGTAAGCAAC<br>TACACTTGTGAAAAACTCGCAAAACTCTTAGAAAATGCCACCATTCCTCCTGCAGTTAATCAGGTGGAGATGAACGTTGTTTGG<br>CAACAGAGAAAACTGCTGCCATTTTGCAAAGAGAAAAACATTCATGTCACCGCATGGTCTCCTCTGTTATCTTATGGTGCCATT<br>TGGGGCAGTAATGCAGTGATGGAGAATCCAGTCCTCGTAGATATTGCAGCTTCCAAGAACAAGATCGTGGTTCAGGTTGCACTA<br>AGATGGATATATGAGCAAGGAGCAAGTTTCATCATGAGGACCTTTAAAAAGGAGAGAATGTTCGAAAATGTTCAGATTTTCGAC<br>TGGGAACTCACTAAAGAAGAATTGGACAAGATTCAACAAATCCCTCAACGTAGGGGCACTTTGGGAGAAGATTTCATCCATCCA<br>GAGGGGCCAATTAAATCAGTGGAGGAACTCTGGGATGGAGATTTGTAA |
| TiSAT<br>*T. iboga*<br>Stemmadenine O-<br>acetyltransferase<br>(This study) | ATGACTTCCCAGATGGATGTAGTATCAGAGGAGCTCATTCAACCATCATCTCCAACACCCGAAAGCTTAAAACACCATAAACTT<br>TCACTCCTAGACCAAGTTTCGTTGACATGTCATATTCCTATTATTTTTTCTACCCAAATCAGTTGATCGCCTCAAATGTCGAC<br>CGTGCCCAAAGATCTGAGCACCTGAAACAATCTTTGTCCAAAGTGTTAACTAATTTTTATGTTTTAGCAGGAAGAATTAATCTA<br>AATTCTTCCATAGATTGTAATGATTCTGGAGTTCCGTTCATCGAAGCTCGAGTTCAGGCTGAACTCTCACAGGCAATTCAGAAC<br>GTCGCCATAGAGGAATTGAATCAATATCTTCCGATCGAACCTTATCCCGGCGGGCAAAGCGAGGTGAAAAAGGATATTCCCCTT<br>GCTGTGAAAATCAGTTTCTTTGACTGTGGAGGCGCAGCAATTGGTGTCTGCTTGTCACATAAGATAGCTGATATATTGTCCTTG<br>GCCACCTTCCTCAATGCTTGGACTGCCACATGCCGAGGGGAAACCCATGTTGTGCTGCCTAATTTTGACTTGGGATCCCATCAT<br>TTTCCGCCAATGGACAACATCCCGGCACCTGACTGGGTACCGGATGAAAAGATTGTGATGAAAAGATTCATTTTTGATAAAGAA |

| | |
|---|---|
| | AAATTAGCAGCCCTCAAAGCACAAGCTTCCTCAGCCTCGGAGGTGAAGAATCCCAGTCGGATAGAAGTTGTTACGGCTTTTATA<br>TGGAAGCACTTCATTGACGTGACCCGAGCGAAATTGGATACTAAAAACAAGTTTGCCGCAGCTCAGGCAGTGAACTTGAGGACA<br>AGAATGAGTACACCCCTTCCTCAGTCTGCTATGGGGAACATCGCCACACTTGCATTCGCTGTTGCAGAGGAGGACAAGGATTTT<br>CACGATCTTGTTGGTCCATTGAGAACTGGCCTAGGGAAAATTGACAACGAACATGTTAAGGACTTACAAAAGGGAGTGACTTAT<br>TTGGATTTTGAAGCGGAACCACAAGAATTGTCTTCTTTCAGCAGTTGGTGTAGGCTTGGCTTTTATGAAATGGACTTTGGGTGG<br>GGGAAGCCTCTTTCTGTGTGCACTGCAAATTTGCCCATGAAGAATACCGTATATATGATGGATACTAGATCTGGAGATGGAATA<br>GAAGCATGGATTAGCATGGCGGAAGATGAAATGGCTATGCTTCCCACTGAATTCCTTTCACTTGTAGACAGTGATTTTAGCAAG<br>TGA |
| CrSGD<br>*C. roseus*<br>Strictosidine β-d-glucosidase<br>(AF112888.1) | ATGGGATCTAAAGATGATCAGTCCCTTGTTGTTGCCATTTCTCCAGCTGCTGAACCAAATGGAAATCATTCTGTCCCCATCCCA<br>TTCGCCTACCCCAGTATCCCCATTCAACCTAGAAAGCACAACAAGCCCATCGTTCATCGTCGAGATTTCCCCTCAGATTTCATC<br>TTGGGTGCCGGAGGATCTGCTTATCAGTGTGAGGGTGCATATAATGAAGGCAACCGCGGTCCCAGTATATGGGATACTTTCACA<br>AACCGATATCCAGCCAAAATAGCTGATGGATCTAATGGCAATCAAGCCATCAATTCTTACAATTTGTACAAGGAAGATATCAAG<br>ATTATGAAGCAAACAGGCTTGGAATCATATAGGTTTTCAATTTCATGGTCAAGAGTATTGCCAGGTGGGAATCTATCCGGTGGA<br>GTGAATAAAGATGGTGTCAAGTTCTATCATGACTTTATAGATGAGCTTCTAGCCAATGGCATCAAACCCTTTGCAACTCTCTTC<br>CACTGGGATCTTCCCCAAGCTCTTGAAGACGAGTATGGAGGCTTCTTGAGTGATCGAATTGTGGAAGATTTTACGGAGTATGCA<br>GAATTTTGCTTTTGGGAATTCGGTGACAAAGTAAAATTTTGGACGACTTTCAATGAACCACATACTTATGTTGCAAGTGGATAT<br>GCCACTGGTGAATTTGCACCAGGAAGAGGTGGTGCAGATGGCAAGGGGGAACCTGGCAAAGAACCCTATATAGCGACACATAAT<br>TTACTTCTTTCTCACAAAGCTGCTGTGGAAGTATATAGGAAAAATTTTCAGAAATGTCAAGGAGGTGAAATTGGAATTGTACTT<br>AATTCAATGTGGATGGAGCCTCTCAATGAAACCAAAGAAGATATTGATGCTCGGGAAAGGGGTCTTGATTTCATGCTCGGATGG<br>TTCATAGAGCCATTAACAACGGGTGAATACCCAAAATCCATGAGAGCTCTTGTAGGAAGCCGTCTTCCAGAATTTTCAACAGAA<br>GTTTCCGAAAAATTAACAGGATGCTATGATTTTATCGGAATGAATTATTATACAACTACTTATGTTTCTAATGCAGACAAAATT<br>CCCGATACTCCGGGTTACGAAACAGATGCTCGAATTAATAAGAATATTTTTGTCAAAAAAGTTGATGGGAAGGAAGTGCGCATT<br>GGTGAACCGTGCTATGGGGGATGGCAGCATGTTGTTCCATCTGGACTCTACAATCTCTTGGTTTACACTAAGGAGAAATACCAT<br>GTTCCAGTGATTTATGTCTCAGAATGTGGTGTGGTTGAGGAAAATAGAACCAACATATTACTTACAGAAGGTAAAACCAACATA<br>TTACTTACAGAAGCTCGTCACGATAAACTCAGGGTTGATTTTCTACAAAGTCATCTCGCTAGCGTGCGAGATGCTATTGATGAT<br>GGTGTGAATGTAAAAGGATTCTTTGTTTGGTCATTCTTCGACAACTTCGAATGGAATTTGGGATATATATGCCGTTATGGAATT<br>ATCCATGTTGATTATAAAACTTTTCAAAGATATCCAAAGGATTCTGCCATATGGTACAAGAATTTCATTAGTGAAGGATTTGTT<br>ACGAATACAGCTAAAAAGAGATTCCGAGAAGAAGATAAACTAGTTGAGTTAGTCAAGAAGCAAAAATACTAA |
| CrGS<br>*C. roseus*<br>Geissoschizine synthase<br>(MF770507.1) | ATGGCCGGAGAAACAACCAAACTCGACCTTTCAGTGAAGGCTGTGGGATGGGGTGCTGCAGATGCATCTGGTGTCCTTCAGCCC<br>ATCAAGTTCTATAGAAGAGTCCCTGGTGAACGGGATGTGAAGATTAGAGTTTTGTACTCTGGTGTTTGCAATTTCGATATGGAA<br>ATGGTCAGAAACAAGTGGGGTTTCACCAGATATCCTTATGTGTTTGGACATGAAACTGCCGGTGAGGTGGTAGAAGTTGGCAGC<br>AAAGTAGAGAAATTCAAGGTTGGAGACAAGGTAGCTGTGGGATGTATGGTCGGATCTTGTGGTCAATGTTATAATTGTCAAAGT<br>GGAATGGAGAATTACTGCCCAGAGCCCAATATGGCTGATGGATCTGTTTACCGTGAGCAAGGGGAACGATCCTATGGGGGTTGT<br>TCAAATGTGATGGTTGTTGATGAAAAGTTCGTCCTCCGATGGCCCGAAAACTTGCCTCAAGATAAAGGGGTTGCTCTCCTTTGT<br>GCTGGGGTTGTTGTTTATAGCCCAATGAAACATTTGGGACTCGATAAGCCAGGAAAGCATATTGGGGTTTTCGGGCTGGGAGGT<br>CTTGGTTCTGTTGCTGTTAAGTTTATTAAGGCTTTTGGTGGTAAGGCTACTGTTATTAGTACATCAAGGCGTAAGGAGAAGGAA<br>GCCATTGAAGAACATGGTGCTGATGCTTTTGTTGTCAACACTGACTCTGAGCAATTGAAGGCTCTGGCAGGTACTATGGATGGT<br>GTTGTGGACACCACCCCAGGTGGCCGCACTCCTATGTCACTTATGCTCAATTTGCTCAAGTTTGACGGCGCGGTTATGCTCGTA<br>GGTGCACCGGAGTCGCTATTTGAGCTCCCTGCGGCACCTCTCATTATGGGAAGGAAAAAGATAATCGGAAGTTCCACTGGAGGT<br>CTCAAAGAGTACCAAGAAATGCTTGATTTCGCAGCCAAACATAACATTGTATGTGATACTGAAGTTATTGGGATTGACTATCTC<br>AGCACTGCTATGGAACGTATCAAGAATTTGGATGTCAAGTACCGTTTTGCGATTGACATTGGAAATACATTGAAATTTGAGGAA<br>TGA |
| CrSBE<br>*C. roseus*<br>Sarpagan bridge enzyme<br>(PV446798.1) | ATGGATGAGATGATGAACTTCTCCCTCACCTCTCCCATTTCCTTTCTCCTCTCCTTACTTCTTATCCTATTTCTTATCCTATTT<br>ATGATGAAAGGAAACCAAACCCACAATGGCAAAAAATTACCACCGGGTCCCAAAAAACTTCCTATTATAGGAAACCTCCATCAA<br>ATGATAGGTTCACATCCTCATCGTGTTCTAAAAAAATTGGCCGATAAATATGGACCTATTATGCACTTGCAAATGGGCCAACGA<br>TCAGCCGTTGTTATATCATCAGCGGAAAAAGCAAAAGAAATATTGAACATTTATGGTGTTCAAGTTGCTGATAGACCTCAACTC<br>ATCGTATCTAAAATTATGTTGTACAATAGTTTGGGTGTCACTTTTGCTCCTTATGGTGACTACTTAAAGCAATTGCGTCAAATT<br>TATGCCGTAGAACTTCTTAGCCCTAAAACAGTTAGGTCCTTTTGGACAATTATGGAAGATGAAGTTTCCACAATGGTTAACTCA<br>ATTAAATCTGAAATTGGTCAACCAATAATTTTGCATGATAAAATGATGACTTATTTGTATACTATGCTTTGTAGAGTTACATTT<br>GGTGGCGTATGTAATGGACGCGAGACACTAATAATGGCAGCTAAAGAAACGTCAGCGCTTTCTGCTGCTATTAGGATTGAGGAT<br>TTGTTTCCTTCAGTGAAAATACTTCCTTTAATTAGTGGATTAAAGTCAAGATTAAACAATTTGTTGAAAACACTTGATACCATT<br>CTTGAGGATATTATCAGTGTTCGTGAGAAGAAATTATTAACTCAGCCATTGTTGGATGATGAAGATATGTTGGGAGTTCTCCTC<br>AAGTACAAAAATGGAAAGGGAAAAGATACTAAATTTAGAGTCACCAACAACGACATCAAGGCAGTCATTTTTGAAATTATCTTA<br>GCTGGCACCCTAAGTTCATCAGCTATAGTAGAATGGTGTATGTCTGAAATGATAAAAAACTCTGAAAAGTTTAAAAAGGCACAA<br>GATGAAGTTAGGAAAGTTTTAAAGGGTAAAAAAACAATTAGTGGAAGTGATGTTGGTAAAATGGAATATGTTAAAATGGTGGTT<br>AAGGAATCTTTGAGATTACACCCTCCTGCTCCCATATTGTTTCCAAGAGAATGTAGGGAAGAATTTGAGATAGATGGAATGACT<br>ATACCTAAAAAAACTTGGTTGATTGTAAACTACTGGCCAATAGGAAGAGATCCCAGACTTTGGGACGACGCCGACAAGTTTGAG<br>CCGGAGAGGTTCAGTAATAGCAGCATTGATTTCAATGGAAGCCATTTTGAGTTGATACCATTTGGTGCTGGAAGAAGGATTTGT<br>CCTGGAATATTACTTGGAACAACAAATGTTGAGCTTTTACTTGCTACATTTCTCTATTATTTTGATTGGAAACTTCCTCAAGGT<br>ATGAAACCCGAAGAATTAGATATGAATGAGGTATTTGGTGCTGGTTGCATAAGGGAAAGTCCATTGTGTCTCATTCCTAGCATC<br>TCATCAGCAGTTGAAGGAAATTAA |
| CrGO<br>*C. roseus*<br>Geissoschizine oxidase<br>(JN613015.1) | ATGGAGTTTTCTTTCTCCTCACCAGCTCTCTACATAGTTTATTTCTTGCTTTTCTTTGTGGTAAGGCAATTATTGAAACCCAAA<br>AGTAAGAAAAAATTGCCACCAGGTCCAAGAACACTACCCTTAATTGGAAACCTTCATCAACTCTCGGGACCTTTACCTCATCGT<br>ACCCTAAAAAATTTGTCCGATAAACATGGTCCTTTGATGCACGTGAAAATGGGCGAACGTTCGGCAATTATAGTATCAGATGCA<br>AGAATGGCAAAAATAGTTCTTCATAATAACGGTTTAGCCGTTGCAGATCGGTCAGTAAATACTGTCGCAAGTATTATGACTTAT<br>AATAGTTTGGGTGTTACCTTTGCTCAGTATGGAGATTACTTAACAAAATTACGTCAAATCTATACTTTGGAACTTTTAAGTCAG<br>AAAAAAGTTCGATCTTTCTACAGTTGTTTTGAAGATGAACTCGATACTTTTGTTAAGTCAATTAAGTCTAACGTTGGACAACCT<br>ATGGTTTTGTACGAGAAAGCTTCTGCTTATTTGTATGCTACCATTTGTAGAACTATTTTTGGGAGTGTTTGCAAGGAAAAGGAG<br>AAAATGATTAAGATAGTGAAGAAAACGTCGTTACTGTCTGGAACACCACTTAGATTGGAGGATCTTTTTCCTTCTATGAGCATT<br>TTTTGTAGATTTAGTAAAACTTTGAATCAATTGAGAGGACTTTTGCAAGAAATGGATGATATTTTGGAAGAAATCATAGTTGAA<br>AGGGAAAAGGCTTCTGAAGTCTCAAAAGAGGCAAAGGATGATGAAGATATGCTTAGTGTTCTGTTGAGGCATAAATGGTACAAT<br>CCAAGTGGTGCTAAATTCAGAATCACTAATGCAGACATCAAAGCCATTATCTTTGAGTTGATCTTAGCAGCAACCCTAAGTGTA<br>GCAGATGTTACAGAATGGGCTATGGTTGAAATTCTAAGGGATCCCAAAAGTTTGAAAAAAGTGTACGAAGAAGTACGAGGTATT<br>TGTAAAGAAAAGAAAAGAGTGACAGGATACGACGTTGAAAAAATGGAATTTATGCGTCTTTGTGTGAAGGAATCAACAAGAATA<br>CATCCAGCTGCTCCTCTTTTAGTTCCTAGAGAATGCAGAGAAGATTTTGAAGTAGATGGATACACAGTTCCAAAAGGGGCTTGG<br>GTTATAACAAATTGCTGGGCTGTTCAAATGGATCCTACAGTATGGCCCGAACCAGAGAAGTTTGATCCAGAAAGATATATTCGT<br>AACCCAATGGATTTCTATGGAAGTAACTTTGAATTGATACCATTTGGGACAGGTAGAAGAGGATGTCCTGGTATTTTATATGGA<br>GTGACAAATGCTGAGTTTATGTTGGCTGCTATGTTTTATCATTTTGATTGGGAAATAGCAGATGGTAAAAAACCAGAAGAAATT<br>GATTTGACTGAGGACTTTGGTGCTGGTTGCATTATGAAGTATCCTTTGAAATTGGTTCCTCATCTTGTTAACGATTAG |
| CrRedox1<br>*C. roseus*<br>Redox enzyme 1<br>(MF770509.1) | ATGGCTGATCGCGTGAAGACCGTAGGATGGGCAGCTCACGACAGCTCCGGCTTCCTCTCTCCCTTCCAATTCACTCGAAGGGCA<br>ACAGGTGAAGAAGATGTGAGGTTGAAGGTGTTGTACTGTGGTGTCTGTCACTCAGACCTTCATAACATCAAGAACGAAATGGGA<br>TTCACCTCCTACCCTTGTGTCCCCGGGCATGAGGTTGTGGGGGAAGTGACGGAAGTGGGGAACAAAGTAAAGAAATTCATAATT<br>GGTGATAAAGTTGGGGTTGGATTATTCGTTGACTCATGTGGCGAATGCGAACAATGTGTGAATGATGTAGAAACCTATTGTCCC<br>AAATTGAAAATGGCTTATTTATCCATTGATGATGATGGAACTGTGATTCAAGGAGGGTACTCAAAGGAAATGGTCATCAAAGAA |

| | |
|---|---|
| | CGCTACGTTTTCCGGTGGCCGGAAAATCTTCCTCTACCCGCCGGTACACCGCTTCTGGGTGCCGGTAGTACAGTTTATAGTCCA<br>ATGAAATACTATGGACTTGATAAGTCAGGACAACATCTAGGAGTTGTTGGCCTTGGTGGACTTGGTCATTTAGCTGTTAAATTT<br>GCAAAGGCTTTTGGACTTAAAGTTACTGTCATTAGTACCTCTCCTAGCAAGAAGGATGAAGCCATCAATCACCTTGGTGCTGAT<br>GCATTTTTAGTTAGCACTGATCAAGAGCAAACCCAGAAAGCAATGAGCACAATGGATGGCATAATTGACACAGTATCAGCTCCT<br>CATGCATTGATGCCATTGTTTTCCCTATTGAAACCAAATGGGAAGCTAATTGTTGTTGGTGCACCAAATAAACCAGTTGAATTA<br>GATATTCTATTTCTTGTCATGGGAAGGAAAATGCTCGGAACATCTGCTGTTGGGGGAGTGAAGGAGACACAAGAGATGATCGAT<br>TTTGCAGCAAAACATGGTATAGTTGCAGATGTAGAAGTTGTGGAAATGGAGAATGTGAACAATGCAATGGAGCGGCTTGCGAAG<br>GGGGATGTTAGATATAGATTTGTTCTTGATATTGGGAATGCAACAGTAGCTGTCTGA |
| CrRedox2<br>*C. roseus*<br>Redox enzyme 1<br>(MF770510.1) | ATGGAAAAGCAAGTTGAGATCCCTGAAGTAGAATTGAATTCAGGACATAAAATGCCAATTGTGGGATACGGAACATGCGTGCCA<br>GAACCCATGCCACCGTTGGAAGAACTAACCGCAATCTTCTTAGATGCAATAAAGGTTGGTTACCGGCACTTCGACACGGCTTCG<br>AGTTACGGCACAGAGGAGGCACTTGGTAAAGCCATAGCTGAGGCTATAAACAGCGGTTTGGTTAAAAGTAGGGAGGAATTCTTC<br>ATCAGTTGTAAGCTGTGGATTGAAGATGCAGATCATGACCTTATCTTGCCTGCCCTCAACCAGTCACTTCAGATTCTTGGGGTT<br>GATTATTTGGATCTATATATGATACACATGCCGGTGAGAGTGAGGAAAGGTGCTCCCATGTTCAATTATTCAAAAGAGGATTTC<br>CTTCCATTTGACATACAAGGTACATGGAAGGCCATGGAAGAGTGCAGCAAACAAGGATTGGCCAAGTCTATTGGTGTCAGCAAC<br>TACTCTGTTGAAAAACTCACTAAACTCCTAGAAACCTCCACTATTCCCCCCGCCGTTAATCAGGTTGAGATGAATGTAGCTTGG<br>CAACAGAGAAAATTGCTGCCATTTTGCAAAGAGAAAAACATTCATATAACATCATGGTCTCCACTCCTATCTTATGGTGTCGCT<br>TGGGGAAGTAATGCTGTCATGGAAAATCCTGTTCTCCAACAAATTGCGGCTTCCAAAGGCAAAACTGTGGCACAGGTGGCACTA<br>AGATGGATATACGAGCAAGGAGCAAGTCTCATTACAAGGACTTCCAACAAGGACAGAATGTTTGAAAATGTTCAAATTTTTGAT<br>TGGGAACTCAGTAAAGAAGAATTGGATCAAATTCATGAAATCCCACAACGTAGGGGTACTTTAGGTGAAGAATTTATGCATCCA<br>GAAGGACCGATCAAATCCCCAGAGGAGCTTTGGGATGGAGACTTGTGA |
| CrSAT<br>*C. roseus*<br>Stemmadenine O-<br>acetyltransferase<br>(MF770511.1) | ATGGCACCCCAGATGCAGATATTGTCAGAGGAACTGATTCAACCATCATCTCCGACACCCCAAACCTTGAAAACCCATAAACTT<br>TCCCATCTTGATCAAGTTTTATTAACATGTCATATCCCTATTATTCTCTTTTATCCAAATCAATTGGACTCAAATCTCGATCGA<br>GCCCAAAGATCTGAGAATCTAAAACGATCTTTATCAACAGTGTTAACTCAATTTTACCCTTTAGCCGGAAGAATCAATATAAAT<br>TCTTCCGTAGATTGTAATGATTCCGGAGTCCCTTTTCTTGAAGCTCGAGTTCATTCCCAACTCTCAGAAGCAATTAAAAATGTC<br>GCCATAGATGAACTCAATCAATACCTGCCATTCCAACCTTATCCCGGTGGGGAGGAAAGTGGGTTGAAAAAAGATATCCCCTTA<br>GCTGTAAAAATCAGTTGTTTCGAATGTGGCGGAACAGCAATTGGGGTCTGTATTTCCCACAAGATTGCCGATGCGTTGTCCTTG<br>GCTACCTTCCTCAATTCATGGACCGCAACATGCCAGGAAGAGACTGATATTGTTCAACCTAATTTCGATCTGGGATCCCATCAT<br>TTTCCGCCTATGGAAAGCATCCCAGCACCTGAATTTCTACCGGATGAAAATATTGTGATGAAACGGTTCGTGTTCGATAAAGAA<br>AAATTAGAAGCTCTAAAAGCCCAATTAGCTTCCTCTGCAACAGAAGTGAAGAATTCAAGTCGGGTACAAATTGTTATTGCTGTT<br>ATATGGAAGCAATTCATTGACGTGACCCGGGCAAAATTCGATACCAAGAACAAATTAGTAGCAGCTCAAGCAGTGAATTTGAGA<br>TCAAGAATGAATCCACCATTTCCTCAATCTGCTATGGGGAATATAGCCACAATGGCATATGCCGTTGCAGAGGAGGATAAGGAT<br>TTTTCAGATCTTGTCGGTCCATTGAAAACCAGTCTTGCAAAAATTGATGATGAACATGTTAAAGAATTACAAAAAGGAGTAACA<br>TATTTGGATTATGAAGCTGAACCACAAGAATTGTTCTCTTTTAGTAGTTGGTGCAGGCTTGGGTTTTATGATTTGGATTTTGGA<br>TGGGGAAAGCCTGTTTCTGTTTGTACAACAACTGTGCCTATGAAGAATTTGGTATATTTGATGGATACAAGAAATGAAGATGGA<br>ATGGAAGCATGGATCAGCATGGCTGAAGATGAGATGTCTATGCTTTCCTCTGATTTTCTTTCACTTCTGGACACTGATTTTAGC<br>AATTGA |

**Table S2**. Primer sequences used for functional characterization of biosynthetic genes in this study.

| Gene | Plasmid | Primer direction | Sequence (5'-3') |
|---|---|---|---|
| *Nicotiana benthamiana* single transcriptional unit constructs (In-Fusion cloning) | | | |
| P19 (TBSV) | 3Ω1 | Forward | **TTTATGAATTTTGCAGCTCG**ATGGAACGAGCTATACAAGGAAA |
| | | Reverse | **GACAACCACAACAAGCACCG**TCACTCGCTTTCTTTTTCGAAGG |
| GFP | 3Ω1 | Forward | **TTTATGAATTTTGCAGCTCG**ATGGTGAGCAAGGGCGAG |
| | | Reverse | **GACAACCACAACAAGCACCG**TCACTTGTACAGCTCGTCCATGC |
| CrSGD | 3Ω1 | Forward | **TTTATGAATTTTGCAGCTCG**ATGGGATCTAAAGATGATCAGTC |
| | | Reverse | **GACAACCACAACAAGCACCG**TTAGTATTTTTGCTTCTTGACTAACT |
| CrGS | 3Ω1 | Forward | **TTTATGAATTTTGCAGCTCG**ATGGCCGGAGAAACAA |
| | | Reverse | **GACAACCACAACAAGCACCG**TCATTCCTCAAATTTCAATGTATTTC |
| CrSBE | 3Ω1 | Forward | **TTTATGAATTTTGCAGCTCG**ATGGATGAGATGATGAACTTCTC |
| | | Reverse | **GACAACCACAACAAGCACCG**TTAATTTCCTTCAACTGCTGATGA |
| CrGO | 3Ω1 | Forward | **TTTATGAATTTTGCAGCTCG**ATGGAGTTTTCTTTCTCCTCAC |
| | | Reverse | **GACAACCACAACAAGCACCG**CTAATCGTTAACAAGATGAGGAA |
| CrRedox1 | 3Ω1 | Forward | **TTTATGAATTTTGCAGCTCG**ATGGCTGATCGCGTGAAGA |
| | | Reverse | **GACAACCACAACAAGCACCG**TCAGACAGCTACTGTTGCAT |
| CrRedox2 | 3Ω1 | Forward | **TTTATGAATTTTGCAGCTCG**ATGGAAAAGCAAGTTGAGATCC |
| | | Reverse | **GACAACCACAACAAGCACCG**TCACAAGTCTCCATCCCAAA |
| CrSAT | 3Ω1 | Forward | **TTTATGAATTTTGCAGCTCG**ATGGCACCCCAGATGCA |
| | | Reverse | **GACAACCACAACAAGCACCG**TCAATTGCTAAAATCAGTGTCCA |
| TiSGD | 3Ω1 | Forward | **TTTATGAATTTTGCAGCTCG**ATGCATTCGTATTCAACAACTATGG |
| | | Reverse | **GACAACCACAACAAGCACCG**TCAAAAGACCATATTATTTGTGCCTG |
| TiGS | 3Ω1 | Forward | **TTTATGAATTTTGCAGCTCG**ATGGCTGCAGAAACAGC |
| | | Reverse | **GACAACCACAACAAGCACCG**TCATTCATCATATTTCAATGTATTGCCA |
| TiSBE | 3Ω1 | Forward | **TTTATGAATTTTGCAGCTCG**ATGGAGATGATGAACTTCTCTTTCA |
| | | Reverse | **GACAACCACAACAAGCACCG**TTAATTTCCTGAAATGATGTTGGGAATTA |
| TiGO | 3Ω1 | Forward | **TTTATGAATTTTGCAGCTCG**ATGGAGCTCTCTTTCTCCTC |
| | | Reverse | **GACAACCACAACAAGCACCG**CTATTCGTTGGCAAGATGTG |
| TiRedox1 | 3Ω1 | Forward | **TTTATGAATTTTGCAGCTCG**ATGGCTGATCGCGTCAA |
| | | Reverse | **GACAACCACAACAAGCACCG**CTAGGCAGCGCCC |
| TiRedox2 | 3Ω1 | Forward | **TTTATGAATTTTGCAGCTCG**ATGGAAAGGGAAGTTCAAATCC |
| | | Reverse | **GACAACCACAACAAGCACCG**TTACAAATCTCCATCCCAGAGT |
| TiSAT | 3Ω1 | Forward | **TTTATGAATTTTGCAGCTCG**ATGACTTCCCAGATGGATGTAG |
| | | Reverse | **GACAACCACAACAAGCACCG**TCACTTGCTAAAATCACTGTCTAC |
| Sequence verification primers for 3Ω1 construct | | Forward | GATGAAAAAGCCCTAAAATTGGAG |
| | | Reverse | ATTATTCACAAATGAGAAACAGAATGG |
| *The In-Fusion cloning overhangs for 3Ω1 cloning are indicated in **bold**. | | | |

| Gene | Plasmid | Primer direction | Sequence (5'-3') |
|---|---|---|---|
| *Escherichia coli* expression using pOPINF constructs (In-Fusion cloning): N-terminal $His_6$-tag | | | |
| CrSGD | pOPINF | Forward | **AAGTTCTGTTTCAGGGCCCG**GGATCTAAAGATGATCAGTC |
| | | Reverse | **ATGGTCTAGAAAGCTTTA**GTATTTTTGCTTCTTGACTAACT |
| CrRedox1 | pOPINF | Forward | **AAGTTCTGTTTCAGGGCCCG**GCTGATCGCGTGAAGA |
| | | Reverse | **ATGGTCTAGAAAGCTTTA**GACAGCTACTGTTGCAT |
| CrRedox2 | pOPINF | Forward | **AAGTTCTGTTTCAGGGCCCG**GAAAAGCAAGTTGAGATCC |
| | | Reverse | **ATGGTCTAGAAAGCTTTA**CAAGTCTCCATCCCAAA |
| CrSAT | pOPINF | Forward | **AAGTTCTGTTTCAGGGCCCG**GCACCCCAGATGCA |
| | | Reverse | **ATGGTCTAGAAAGCTTTA**ATTGCTAAAATCAGTGTCCA |
| TiSGD | pOPINF | Forward | **AAGTTCTGTTTCAGGGCCCG**CATTCGTATTCAACAACTATGG |
| | | Reverse | **ATGGTCTAGAAAGCTTTA**AAAGACCATATTATTTGTGCCTG |
| TiRedox1 | pOPINF | Forward | **AAGTTCTGTTTCAGGGCCCG**GCTGATCGCGTCAA |
| | | Reverse | **ATGGTCTAGAAAGCTTTA**GGCAGCGCCCA |
| TiRedox2 | pOPINF | Forward | **AAGTTCTGTTTCAGGGCCCG**GAAAGGGAAGTTCAAATCC |
| | | Reverse | **ATGGTCTAGAAAGCTTTA**CAAATCTCCATCCCAGAGT |
| TiSAT | pOPINF | Forward | **AAGTTCTGTTTCAGGGCCCG**ACTTCCCAGATGGATGTAG |
| | | Reverse | **ATGGTCTAGAAAGCTTTA**CTTGCTAAAATCACTGTCTAC |
| Sequence verification primers for pOPINF construct | | Forward | TAATACGACTCACTATAGGG |
| | | Reverse | TAGCCAGAAGTCAGATGCT |
| *Escherichia coli* expression using pET-28a(+) constructs (In-Fusion cloning): C-terminal $His_6$-tag | | | |
| CrGS | pET28a(+) | Forward | **AGGAGATATACCATGG**ATGGCCGGAGAAACA |
| | | Reverse | **GGTGGTGGTGCTCGAG**TTCCTCAAATTTCAATGTATTTCC |
| TiGS | pET28a(+) | Forward | **AGGAGATATACCATGG**ATGGCTGCAGAAACAG |
| | | Reverse | **GGTGGTGGTGCTCGAG**TTCATCATATTTCAATGTATTGCCAA |
| Sequence verification primers for pET28a construct | | Forward | TAATACGACTCACTATAGGG |
| | | Reverse | GCTAGTTATTGCTCAGCGG |
| *In-Fusion cloning overhangs for pOPINF and pET-28a(+) cloning are indicated in **bold**. | | | |

| Gene | Plasmid | Primer direction | Sequence (5'-3') |
|---|---|---|---|
| *Saccharomyces cerevisiae* expression using pESC-HIS (MCS2) constructs (In-Fusion cloning) | | | |
| CrGO | pESC-HIS | Forward | **GAGAAAAAACCCCGGATCC**ATGGAGTTTTCTTTCTCCTCAC |
| | | Reverse | **ACTTCTGTTCCATGTCGAC**CTAATCGTTAACAAGATGAGGAA |
| CrSBE | pESC-HIS | Forward | **GAGAAAAAACCCCGGATCC**ATGGATGAGATGATGAACTTCTC |
| | | Reverse | **ACTTCTGTTCCATGTCGAC**TTAATTTCCTTCAACTGCTGATGA |
| TiGO | pESC-HIS | Forward | **GAGAAAAAACCCCGGATCC**ATGGAGCTCTCTTTCTCCTC |
| | | Reverse | **ACTTCTGTTCCATGTCGAC**CTATTCGTTGGCAAGATGTG |
| TiSBE | pESC-HIS | Forward | **GAGAAAAAACCCCGGATCC**ATGGAGATGATGAACTTCTCTTTCA |
| | | Reverse | **ACTTCTGTTCCATGTCGAC**TTAATTTCCTGAAATGATGTTGGGAATTA |
| Sequence verification primers pESC-HIS construct | | Forward | ATGATTTTTGATCTATTAACAGATA |
| | | Reverse | GTATAATGTTACATGCGTACAC |
| *In-Fusion cloning overhangs for pESC-HIS (MCS2) cloning are indicated in **bold**. | | | |

**Table S3**. Primer sequences used for the generation CDS parts for the assembly of multigene constructs using GoldenBraid 2.0 cloning in this study.

| Gene | Level 1 CDS | Primer direction | Sequence (5'-3') |
|---|---|---|---|
| *Nicotiana benthamiana* multigene constructs | | | |
| Level 1 CDS fragment for 3α1 or 3α2 GoldenBraid 2.0 assembly | | | |
| Domestication of TiRedox2 for pUPD entry | Frag. A | Forward | GCGCCGTCTCGCTCGAATGGAAAGGGAAGTTCAAATCC |
| | | Reverse | GCGCCGTCTCGGGGACCATGCGGTGACATGA |
| | Frag. B | Forward | GCGCCGTCTCGTCCCCTCTGTTATCTTATGGT |
| | | Reverse | GCGCCGTCTCGCTCGAAGCTTACAAATCTCCATCCCAGAG |
| | Full length | Forward | GCGCCGTCTCGCTCGAATGGAAAGGGAAGTTCAAATCC |
| | | Reverse | GCGCCGTCTCGCTCGAAGCTTACAAATCTCCATCCCAGAG |
| Domestication of TiSBE for pUPD entry | Frag. A | Forward | GCGCCGTCTCGCTCGAATGGAGATGATGAACTTCTCTTTC |
| | | Reverse | GCGCCGTCTCGGGATCTCTTCCCACTGCCCA |
| | Frag. B | Forward | GCGCCGTCTCGATCCTGAACTTTGGGACGATG |
| | | Reverse | GCGCCGTCTCGCTCGAAGCTTAATTTCCTGAAATGATGTTGGG |
| | Full length | Forward | GCGCCGTCTCGCTCGAATGGAGATGATGAACTTCTCTTTC |
| | | Reverse | GCGCCGTCTCGCTCGAAGCTTAATTTCCTGAAATGATGTTGGG |
| TiGO | Full length | Forward | **TTCAGAGGTCTCTA**ATGGAGCTCTCTTTCTCCTC |
| | | Reverse | **AGCGTGGGTCTCGAAGC**CTATTCGTTGGCAAGATGTG |
| TiRedox1 | Full length | Forward | **TTCAGAGGTCTCTA**ATGGCTGATCGCGTCAAGA |
| | | Reverse | **AGCGTGGGTCTCGAAGC**CTAGGCAGCGCCCATTG |
| TiSAT | Full length | Forward | **TTCAGAGGTCTCTA**ATGACTTCCCAGATGGATGT |
| | | Reverse | **AGCGTGGGTCTCGAAGC**TCACTTGCTAAAATCACTGTCTA |
| P19 | Full length | Forward | **TTCAGAGGTCTCTA**ATGGAACGAGCTATACAAGGAAA |
| | | Reverse | **AGCGTGGGTCTCGAAGC**TCACTCGCTTTCTTTTTCGAAGG |
| Plasmid validation | P19 | Forward | TATTTAAGAGATATCTCAGATACG |
| | | Reverse | CTGGTCGAAACCGAAAAATC |

| | | | |
|---|---|---|---|
| | TiGO | Forward | CAGGTTGGAGGATCTGTTCCC |
| | | Reverse | TGCCTCAAGAGAACGCTGAG |
| | TiRedox1 | Forward | CTTCCTCTCGATGCTGGTGC |
| | | Reverse | AGTCCAAAAGCCTTGGCGAA |
| | TiRedox2 | Forward | GTAAGAGTGAGGCAGGGTGC |
| | | Reverse | AACTGCAGGAGGAATGGTGG |
| | TiSAT | Forward | ACCCATGTTGTGCTGCCTAA |
| | | Reverse | GGATTCTTCACCTCCGAGGC |
| Overhangs designed for non-domesticated sequences 3α1 or 3α2 GoldenBraid assembly | | | |

**Table S4**. Primer sequences used for CDS (domesticated) amplification to generate C-terminal eYFP fusion constructs for subcellular localization with GoldenBraid 2.0 in this study.

| **Gene** | **Plasmid** | **Primer direction** | **Sequence (5'-3')** |
|---|---|---|---|
| *Nicotiana benthamiana* single transcriptional unit constructs (3$\alpha$1, GoldenBraid 2.0) | | | |
| TiSGD | 3$\alpha$1 | Forward | **TTCAGAGGTCTCTA**ATGCATTCGTATTCAACAACTATG |
| | | Reverse | **AGCGTGGGTCTCGCTGC***AGAAGAACCAGAACC*AAAGACCATATTATTTGTGCCTG |
| TiGS | 3$\alpha$1 | Forward | **TTCAGAGGTCTCTA**ATGGCTGCAGAAACAGC |
| | | Reverse | **AGCGTGGGTCTCGCTGC***AGAAGAACCAGAACC*TTCATCATATTTCAATGTATTGCCAAT |
| TiSBE | 3$\alpha$1 | Forward | **TTCAGAGGTCTCTA**ATGGAGATGATGAACTTCTCTTTCA |
| | | Reverse | **AGCGTGGGTCTCGCTGC***AGAAGAACCAGAACC*ATTTCCTGAAATGATGTTGGGAATTA |
| TiGO | 3$\alpha$1 | Forward | **TTCAGAGGTCTCTA**ATGGAGCTCTCTTTCTCCT |
| | | Reverse | **AGCGTGGGTCTCGCTGC***AGAAGAACCAGAACC*TTCGTTGGCAAGATGTG |
| TiRedox1 | 3$\alpha$1 | Forward | **TTCAGAGGTCTCTA**ATGGCTGATCGCGT |
| | | Reverse | **AGCGTGGGTCTCGCTGC***AGAAGAACCAGAACC*GGCAGCGCCCA |
| TiRedox2 | 3$\alpha$1 | Forward | **TTCAGAGGTCTCTA**ATGGAAAGGGAAGTTCAAATC |
| | | Reverse | **AGCGTGGGTCTCGCTGC***AGAAGAACCAGAACC*CAAATCTACATCCCAGAGTTC |
| TiSAT | 3$\alpha$1 | Forward | **TTCAGAGGTCTCTA**ATGACTTCCCAGATGGATGTA |
| | | Reverse | **AGCGTGGGTCTCGCTGC***AGAAGAACCAGAACC*CTTGCTAAAATCACTGTCTACAAG |
| Sequence verification primers 3$\alpha$1 construct | | Forward | GATGAAAAAGCCCTAAAATTGGAG |
| | | Reverse | ATTATTCACAAATGAGAAACAGAATGG |
| *GoldenBraid 2.0 compatible cloning overhangs for 3$\alpha$1 transcriptional unit assembly are indicated in **bold**.<br>**GSGSS linker sequences are indicated by *italics*.<br>Transcriptional unit = Ubq10Promoter+5UTR+Gene+C-terminal-YFP+3UTR+Ubq10Terminator | | | |

**Table S5**. Primer sequences used for qPCR analysis in this study.

| **Target gene** | **Primer direction** | **Sequence (5'-3')** |
|---|---|---|
| CrSGD | Forward | ATTGATGCTCGGGAAAGGGG |
| | Reverse | AGACGGCTTCCTACAAGAGC |
| CrGS | Forward | ATGGCCGGAGAAACAACCAA |
| | Reverse | CCGTTCACCAGGGACTCTTC |
| TiSGD | Forward | TGGTGGCTTCTTGAGTTCCC |
| | Reverse | GTTCTGTGCCTGGATCACCT |
| TiGS | Forward | ATTACTGCCCAGAGCCCAAC |
| | Reverse | GTCCTGTGGGAAGTTCTCGG |
| *Nicotiana benthamiana* Elongation Factor 1 alpha (NbEF1$\alpha$) – Reference gene (PQ008965.1) | Forward | AGCGTGGTTATGTTGCCTCA |
| | Reverse | GACCTGGGAGGTGAAACTGG |

**Supporting NMR Tables**

**Table NMR S1**. NMR shifts of stemmadenine as formate salt.

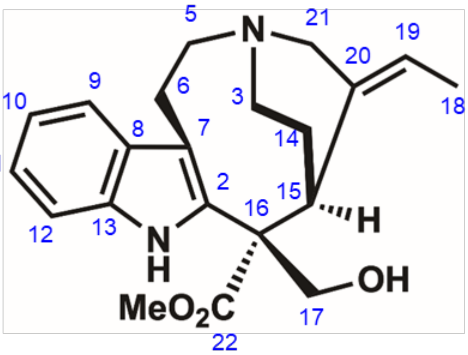

**stemmadenine (formate salt)**
**(MeOH-*d*3, 700 MHz, 298 K)**

| pos. | δH | mult. | JHH | δC |
|---|---|---|---|---|
| 1(NH) | 10.14 | - | - | - |
| 2 | - | - | - | 134.9 |
| 3α | 3.10 | *ddd* | 13.6/13.4/6.4 | 47.2 |
| 3β | 3.27 | *dd* | 13.4/6.1 | 47.2 |
| 5α | 3.47 | *m* | - | 56.8 |
| 5β | 3.38 | *m*** | - | 56.8 |
| 6α | 3.54 | *m* | - | 24.4 |
| 6β | 3.38 | *m*** | - | 24.4 |
| 7 | - | - | - | 110.9 |
| 8 | - | - | - | 128.2 |
| 9 | 7.53 | *d* | 7.9 | 118.6 |
| 10 | 7.07 | *dd* | 7.9/7.6 | 120.4 |
| 11 | 7.13 | *dd* | 8.1/7.6 | 123.0 |
| 12 | 7.39 | *d* | 8.1 | 112.5 |
| 13 | - | - | - | 137.0 |
| 14α | 2.36 | *ddd* | 16.1/12.9/6.6 | 26.1 |
| 14β | 2.61 | *m* | - | 26.1 |
| 15α | 3.80 | *dd* | 12.7/3.0 | 36.5 |
| 16 | - | - | - | 61.7 |
| 17a | 4.35 | *s* | - | 69.4 |
| 17b | 4.35 | *s* | - | 69.4 |
| 18 | 1.77 | *dd* | 6.9/2.0 | 14.3 |
| 19 | 5.57 | *q* | 6.9 | 130.9 |
| 20 | - | - | - | 129.0 |
| 21α | 3.37 | *m*** | - | 55.1 |
| 21β | 2.91 | *brd* | 14.8 | 55.1 |
| 22 | - | - | - | 174.2 |
| OMe | 3.80 | *s* | - | 53.0 |

** overlapped signals *J* unresolved, 700 MHz in MeOH-*d*3

**Table NMR S2**. NMR shifts of 16(*R*)-19*E*-isositirikine as acetate salt.

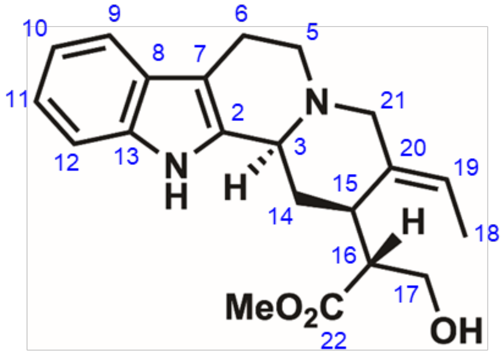

**16(*R*)-19*E*-isositirikine (acetate salt)**
**($CDCl_3$, 700 MHz, 298 K)**

| pos. | δH | mult. | JHH | δC |
|---|---|---|---|---|
| 1(NH) | 9.00 | - | - | - |
| 2 | - | - | - | 131.5 |
| 3α | 4.55 | *m* | - | 52.7 |
| 5α | 3.24 | *ddd* | 13.2/11.3/5.2 | 50.4 |
| 5β | 3.34 | *ddd* | 13.2/5.8/2.1 | 50.4 |
| 6α | 2.76 | *m* | - | 17.4 |
| 6β | 2.98 | *m* | - | 17.4 |
| 7 | - | - | - | 107.2 |
| 8 | - | - | - | 127.3 |
| 9 | 7.44 | *d* | 7.8 | 118.3 |
| 10 | 7.09 | *dd* | 7.8/7.4 | 120.0 |
| 11 | 7.16 | *dd* | 8.0/7.4 | 122.3 |
| 12 | 7.37 | *d* | 8.0 | 111.7 |
| 13 | - | - | - | 136.7 |
| 14α | 2.30 | *m*** | - | 29.2 |
| 14β | 2.31 | *m*** | - | 29.2 |
| 15α | 3.10 | *ddd* | 11.2/4.7/4.7 | 32.5 |
| 16 | 2.41 | *ddd* | 11.2/7.8/5.0 | 49.3 |
| 17a | 3.54 | *dd* | 11.5/7.8 | 62.1 |
| 17b | 3.47 | *dd* | 11.5/5.0 | 62.1 |
| 18 | 1.64 | *brd* | 6.7 | 13.7 |
| 19 | 5.70 | *q* | - | 126.8 |
| 20 | - | - | - | 131.4 |
| 21α | 3.21 | *d* | 12.9 | 51.6 |
| 21β | 3.56 | *d* | 12.9 | 51.6 |
| 22 | - | - | - | 175.3 |
| OMe | 3.79 | *s* | - | 52.6 |

** overlapped signals *J* unresolved, 700 MHz in $CDCl_3$

**Table NMR S3**. NMR shifts of 16(*S*)-19*E*-isositirikine as acetate salt.

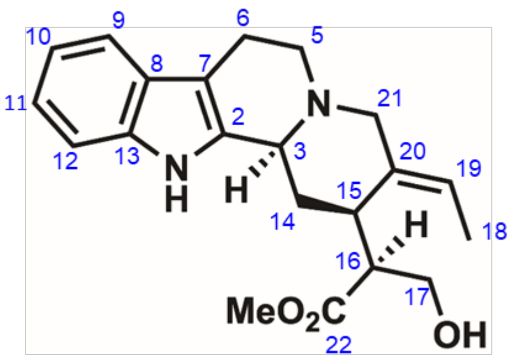


| 16(*R*)-19*E*-isositirikine (acetate salt) (CDCl3, 700 MHz, 298 K) | | | | |
|---|---|---|---|---|
| pos. | δH | mult. | JHH | δC |
| 1(NH) | 9.76 | - | - | - |
| 2 | - | - | - | 130.6 |
| 3α | 4.65 | *m* | - | 52.8 |
| 5α | 3.16 | *ddd* | 12.2/11.1/5.1 | 50.2 |
| 5β | 3.35 | *m*** | - | 50.2 |
| 6α | 2.77 | *m* | - | 17.4 |
| 6β | 3.00 | *m* | - | 17.4 |
| 7 | - | - | - | 106.5 |
| 8 | - | - | - | 127.0 |
| 9 | 7.41 | *d* | 7.7 | 118.4 |
| 10 | 7.07 | *brdd* | 7.7/7.5 | 120.0 |
| 11 | 7.13 | *brdd* | 8.1/7.5 | 122.4 |
| 12 | 7.31 | *d* | 8.1 | 111.6 |
| 13 | - | - | - | 136.9 |
| 14α | 2.37 | *m*** | - | 28.5 |
| 14β | 2.59 | *m* | - | 28.5 |
| 15α | 3.25 | *m* | - | 32.8 |
| 16 | 2.35 | *m*** | - | 48.2 |
| 17a | 3.90 | *m* | - | 62.1 |
| 17b | 3.90 | *m* | - | 62.1 |
| 18 | 1.57 | *dd* | 6.8/1.1 | 13.4 |
| 19 | 5.64 | *q* | 6.8 | 126.9 |
| 20 | - | - | - | 131.1 |
| 21α | 3.38 | *d* | 12.5 | 52.3 |
| 21β | 3.82 | *d* | 12.5 | 52.3 |
| 22 | - | - | - | 174.5 |
| OMe | 3.47 | *s* | - | 51.8 |

** overlapped signals *J* unresolved, 700 MHz in $CDCl_3$

**Table NMR S4**. NMR shifts of 16(*R*)-17D-19*E*-isositirikine as acetate salt.

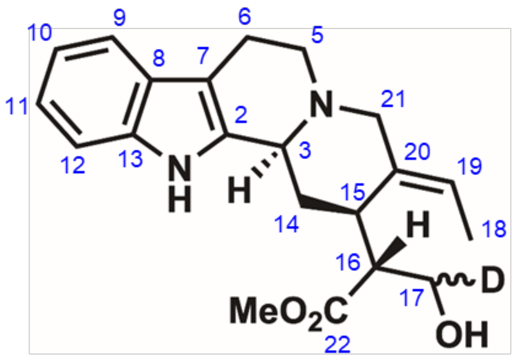


| 16(*R*)-17D-19*E*-isositirikine (acetate salt) (CDCl3, 700 MHz, 298 K) | | | | |
|---|---|---|---|---|
| pos. | δH | mult. | JHH | δC |
| 1(NH) | 9.31 | - | - | - |
| 2 | - | - | - | 130.8 |
| 3α | 4.61 | *m* | - | 52.6 |
| 5α | 3.26 | *m* | - | 49.9 |
| 5β | 3.31 | *m*** | - | 49.9 |
| 6α | 2.78 | *m* | - | 17.4 |
| 6β | 2.94 | *m* | - | 17.4 |
| 7 | - | - | - | 106.7 |
| 8 | - | - | - | 127.0 |
| 9 | 7.41 | *d* | 7.9 | 118.3 |
| 10 | 7.07 | *brdd* | 7.9/7.5 | 119.9 |
| 11 | 7.14 | *brdd* | 8.0/7.5 | 122.4 |
| 12 | 7.35 | *brd* | 8.0 | 111.7 |
| 13 | - | - | - | 136.7 |
| 14α | 2.35 | *ddd* | 15.3/5.7/5.7 | 28.7 |
| 14β | 2.29 | *m* | - | 28.7 |
| 15α | 3.10 | *m* | - | 32.4 |
| 16 | 2.40 | *m* | - | 49.1 |
| 17a* | 3.52 | *d* | 7.5 | 61.5 |
| 17b* | 3.46 | *d* | 5.1 | 61.5 |
| 18 | 1.61 | *brd* | 6.8 | 13.7 |
| 19 | 5.71 | *q* | 6.8 | 128.0 |
| 20 | - | - | - | 130.4 |
| 21α | 3.35 | *d* | 12.3 | 51.8 |
| 21β | 3.55 | *d* | 12.3 | 51.8 |
| 22 | - | - | - | 175.0 |
| OMe | 3.75 | *s* | - | 52.5 |

** overlapped signals *J* unresolved, *0.5H, 700 MHz in $CDCl_3$

**Table NMR S5**. NMR shifts of 16(*S*)-17D-19*E*-isositirikine as acetate salt.

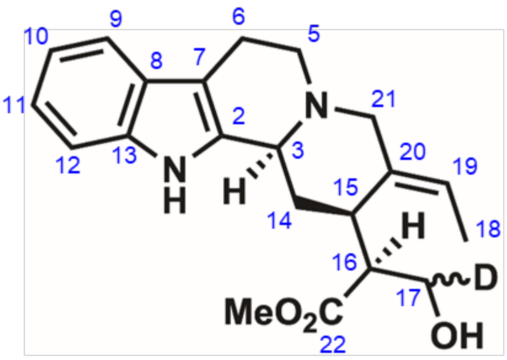


| 16(*R*)-17D-19*E*-isositirikine (acetate salt) (CDCl3, 700 MHz, 298 K) | | | | |
|---|---|---|---|---|
| pos. | δH | mult. | JHH | δC |
| 1(NH) | 10.29 | - | - | - |
| 2 | - | - | - | 130.1 |
| 3α | 4.71 | *m* | - | 52.8 |
| 5α | 3.17 | *m*** | - | 50.1 |
| 5β | 3.35 | *m* | - | 50.1 |
| 6α | 2.76 | *m* | - | 17.1 |
| 6β | 3.00 | *m* | - | 17.1 |
| 7 | - | - | - | 106.0 |
| 8 | - | - | - | 126.8 |
| 9 | 7.38 | *d* | 7.8 | 118.3 |
| 10 | 7.05 | *brdd* | 7.8/7.5 | 119.9 |
| 11 | 7.11 | *brdd* | 8.1/7.5 | 122.4 |
| 12 | 7.30 | *brd* | 8.1 | 111.7 |
| 13 | - | - | - | 137.0 |
| 14α | 2.37 | *ddd* | 15.1/5.9/5.9 | 28.2 |
| 14β | 2.64 | *m* | - | 28.2 |
| 15α | 3.19 | *m* | - | 32.6 |
| 16 | 2.29 | *m* | - | 48.1 |
| 17a* | 3.87 | *m* | - | 61.6 |
| 17b* | 3.87 | *m* | - | 61.6 |
| 18 | 1.55 | *brd* | 6.7 | 13.4 |
| 19 | 5.64 | *q* | 6.7 | 127.4 |
| 20 | - | - | - | 130.7 |
| 21α | 3.40 | *d* | 12.3 | 51.7 |
| 21β | 3.82 | *d* | 12.3 | 51.7 |
| 22 | - | - | - | 174.4 |
| OMe | 3.44 | *s* | - | 51.7 |
| ** overlapped signals *J* unresolved, *0.5H, 700 MHz in CDCl3 | | | | |

**Table NMR S6**. NMR shifts of condylocarpine as formate salt.

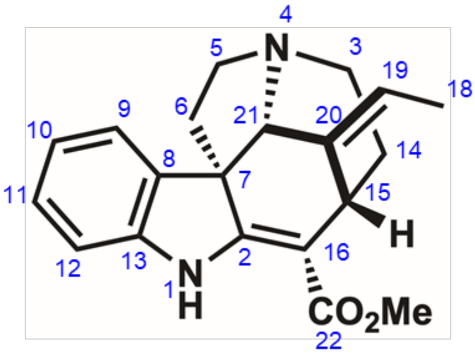


| condylocarpine (formate salt) (MeOH-*d3*, 700 MHz, 298 K) | | | | |
|---|---|---|---|---|
| pos. | δH | mult. | JHH | δC |
| 1(NH) | 9.13 | - | - | - |
| 2 | - | - | - | 168.7 |
| 3α | 3.05 | *m* | - | 47.2 |
| 3β | 3.22 | *m*** | - | 47.2 |
| 5α | 3.23 | *m*** | - | 52.9 |
| 5β | 3.40 | *m* | - | 52.9 |
| 6α | 2.87 | *m* | - | 44.8 |
| 6β | 2.08 | *m* | - | 44.8 |
| 7 | - | - | - | 60.0 |
| 8 | - | - | - | 134.6 |
| 9 | 7.32 | *d* | 7.3 | 121.1 |
| 10 | 6.91 | *dd* | 7.6/7.3 | 122.2 |
| 11 | 7.15 | *dd* | 7.8/7.6 | 129.5 |
| 12 | 6.92 | *d* | 7.8 | 111.3 |
| 13 | - | - | - | 146.0 |
| 14a | 2.01 | *m*** | - | 27.9 |
| 14b | 2.01 | *m*** | - | 27.9 |
| 15 | 4.07 | *dd* | 4.7/4.7 | 29.4 |
| 16 | - | - | - | 101.6 |
| 18 | 1.64 | *d* | 6.7 | 13.0 |
| 19 | 5.49 | *q* | 6.7 | 122.0 |
| 20 | - | - | - | 135.2 |
| 21β | 4.51 | *brs* | - | 69.6 |
| 22 | - | - | - | 168.5 |
| OMe | 3.80 | *s* | - | 51.7 |
| ** overlapped signals *J* unresolved, 700 MHz in MeOH-*d3* | | | | |

### Supporting Figures

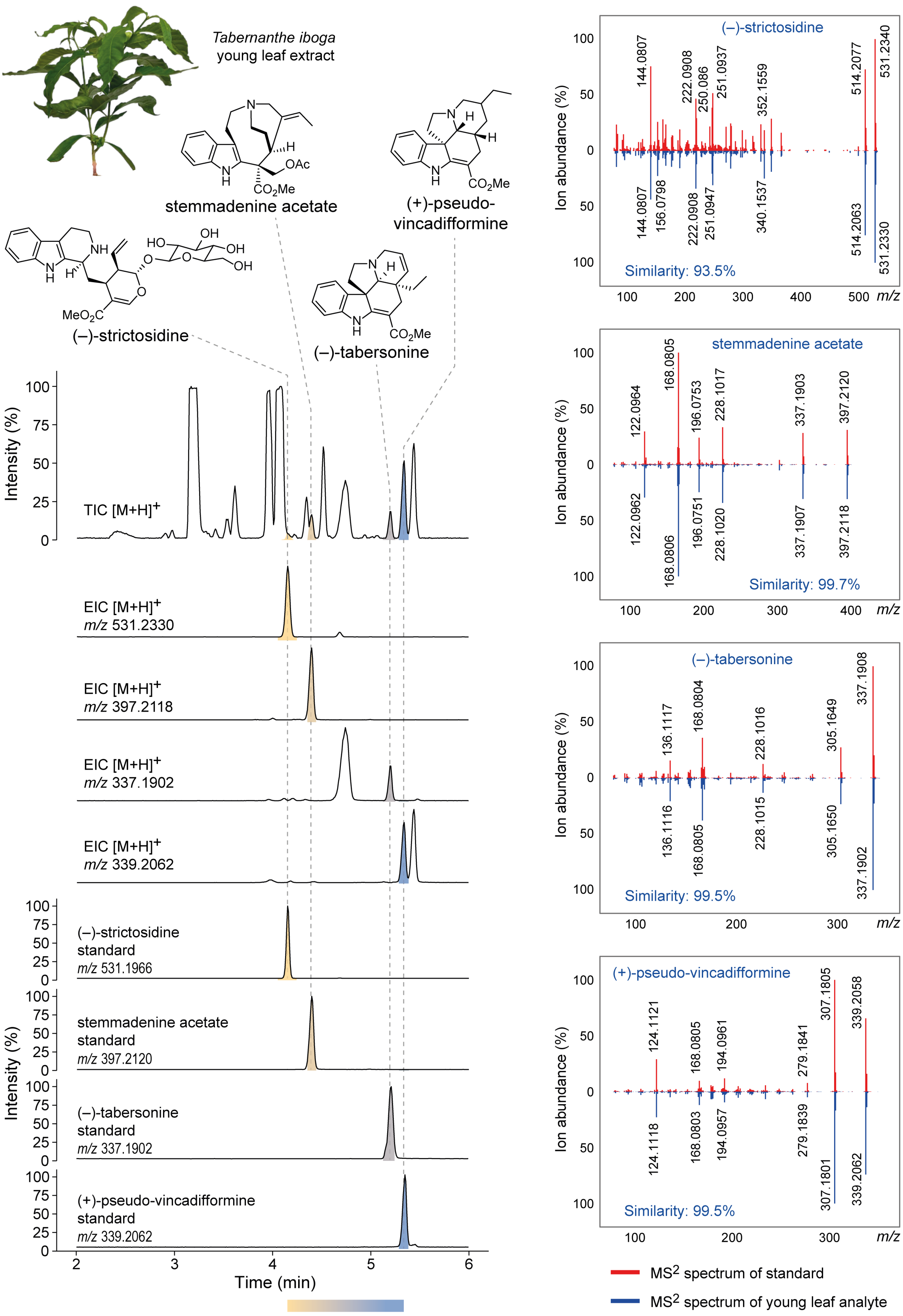


Abbreviation: TIC = Total ion chromatogram, EIC = Extracted ion chromatogram

**Figure S1. Monoterpene indole alkaloids (MIAs) detected in the young leaf of *Tabernanthe iboga*.** LC-MS chromatograms showing the presence of early-pathway MIAs in young leaves of *T. iboga*. $MS^2$ spectra of authentic standards and corresponding analytes are shown.

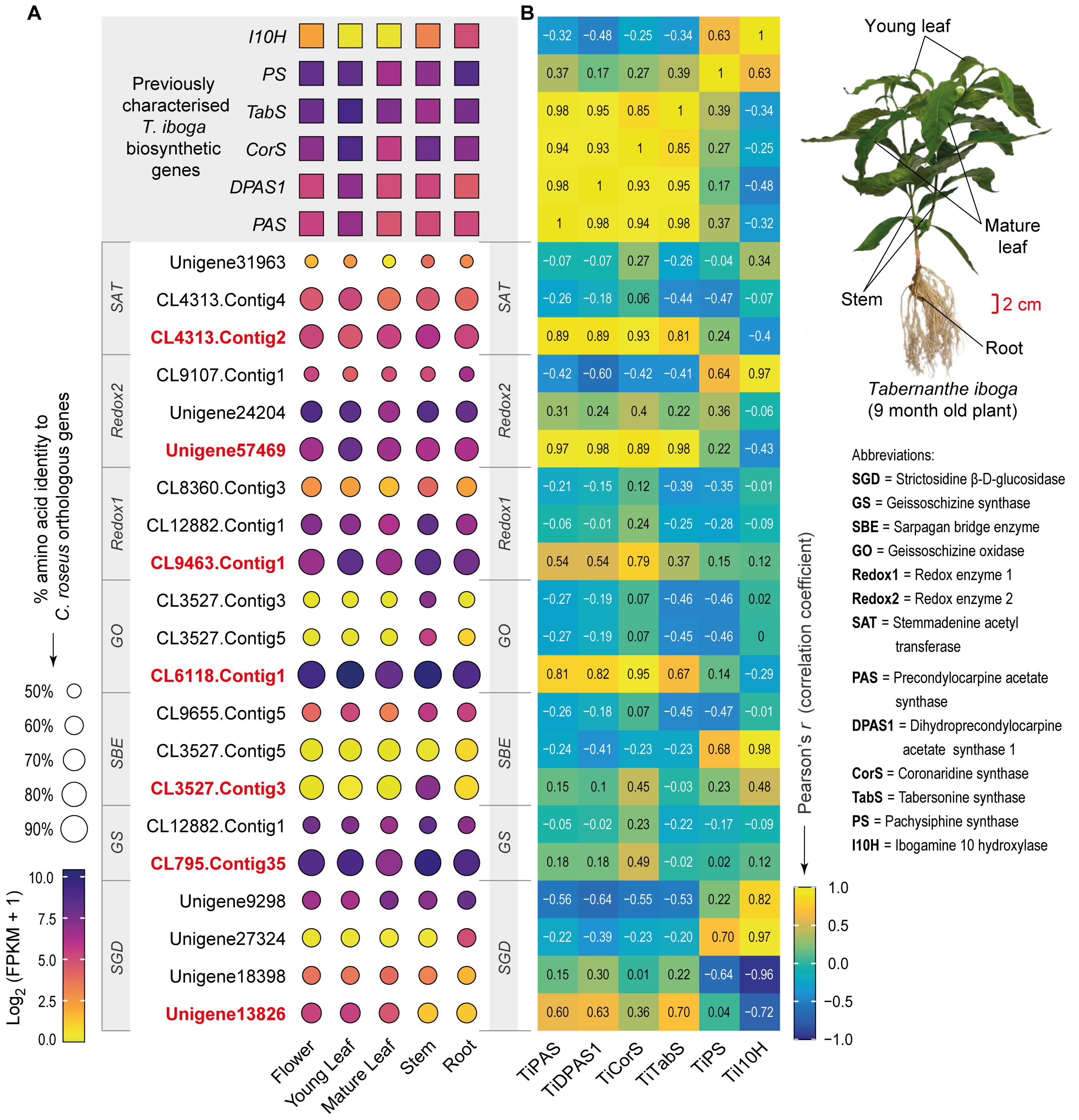


**Figure S2. Early pathway candidate identification and selection.** (**A**) Expression profiles of *Tabernanthe iboga* genes orthologous to functionally characterised *Catharanthus roseus* biosynthetic genes (SGD to SAT) across various tissues. The heatmap indicates transcript abundance in bulk tissue samples across five tissues of the *T. iboga* plant. At the same time, the size of each dot in the dot plot reflects the % amino acid identity to the corresponding *C. roseus* enzyme. (**B**) Pearson correlation coefficients of *T. iboga* early pathway candidates (SGD – SAT) to previously characterised *T. iboga* genes are also shown. The *T. iboga* enzymes PAS (1), DPAS1 (1), CorS (1), TabS (1), PS (2), and I10H (1) were used as bait genes to identify candidates by co-expression, with positively correlated genes prioritised (highlighted in red) for further functional testing as putative biosynthetic genes of the stemmadenine acetate biosynthetic pathway.

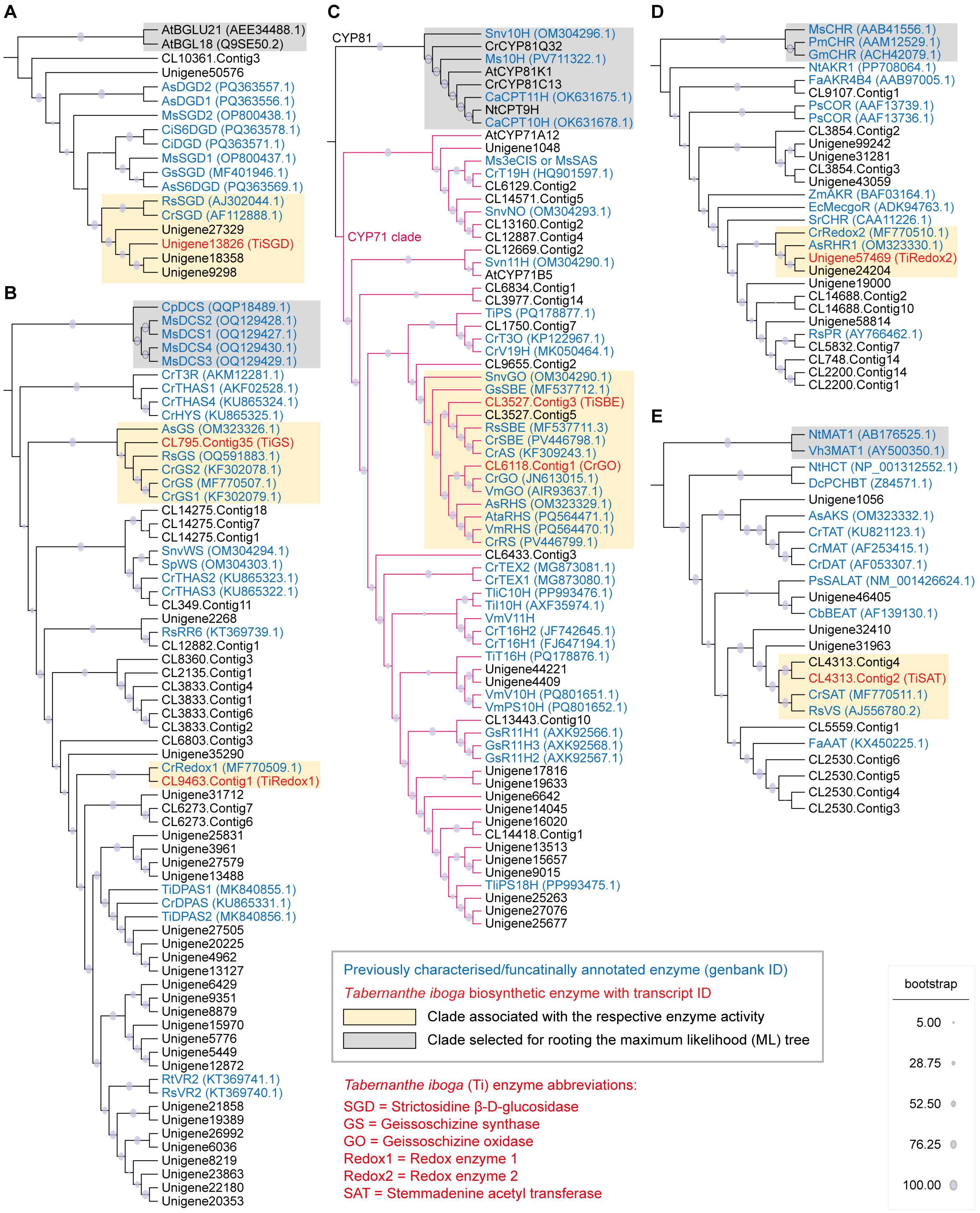


**Figure S3. Phylogenetic relationtionsips of stemmadenine biosynthetic enzymes with previously characterized plant natural product biosynthetic enzymes.** Maximum likelihood phylogenetic trees were generated to examine the relationships between stemmadenine biosynthetic enzymes and their plant homologs. (**A**) Relationship of SGD with other characterized and annotated plant SGD enzymes. (**B**) Relationship of GS and Redox1 with medium chain alcohol dehydrogenase/reductase (MDR) family enzymes involved in monoterpene indole alkaloid (MIA) biosynthesis. (**C**) Relationship of GO with cytochrome P450 enzymes of the CYP71 family associated with MIA biosynthesis. (**D**) Relationship of Redox 2 with known plant aldo keto reductases (AKRs). (**E**) Relationship of SAT with a subset of plant acetyl transferases involved in plant natural product biosynthesis. All trees shown were inferred using maximum likelihood methods (1000 bootstraps). Trees are rooted at the clades highlighted in grey boxes.

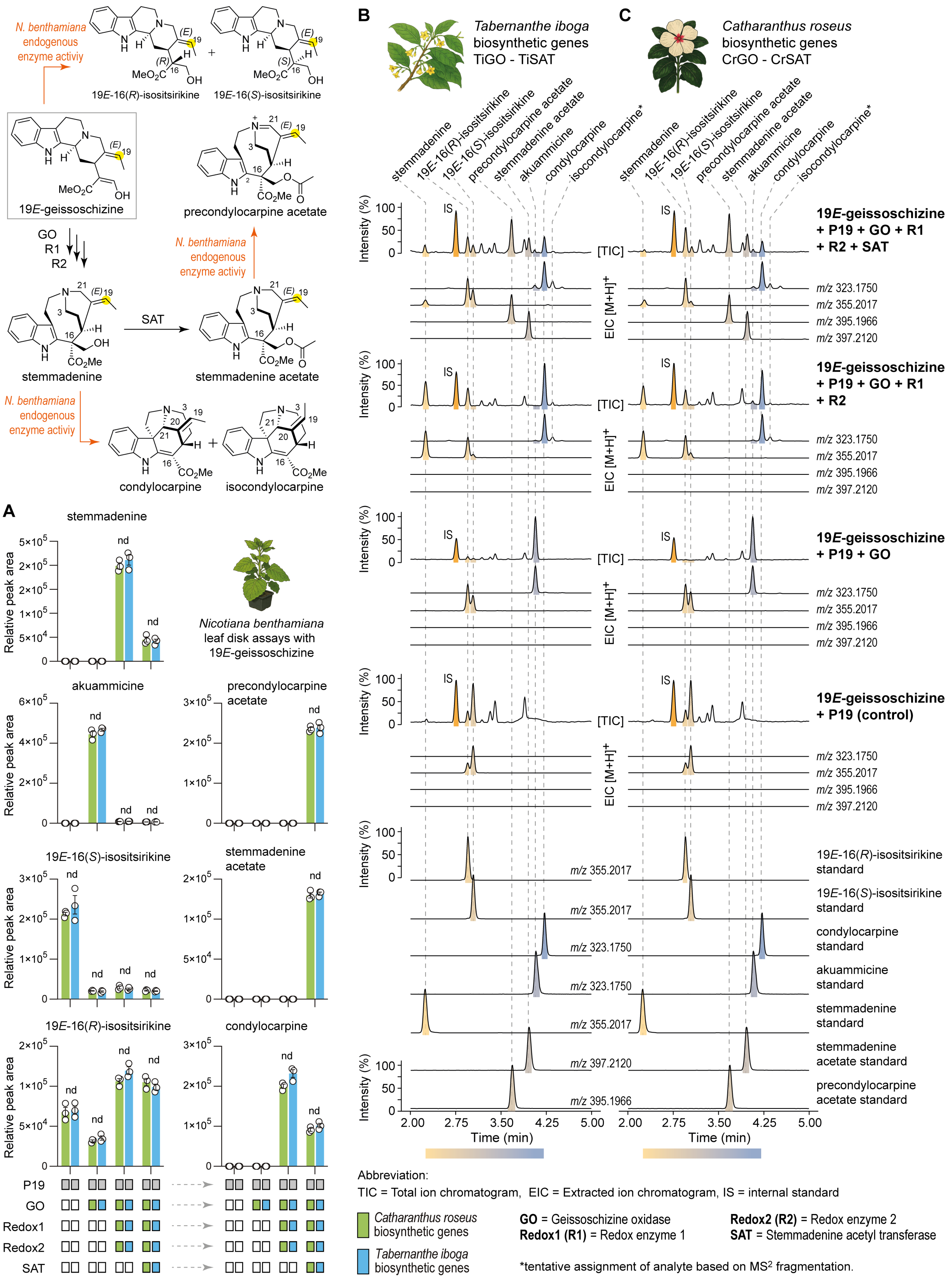

**Figure S4. Heterologous production of stemmadenine acetate in *N. benthamiana* catalyzed by *Tabernanthe iboga* genes. (A)** Stepwise pathway reconstitution of *T. iboga* candidate genes (blue) involved in stemmadenine acetate biosynthesis in *Nicotiana benthamiana*, starting from the substrate 19*E*-geissoschizine, and benchmarked against functionally characterized *Catharanthus roseus* genes (green) (3). Quantitative analysis shows no statistically significant difference in the accumulation of biosynthetic products from GO to SAT between the *T. iboga* and *C. roseus* gene sets (unpaired t-test, $n = 3$ biological replicates, nd = not significant $P > 0.05$). (**B**) Representative LC-MS chromatogram of biosynthetic products generated by stepwise reconstitution of *T. iboga* biosynthetic genes. (**C**) Representative LC-MS chromatogram of products generated using the corresponding *C. roseus* biosynthetic genes. Endogenous *N. benthamiana* metabolic activity affecting monoterpene indole alkaloids (MIAs) is also shown, revealing background modifications that occur during pathway reconstitution in this heterologous host. Relative peak area is normalized to the internal standard (IS). Bar graphs are ploted as the mean ± S.D

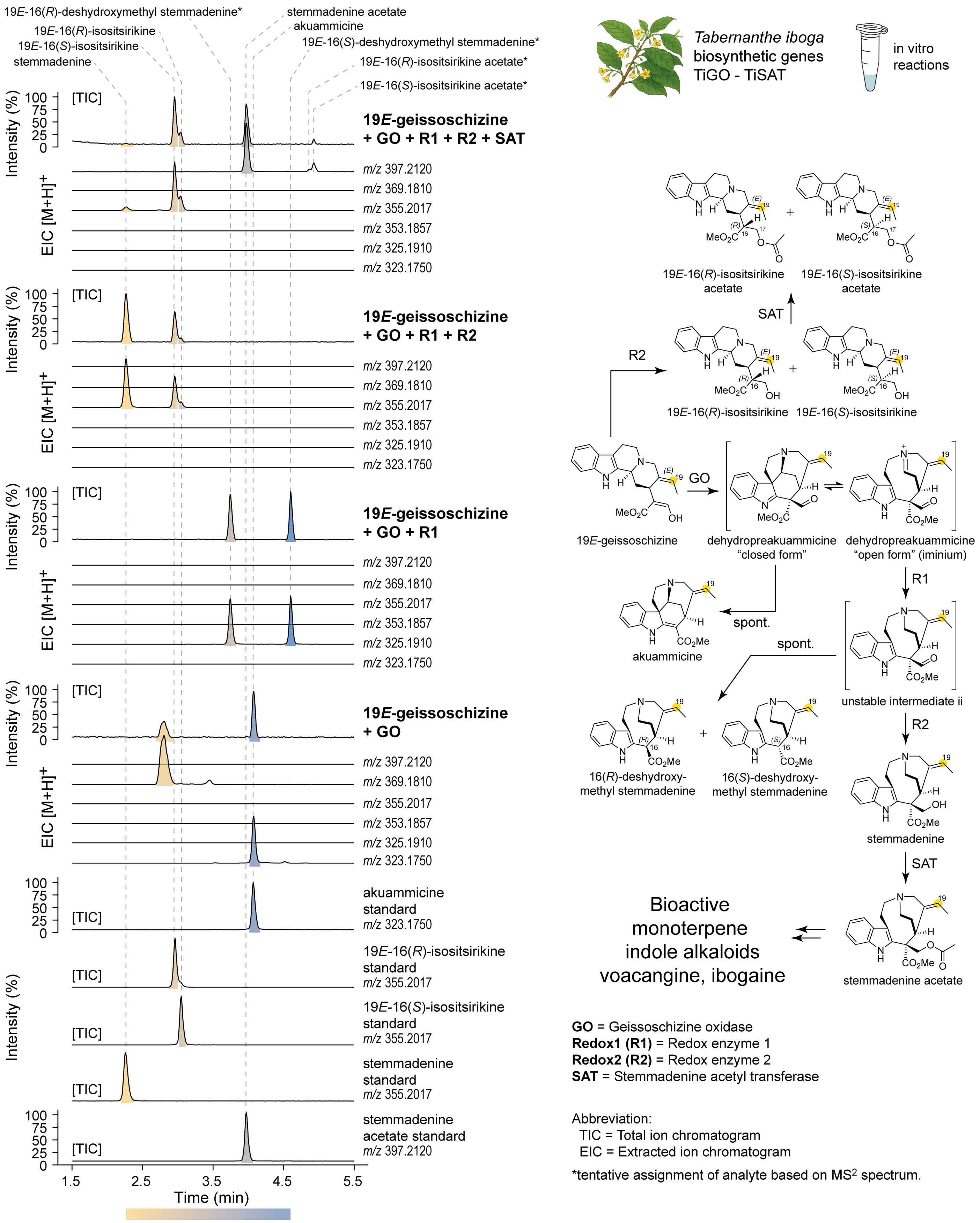


**Figure S5. Stepwise in vitro reconstitution of the stemmadenine acetate biosynthetic pathway from 19*E*-geissoschizine using *Tabernanthe iboga* biosynthetic enzymes.** LC-MS chromatograms showing the sequential enzymatic conversion of 19*E*-geissoschizine to stemmadenine acetate via the action of *T. iboga* GO, Redox 1, Redox2, and SAT. Enzymes were incubated in combination to demonstrate stepwise pathway progression from 19*E*-geissoschizine substrate. Reaction products were identified based on comparison with authentic standards using retention times and $MS^2$ fragmentation spectra.

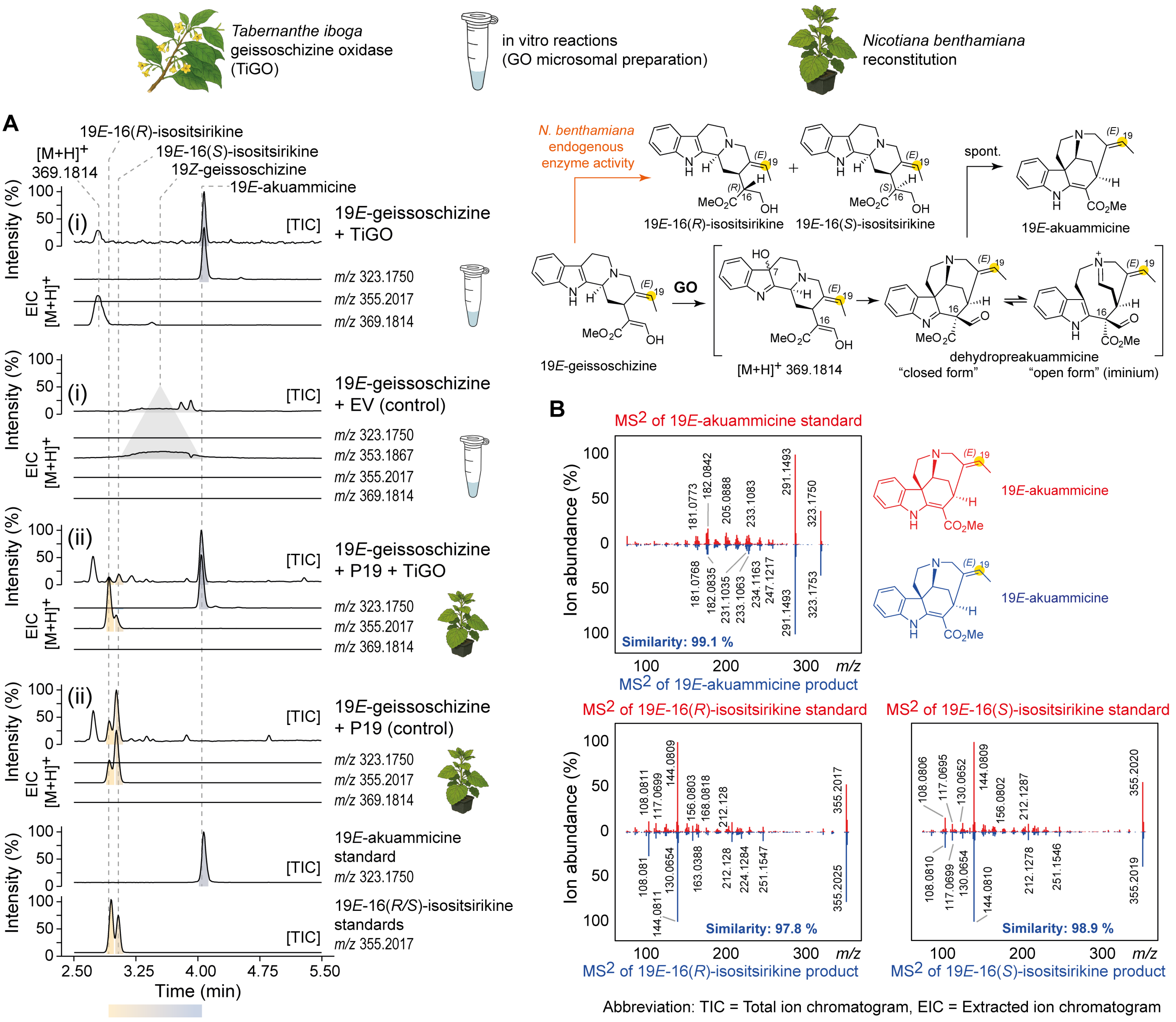


**Figure S6. Functional characterization of *Tabernanthe iboga* geissoschizine oxidase (TiGO) activity with 19*E*-geissoschizine.** (**A**) (i) LC-MS chromatograms of in vitro enzymatic assays using yeast microsomes expressing TiGO incubated with 19*E*-geissoschizine as substrate. (ii) LC-MS chromatograms of TiGO transiently expressed in *N. benthamiana* with the P19 viral silencing suppressor, assayed in leaf discs supplied with 19*E*-geissoschizine. In both systems, TiGO catalysed the oxidation of 19*E*-geissoschizine to produce the thermodynamic product akuammicine (19*E*-akuammicine) in the absence of downstream biosynthetic enzymes. The endogenous activity of *N. benthamiana* metabolic activity led to the reduction of 19*E*-geissoschizine to 19*E*-isositsirikine isomers. (**B**) $MS^2$ spectrum of TiGO reaction products detected in both in vivo and in vitro assays, compared with spectra of authentic standards.

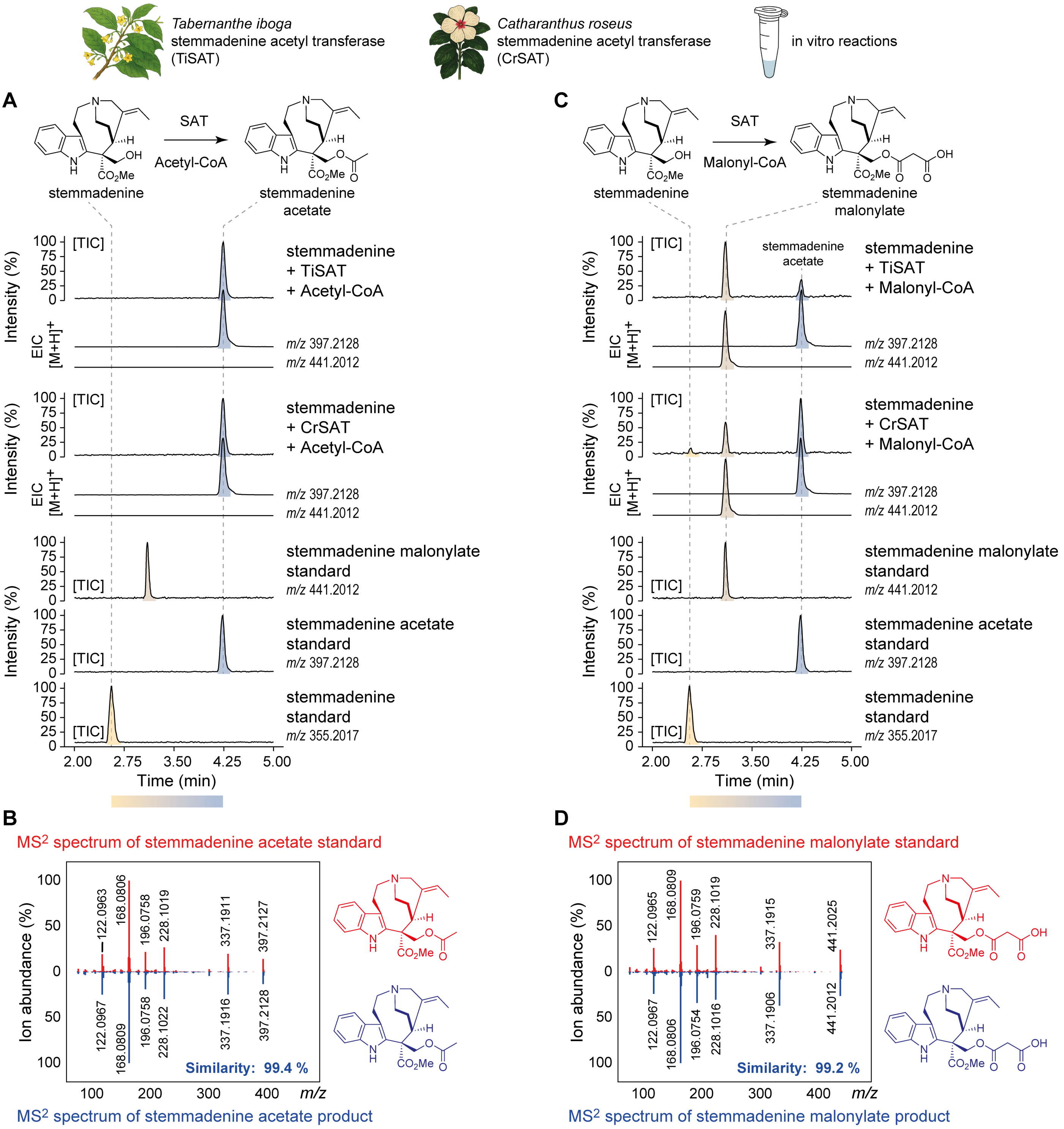


Abbreviation: TIC = Total ion chromatogram, EIC = Extracted ion chromatogram

**Figure S7. Functional characterization of *Tabernanthe iboga* stemmadenine acetyltransferase (TiSAT) in vitro.** (**A**) LC-MS chromatograms of in vitro enzymatic assays using heterologously expressed TiSAT incubated with the substrate stemmadenine and acetyl-CoA as the acyl donor. The previously characterized *Catharanthus roseus* stemmadenine acetyltransferase (CrSAT) (3, 4) was incubated as a positive control under identical conditions. TiSAT catalyzed the acetylation of stemmadenine to yield stemmadenine acetate, consistent with the activity of CrSAT. (**B**) $MS^2$ spectrum of stemmadenine acetate generated by TiSAT, compared with that of the authentic standard. (**C**) LC-MS chromatograms of in vitro assays of TiSAT incubated with stemmadenine and malonyl-CoA as donor substrate. CrSAT was assayed in parallel. Both TiSAT and CrSAT catalyzed the malonylation of stemmadenine to produce stemmadenine malonylate. In addition, stemmadenine acetate was detected in both reactions, likely due to the co-purification of acetyl-CoA from the heterologous expression systems. (**D**) $MS^2$ spectra of stemmadenine malonylate produced in the presence of malonyl-CoA by SAT, confirming the identity of the product.

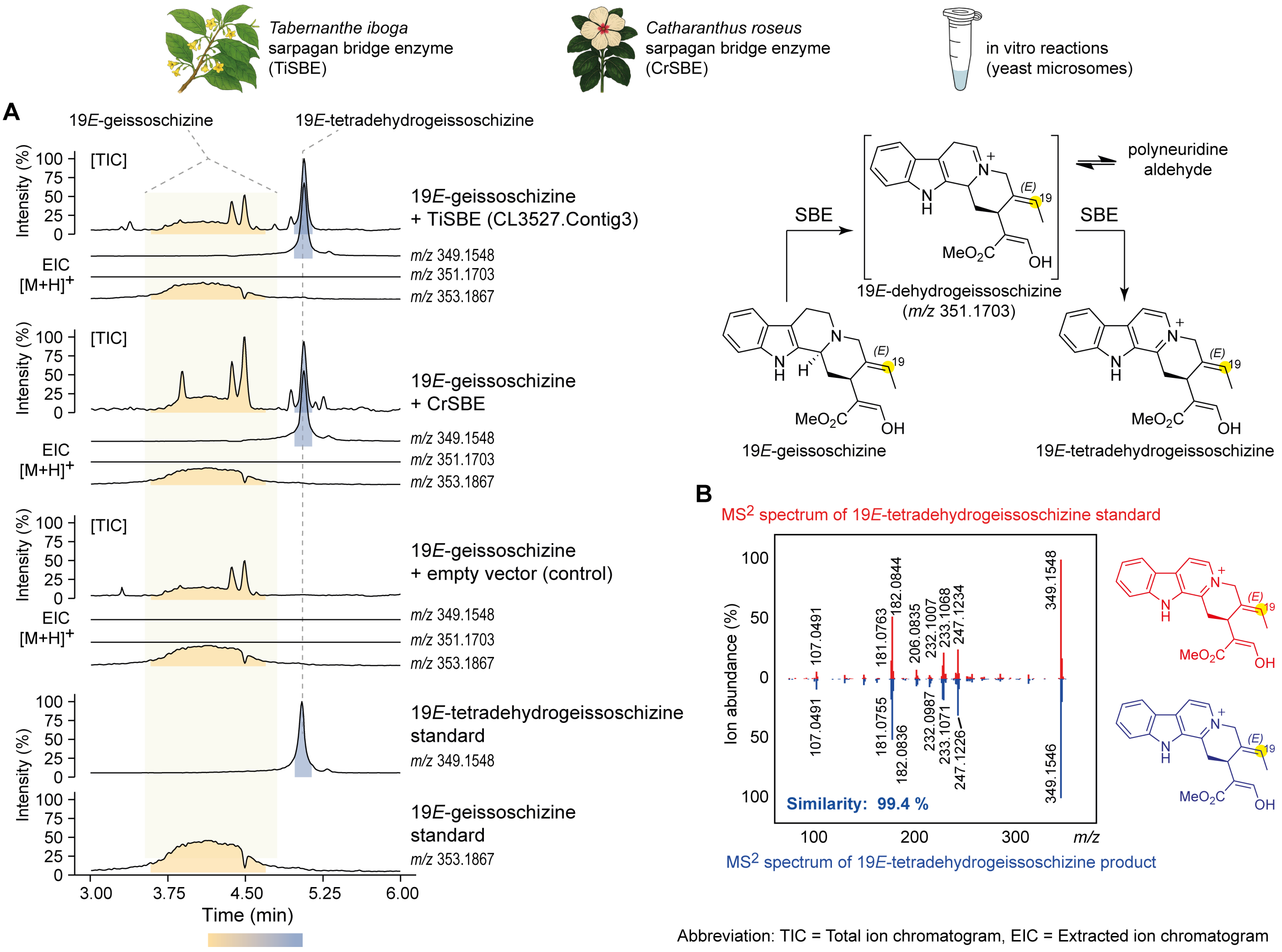


**Figure S8. Functional characterization of *Tabernanthe iboga* sarpagan bridge enzyme (TiSBE) enzyme activity in vitro using 19*E*-geissoschizine.** (**A**) LC-MS chromatograms of in vitro enzymatic assays using yeast microsomes expressing TiSBE and previously characterized *Catharanthus roseus* SBE (CrSBE) (5). Microsomal fractions were assayed with 19*E*-geissoschizine as substrate under identical conditions. CrSBE, previously characterised, was used as a positive control (5). Assays revealed that TiSBE catalyzed the formation of 19*E*-tetradehydrogeissoschizine via the unstable intermediate 19*E*-dehydrogeissoschizine, consistent with the activity observed for CrSBE. (**B**) MS$^2$ spectrum of 19*E*-tetradehydrogeissoschizine produced by TiSBE, compared with that of the authentic standard.

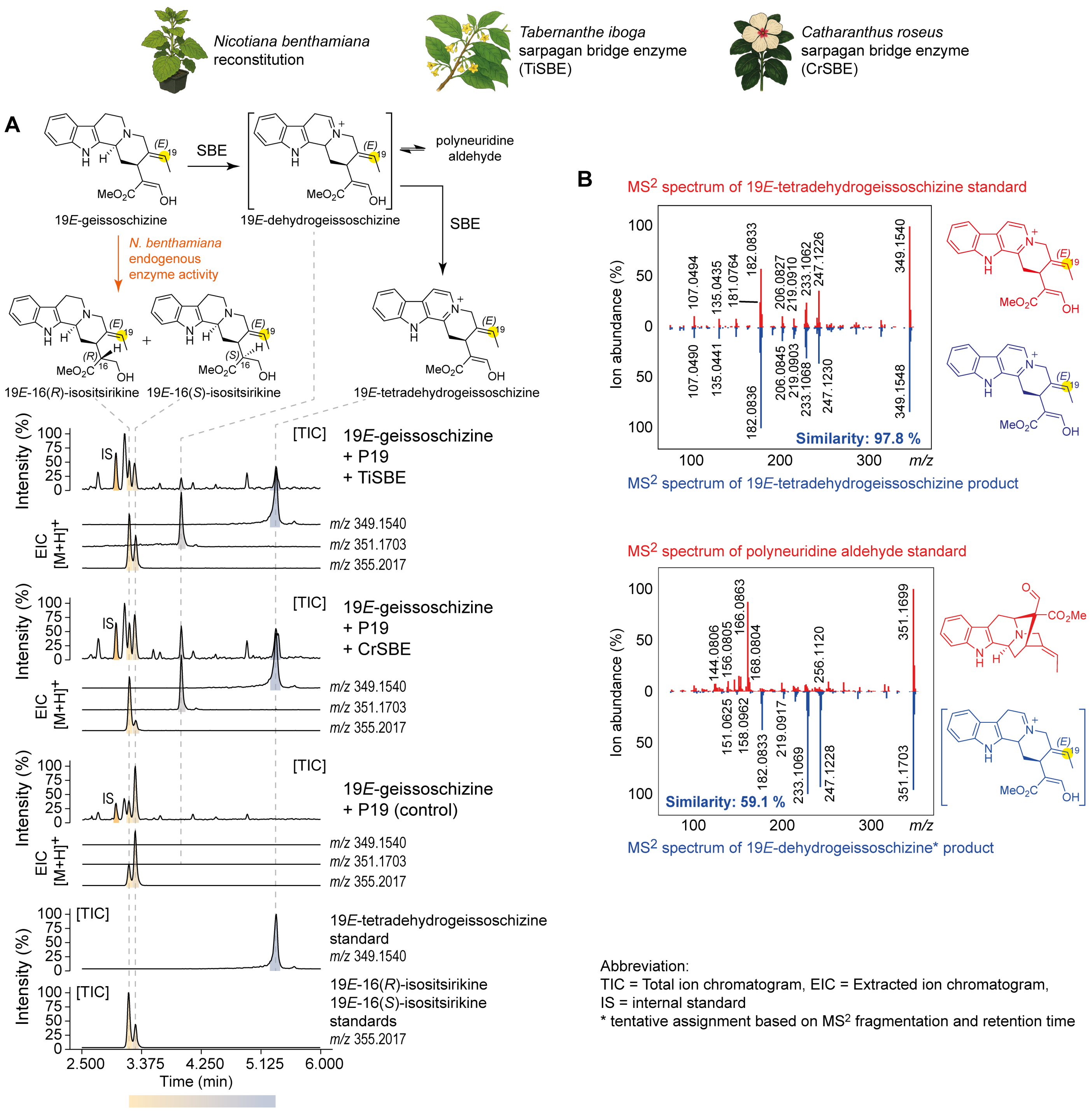


**Figure S9. Functional characterization of *Tabernanthe iboga* sarpagan bridge enzyme (TiSBE) activity in *Nicotiana benthamiana* using 19*E*-geissoschizine as substrate.** (**A**) LC-MS chromatograms of TiSBE transiently expressed in *N. benthamiana* with the P19 viral silencing suppressor, assayed in leaf-disks supplied with the substrate 19*E*-geissoschizine. The previously characterised *Catharanthus roseus* SBE (CrSBE) (5) was included as a positive control under identical conditions. Similar to CrSBE, TiSBE catalyzed the formation of 19*E*-tetradehydrogeissoschizine via the unstable intermediate 19*E*-dehydrogeissoschizine. The final product was confirmed using an authentic standard, while the intermediate was annotated based on its $MS^2$ fragmentation pattern. Notably, the endogenous activity of *N. benthamiana* metabolic activity resulted in the reduction of 19*E*-geissoschizine to 19*E*-isositsirikine isomers. (**B**) $MS^2$ spectrum of 19*E*-tetradehydrogeissoschizine produced by TiSBE, compared with the spectrum of the authentic standard. $MS^2$ spectrum of the 19*E*-dehydrogeissoschizine differs from that of polyneuridine aldehyde, which is known to exist in equilibrium with 19*E*-dehydrogeissoschizine, further supporting its structural annotation.

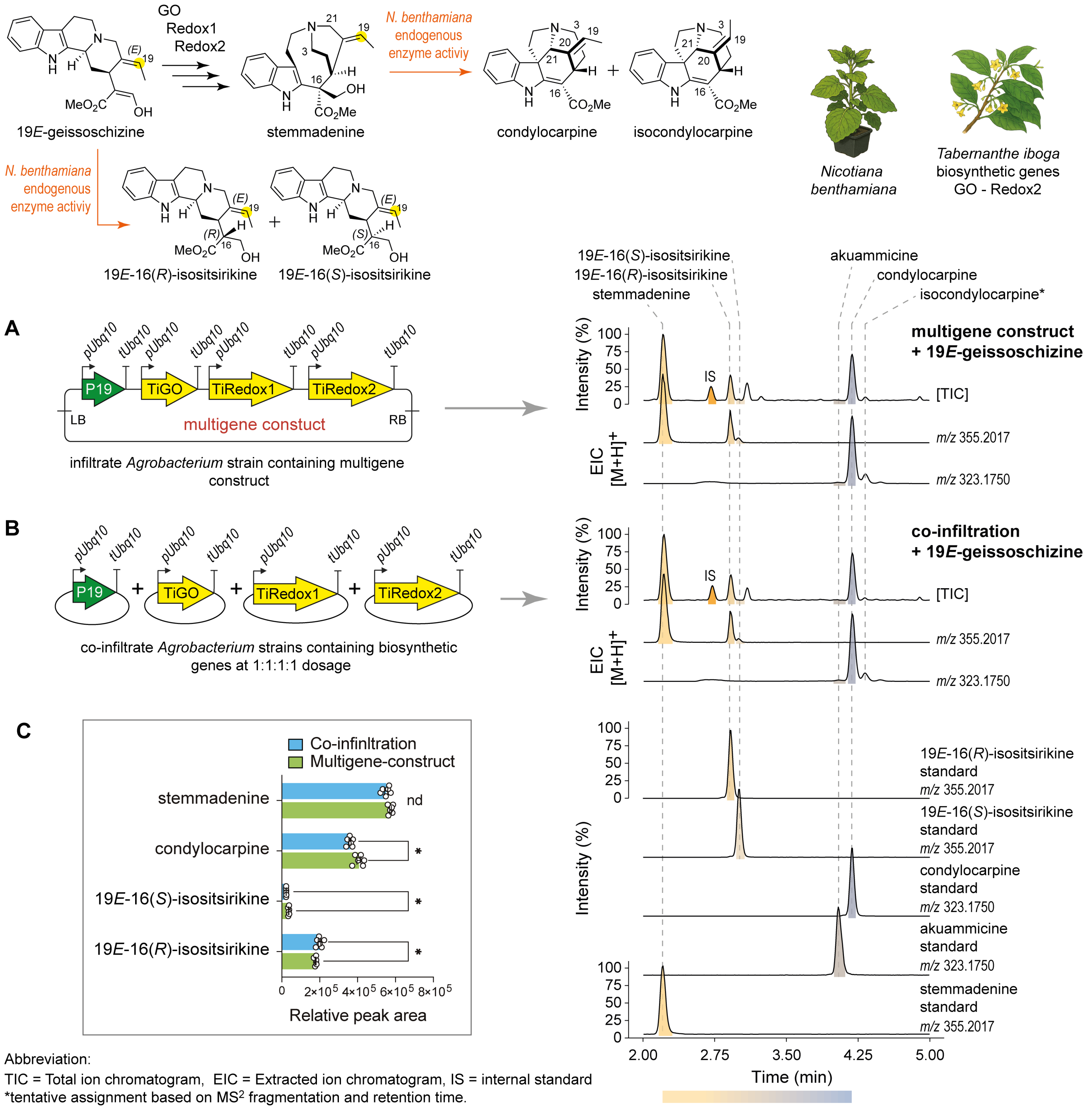


**Figure S10. Biosynthesis of stemmadenine in *Nicotiana benthamiana* using a multigene construct of *Tabernanthe iboga* biosynthetic genes.** (**A**) A multigene construct comprising *T. iboga* biosynthetic genes GO, Redox1, and Redox2, along with the P19 viral silencing suppressor, enables the production of stemmadenine in *N. benthamiana* when the substrate 19*E*-geissoschizine is supplied exogenously. A Representative LC-MS chromatogram of the biosynthetic product, stemmadenine, is shown. (**B**) Co-infiltration of individual GO, Redox1, and Redox2 constructs with P19 in *N. benthamiana* also resulted in the production of stemmadenine, as shown in the representative LC-MS chromatogram. (**C**) Comparison of stemmadenine yields from multigene versus co-infiltration-based transient expression systems, using equivalent substrate concentrations. No statistically significant difference (nd) in stemmadenine production was observed between the two expression strategies (unpaired t-test, $n$ = 6 biological replicates). However, the multigene construct produced statistically lower (*) levels of undesired side-products, resulting from endogenous *N. benthamiana* metabolic activity (unpaired t-test, *$P < 0.05$, nd = not significant $P > 0.05$). Relative peak area is normalized to the internal standard (IS). Bar graphs are ploted as the mean ± S.D.

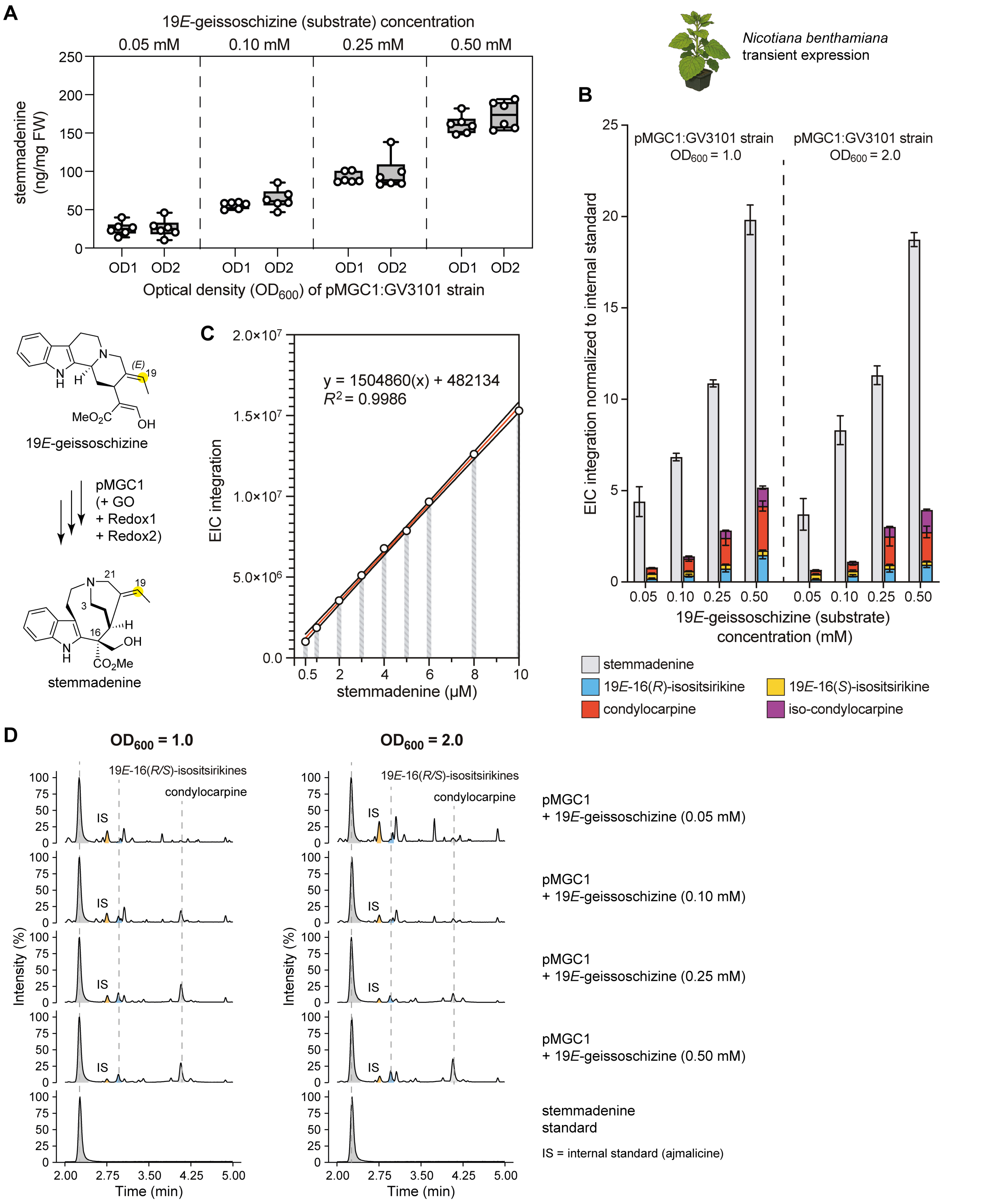


**Figure S11. Quantification of stemmadenine production in *Nicotiana benthamiana* using a *Tabernanthe iboga* multigene construct with 19*E*-geissoschizine feeding. (A)** Stemmadenine titers from leaves transiently expressing pMGC1 at *Agrobacterium* densities $OD_{600}$ = 1.0 or 2.0, followed by exogenous 19*E*-geissoschizine at 0.05, 0.10, 0.25, or 0.50 mM. Tissue was harvested 12–16 h post-substrate infiltration (n = 6 biological replicates). (**B**) Qualitative product profiles (stemmadenine and co-produced MIAs) normalized to internal standard and dilution factor. (**C**) Calibration curve for stemmadenine quantification; best-fit line (red) with 95% confidence interval (black). (**D**) Representative LC–MS chromatograms illustrating product profile under the conditions in (A).

**A**

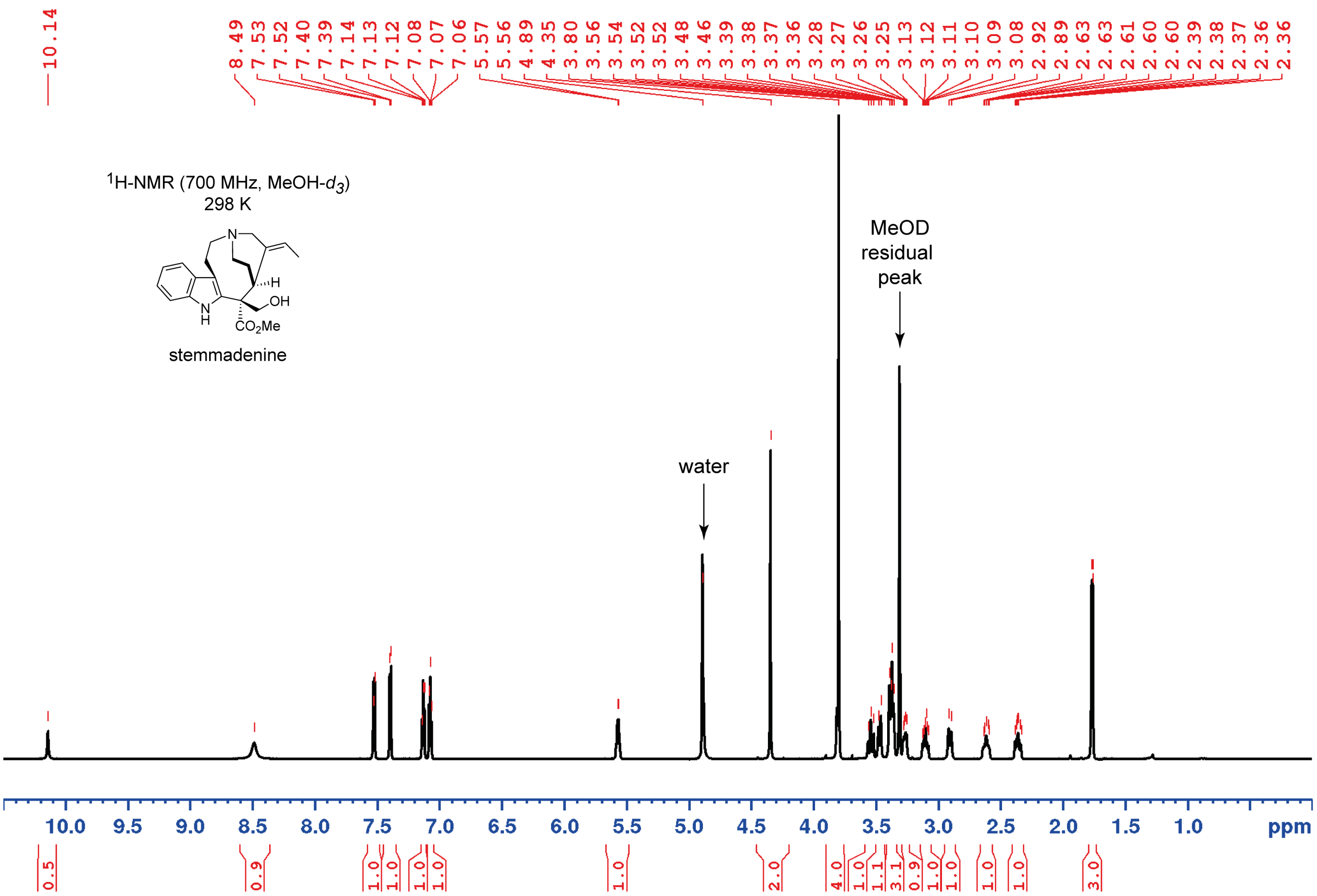


**B**

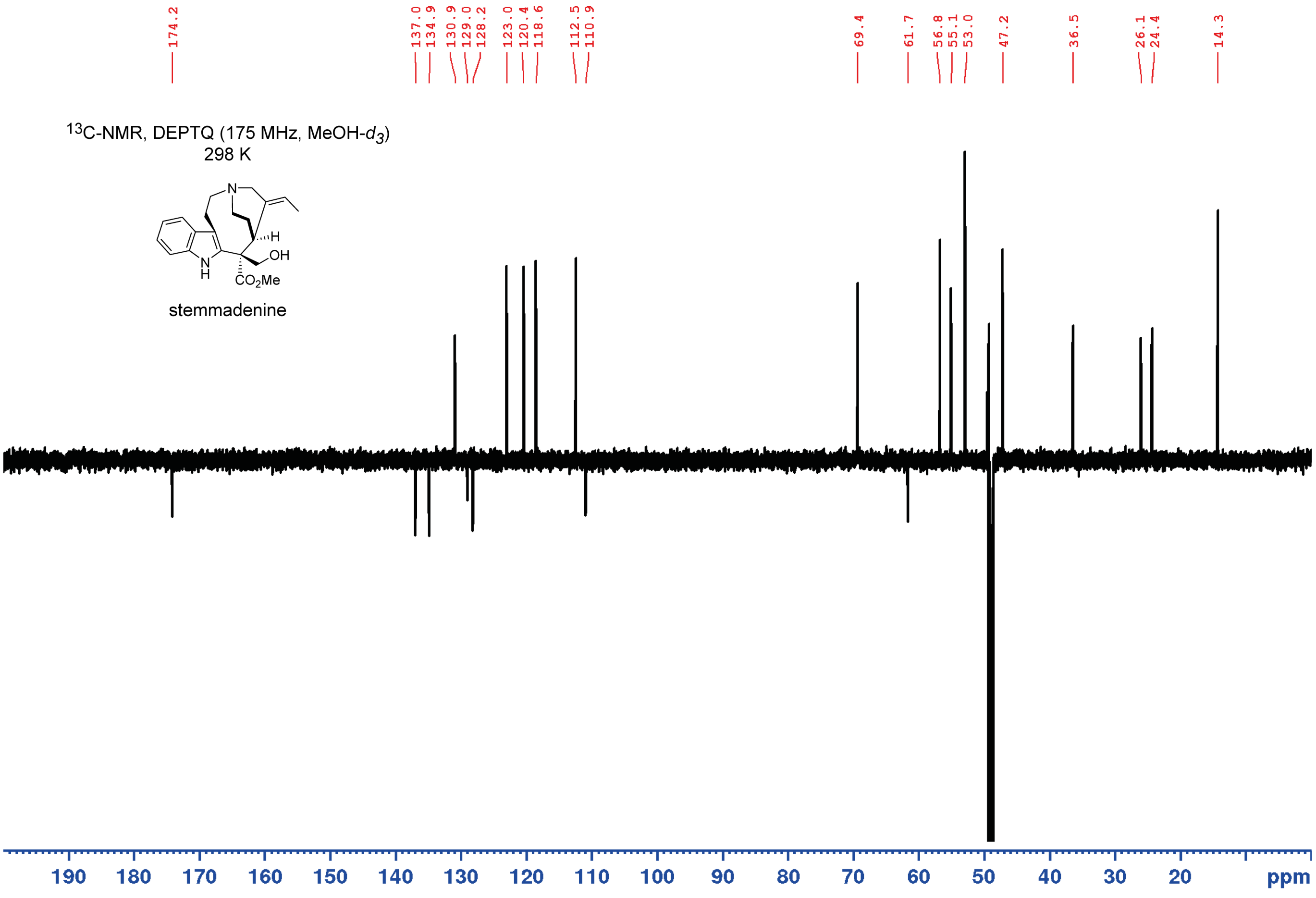


**Figure S12. NMR of stemmadenine.** (A) $^{1}$H NMR with water presaturation full range in MeOH-$d_3$ of stemmadenine as formate. (B) DEPTQ full range in MeOH-$d_3$ of stemmadenine as formate.

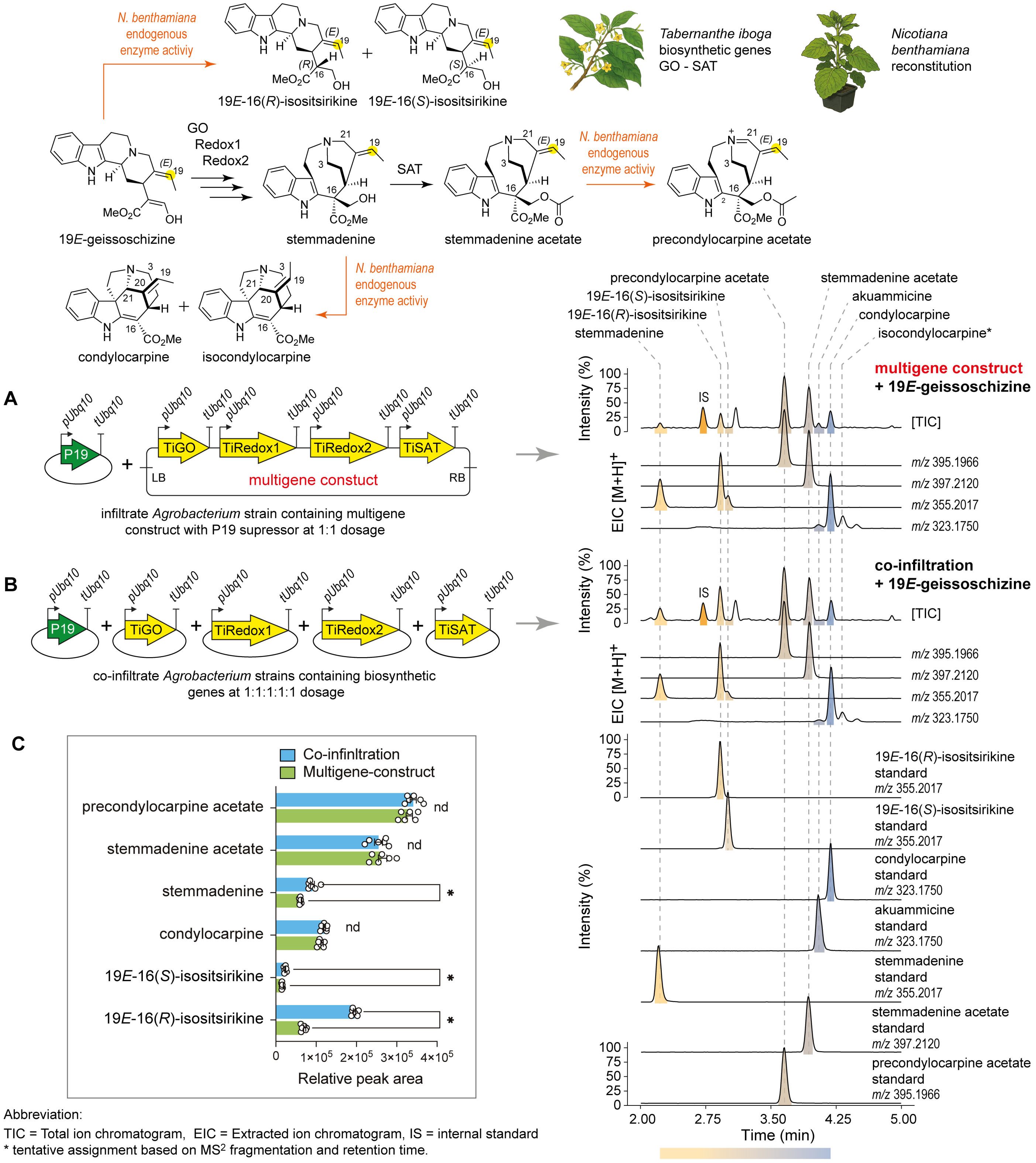


**Figure S13. Biosynthesis of stemmadenine acetate in *Nicotiana benthamiana* using a multigene construct of *Tabernanthe iboga* biosynthetic genes.** (**A**) A multigene construct comprising *T. iboga* biosynthetic genes GO, Redox1, Redox2, and SAT, co-infiltrated with P19 viral silencing suppressor, enables the production of stemmadenine acetate in *N. benthamiana* upon exogenous supply of 19*E*-geissoschizine substrate. LC-MS chromatograms of the biosynthetic product, along with side-products, are shown. In *N. benthamiana*, stemmadenine acetate is readily oxidized to precondylocarpine acetate by endogenous metabolic activity. (**B**) Co-infiltration of individual GO, Redox1, Redox2, and SAT constructs with P19 also resulted in stemmadenine acetate production, as shown in the corresponding LC-MS chromatogram. (**C**) Comparison of stemmadenine acetate yields between the multigene and co-infiltration-based transient expression strategies, using equivalent substrate concentrations. No statistical difference (nd) was observed in stemmadenine acetate production between the two approaches (unpaired t-test, $n = 6$ biological replicates). However, the multigene construct produced statistically lower (*) levels of undesired side-products, arising from endogenous *N. benthamiana* metabolic activity (unpaired *t-test*, *$P < 0.05$, nd = not significant, $P > 0.05$, n = 6 biological replicates). Relative peak area is normalized to the internal standard (IS). Bar graphs are ploted as the mean ± S.D.

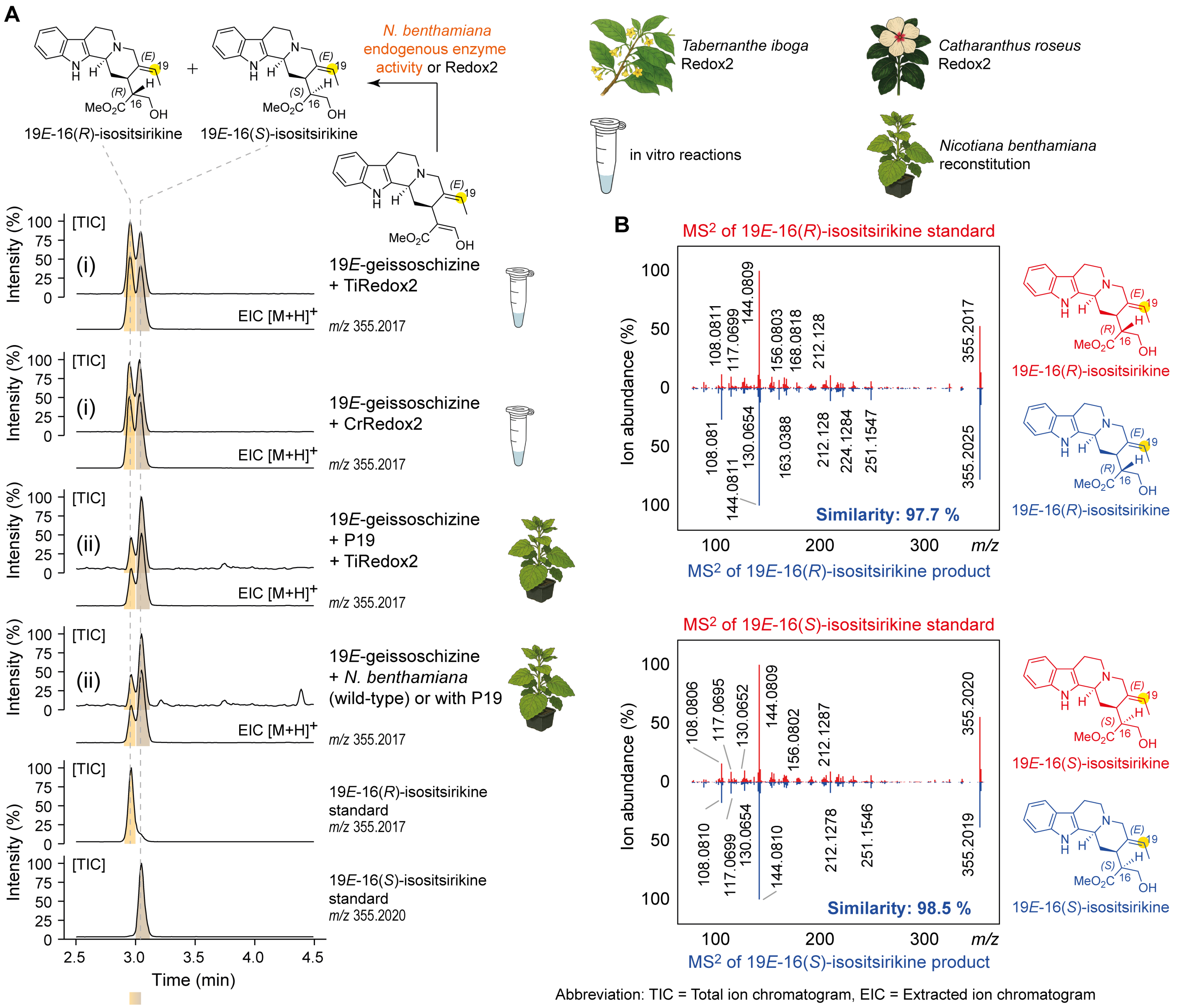


**Figure S14. Functional characterization of *Tabernanthe iboga* redox enzyme 2 (TiRedox2) activity using 19*E*-geissoschizine.** (**A**) (i) LC-MS chromatograms of in vitro enzymatic assays using heterologously expressed TiRedox2 and *Catharanthus roseus* (CrRedox2), assayed with the substrate 19*E*-geissoschizine. Both TiRedox2 and CrRedox2 catalyzed the reduction of 19*E*-geissoschizine to the isomeric products 19*E*-16(*R*)-isositsirikine and 19*E*-16(*S*)-isositsirikine. The absolute structure of 19*E*-16(*R*)-isositsirikine and 19*E*-16(*S*)-isositsirikine was confirmed after purification and NMR analysis. (ii) LC-MS chromatograms showing that endogenous *Nicotiana benthamiana* metabolic activity also catalyzes the reduction of 19*E*-geissoschizine to the identical 19*E*-isositsirikine isomers. (**B**) $MS^2$ spectra of 19*E*-16(*R*)-isositsirikine and 19*E*-16(*S*)-isositsirikine isomers generated by TiRedox2, shown alongside spectra of authentic synthetic standards.

A

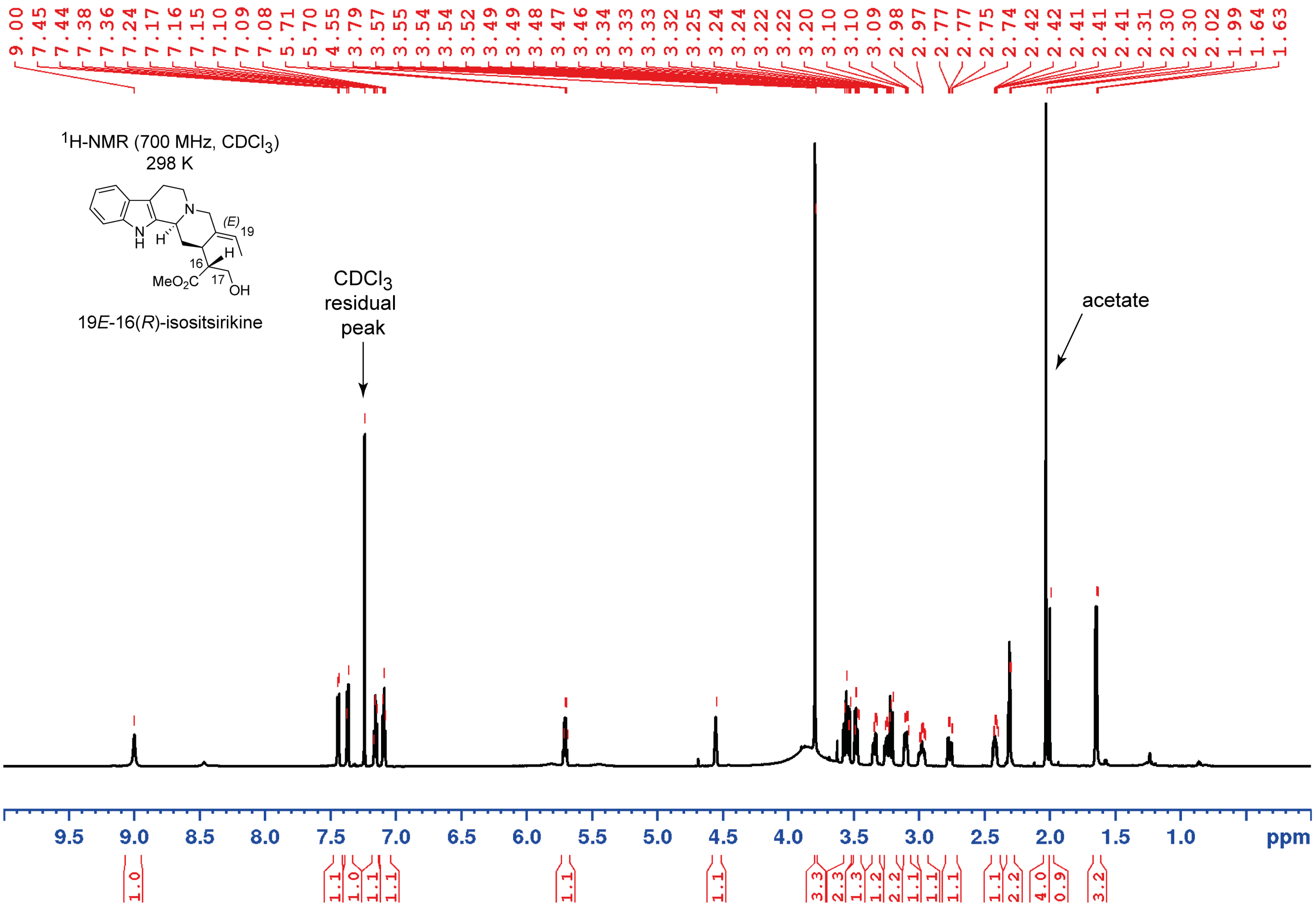


B

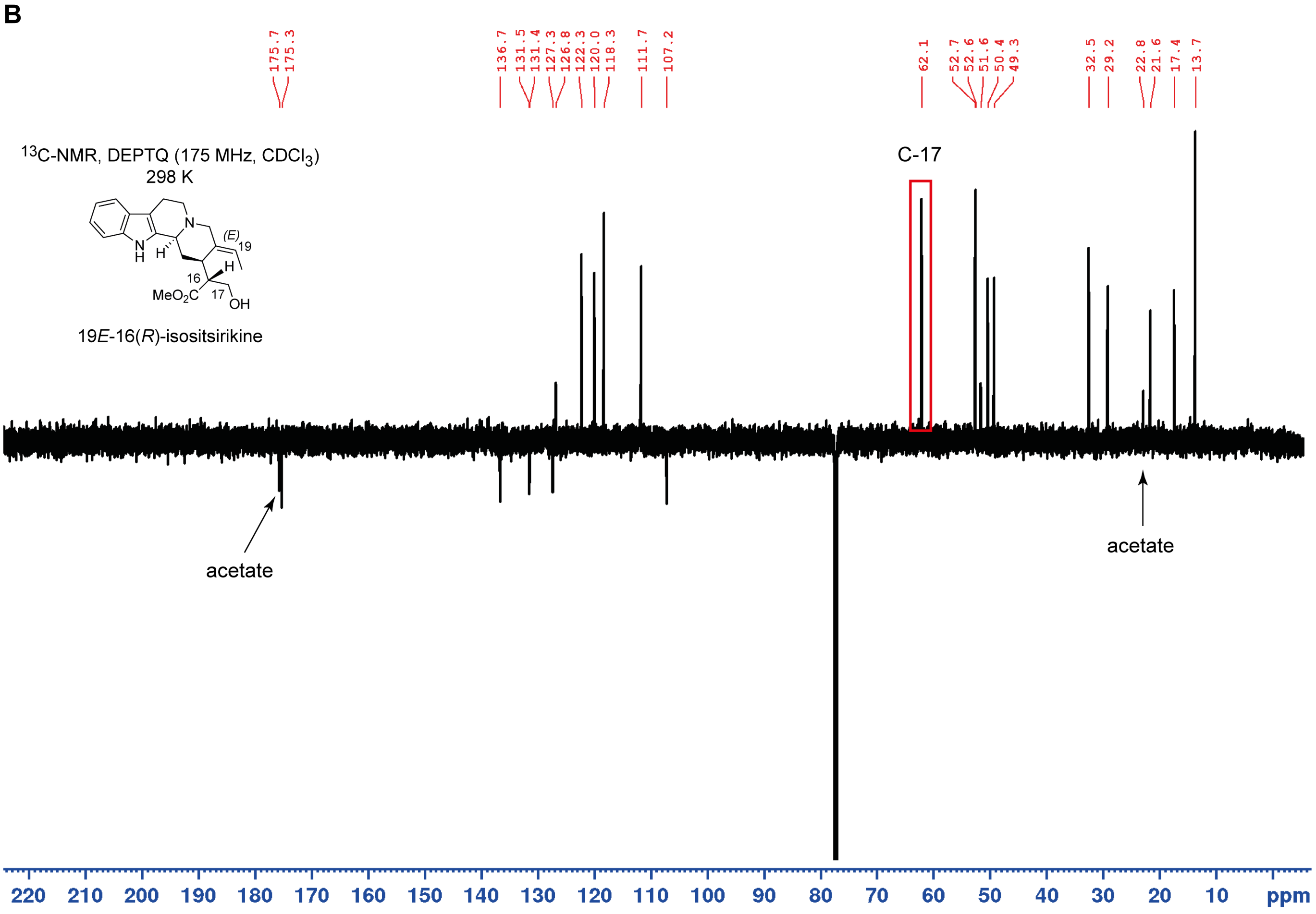


**Figure S15. NMR of 19*E*-16(*R*)-isositsirikine.** (**A**) $^{1}$H NMR with water presaturation, full range in $CDCl_3$ of 19*E*-16(*R*)-isositsirikine as acetate. (**B**) DEPTQ full range in $CDCl_3$ of 19*E*-16(*R*)-isositsirikine as acetate.

**A**

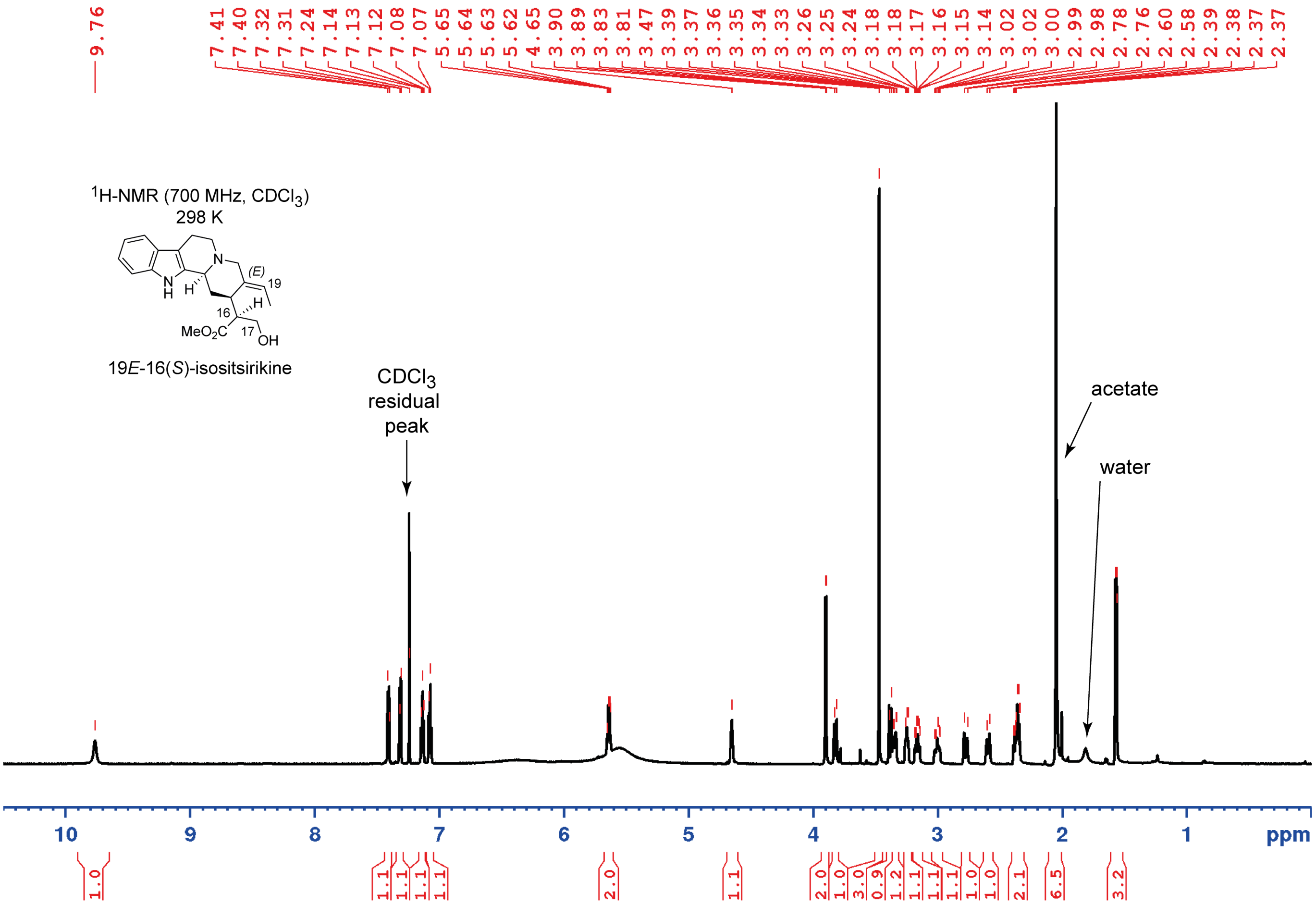


**B**

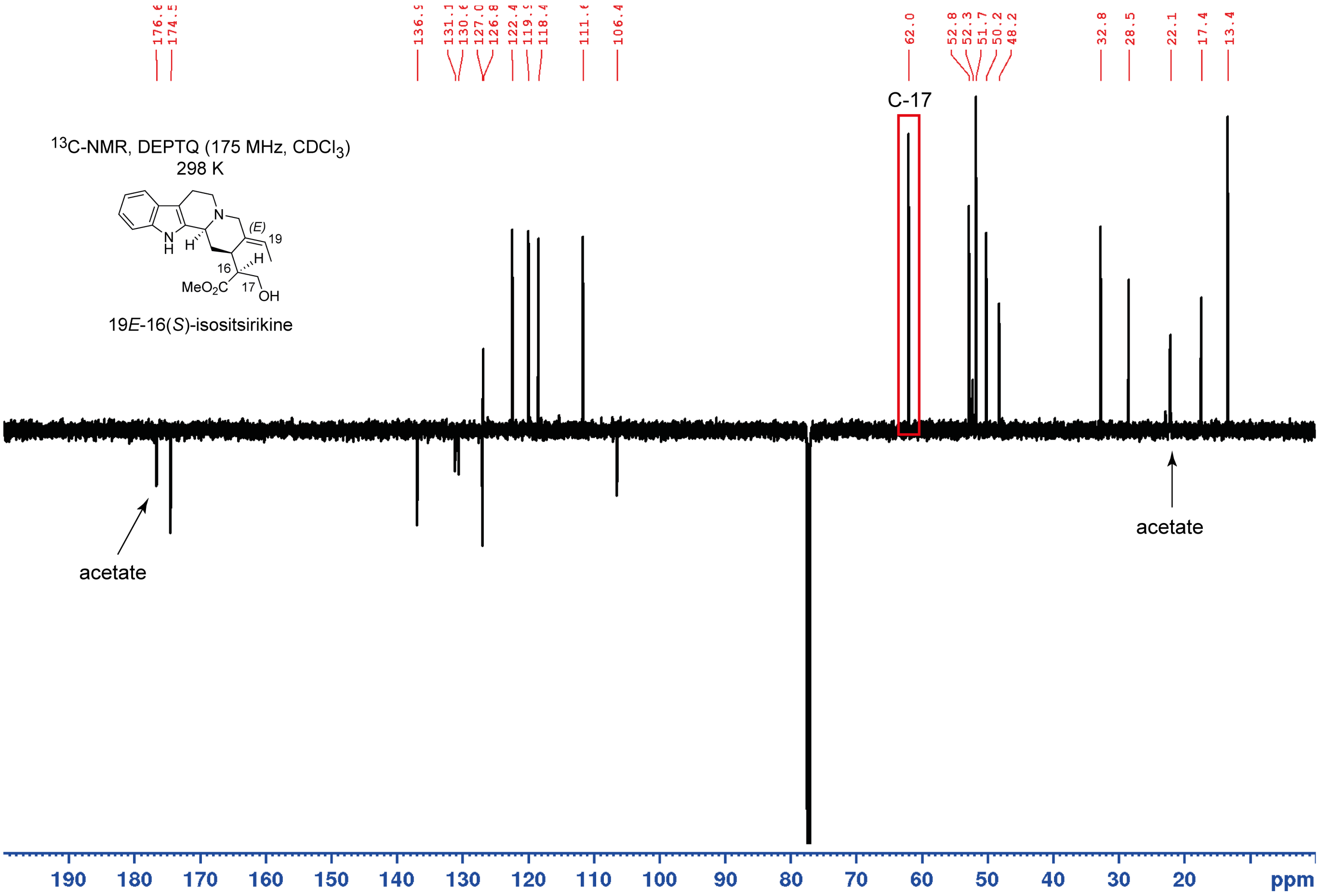


**Figure S16. NMR of 19*E*-16(*S*)-isositsirikine.** (**A**) $^{1}$H NMR with water presaturation, full range in $CDCl_3$ of 19*E*-16(*S*)-isositsirikine as acetate. (**B**) DEPTQ full range in $CDCl_3$ of 19*E*-16(*S*)-isositsirikine as acetate.

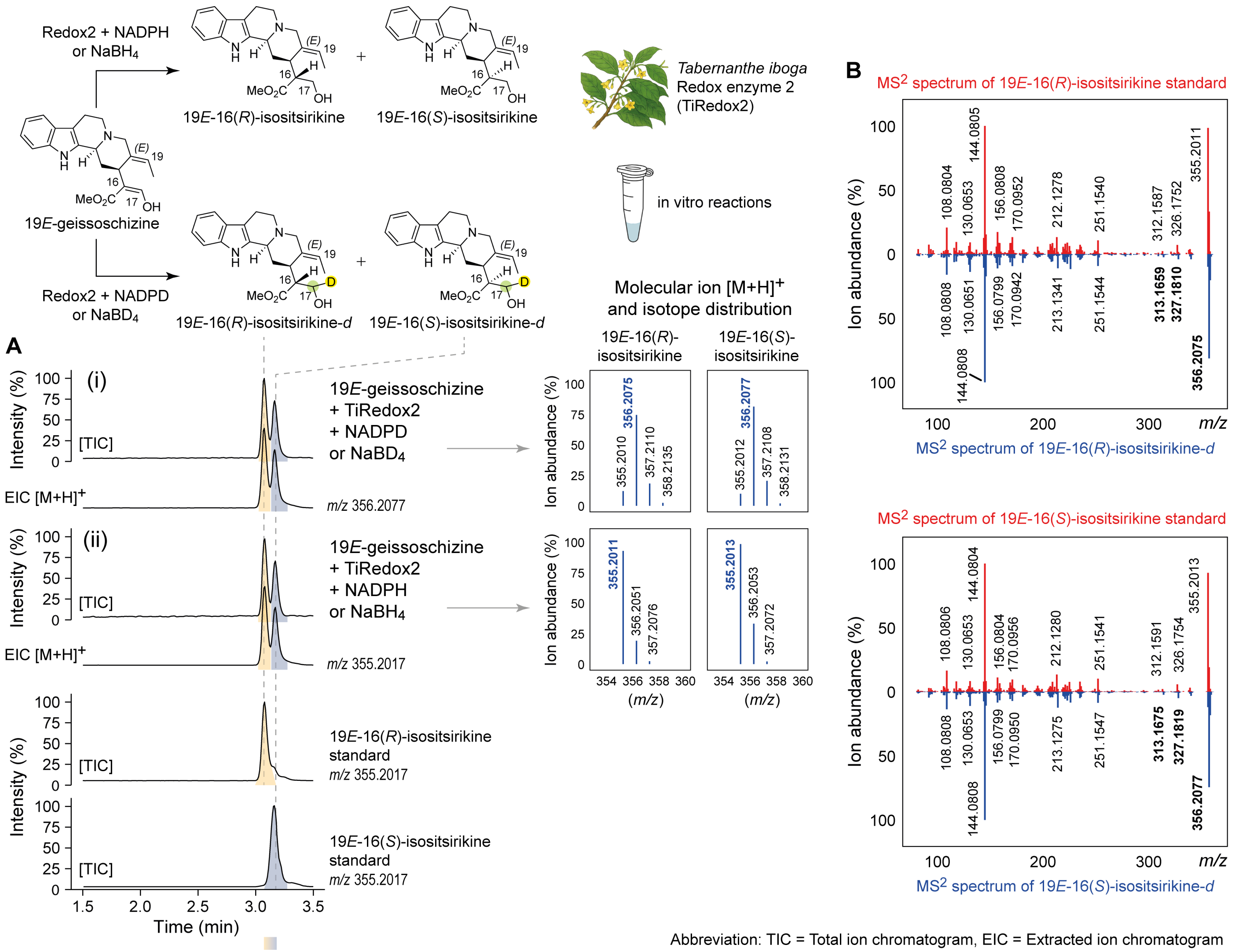


**Figure S17. Deuterium-labelled (-*d*) reduction assays of 19*E*-geissoschizine.** (**A**) (i) LC-MS chromatograms and the molecular ion mass spectra of 19*E*-16(*R*)-isositsirikine-*d* and 19*E*-16(*S*)-isositsirikine-*d* isomers generated by the chemical reduction of 19*E*-geissoschizine using $NaBD_4$ or by enzymatic reduction with TiRedox2 in the presence of isotopically labelled NADPH (NADPD). (ii) LC-MS chromatograms and the molecular ion mass spectra of the corresponding unlabeled 19*E*-16(*R*)-isositsirikine and 19*E*-16(*S*)-isositsirikine isomers, generated by the chemical reduction of 19*E*-geissoschizine using $NaBH_4$ or enzymatic assays using TiRedox2 with NADPH. (**B**) $MS^2$ spectra of the deuterium labelled isomers 19*E*-16(*R*)-isositsirikine-*d* and 19*E*-16(*S*)-isositsirikine-*d* compared to those of the authentic 19*E*-isomers standards, highlighting the incorporation of deuterium into each isomer and its corresponding fragment.

A

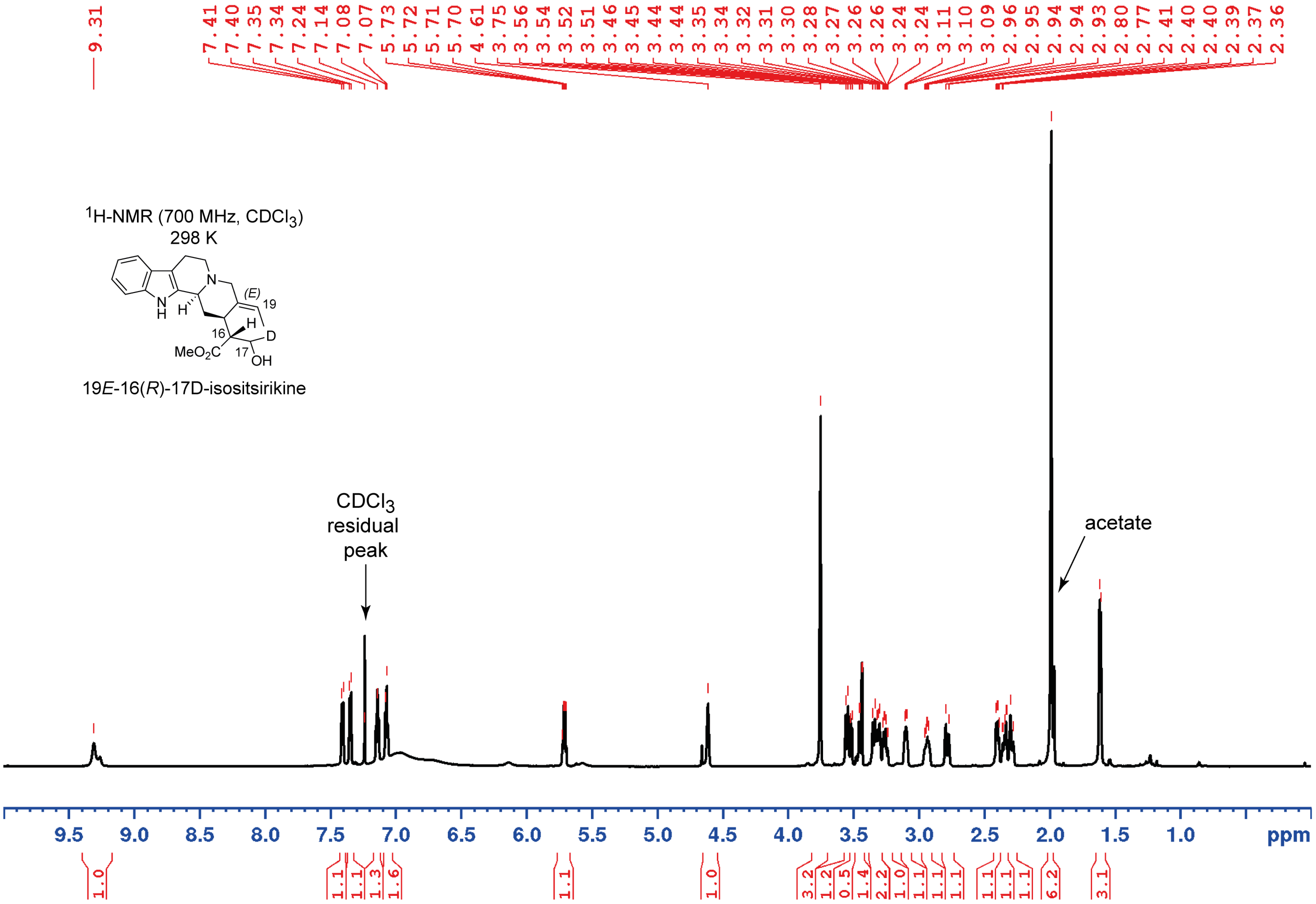


B

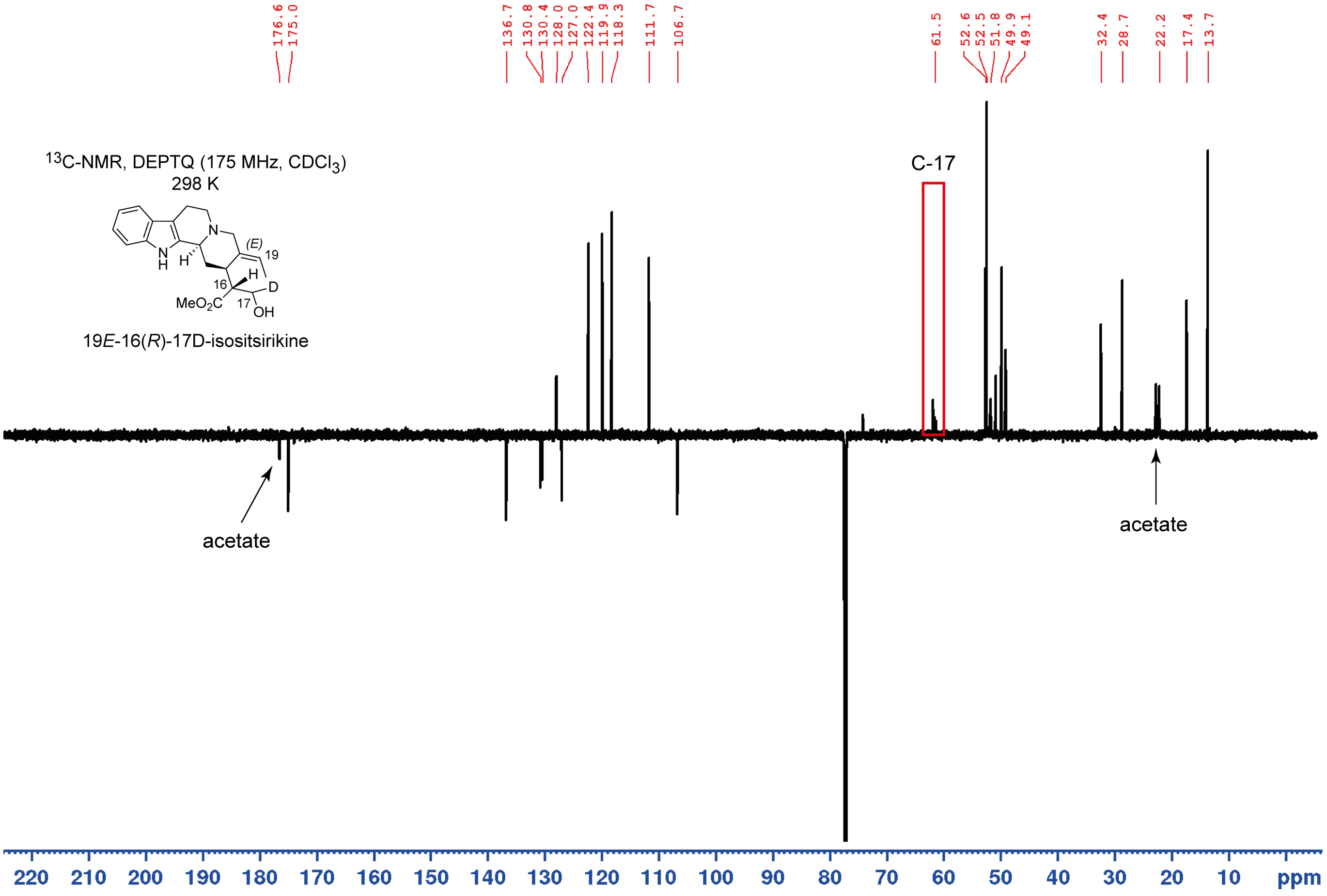


**Figure S18. NMR of 19*E*-16(*R*)-17D-isositsirikine.** (**A**) $^1$H NMR with water presaturation, full range in $CDCl_3$ of 19*E*-16(*R*)-17D-isositsirikine as acetate. (**B**) DEPTQ full range in $CDCl_3$ of 19*E*-16(*R*)-17D-isositsirikine as acetate.

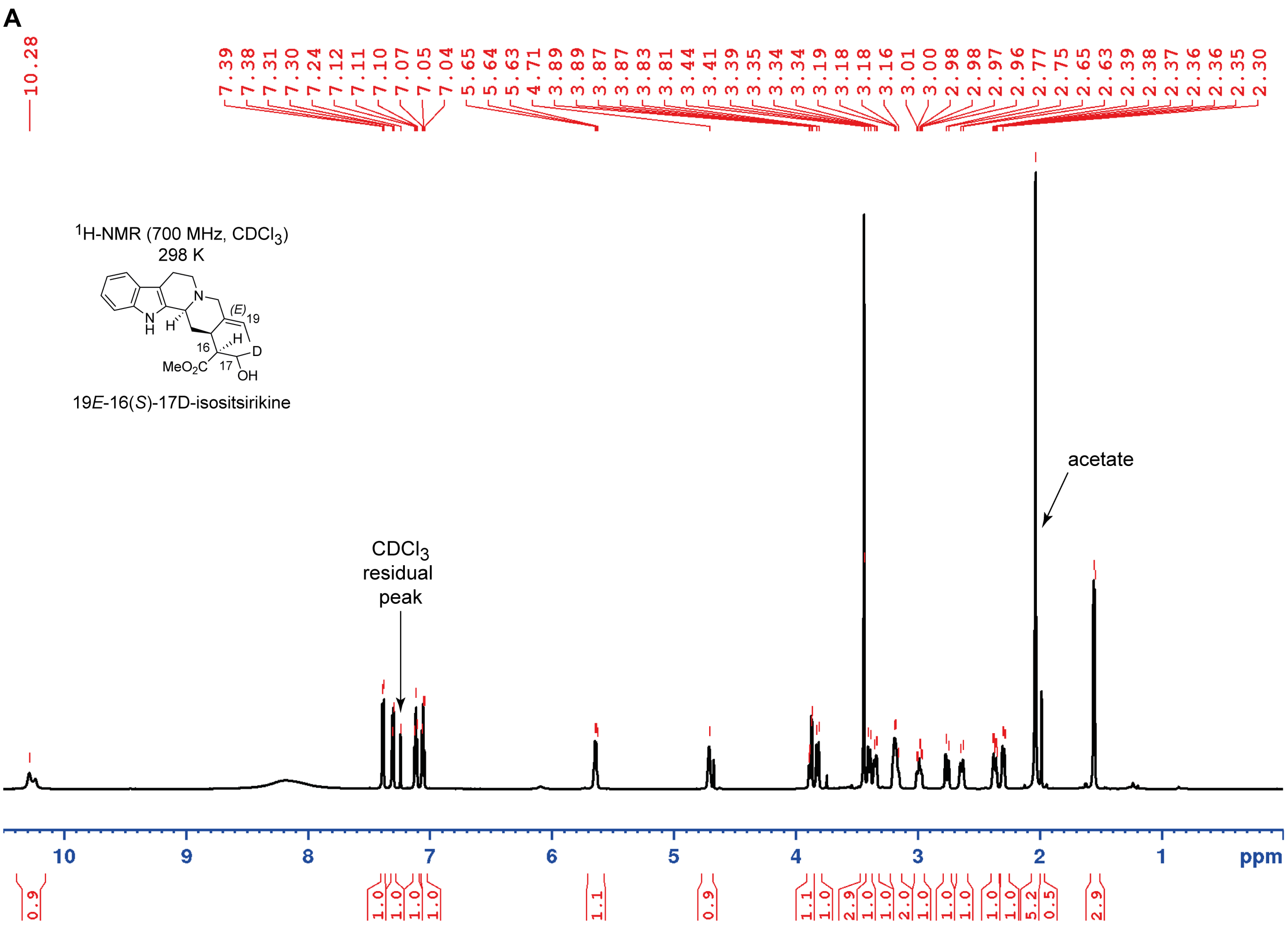


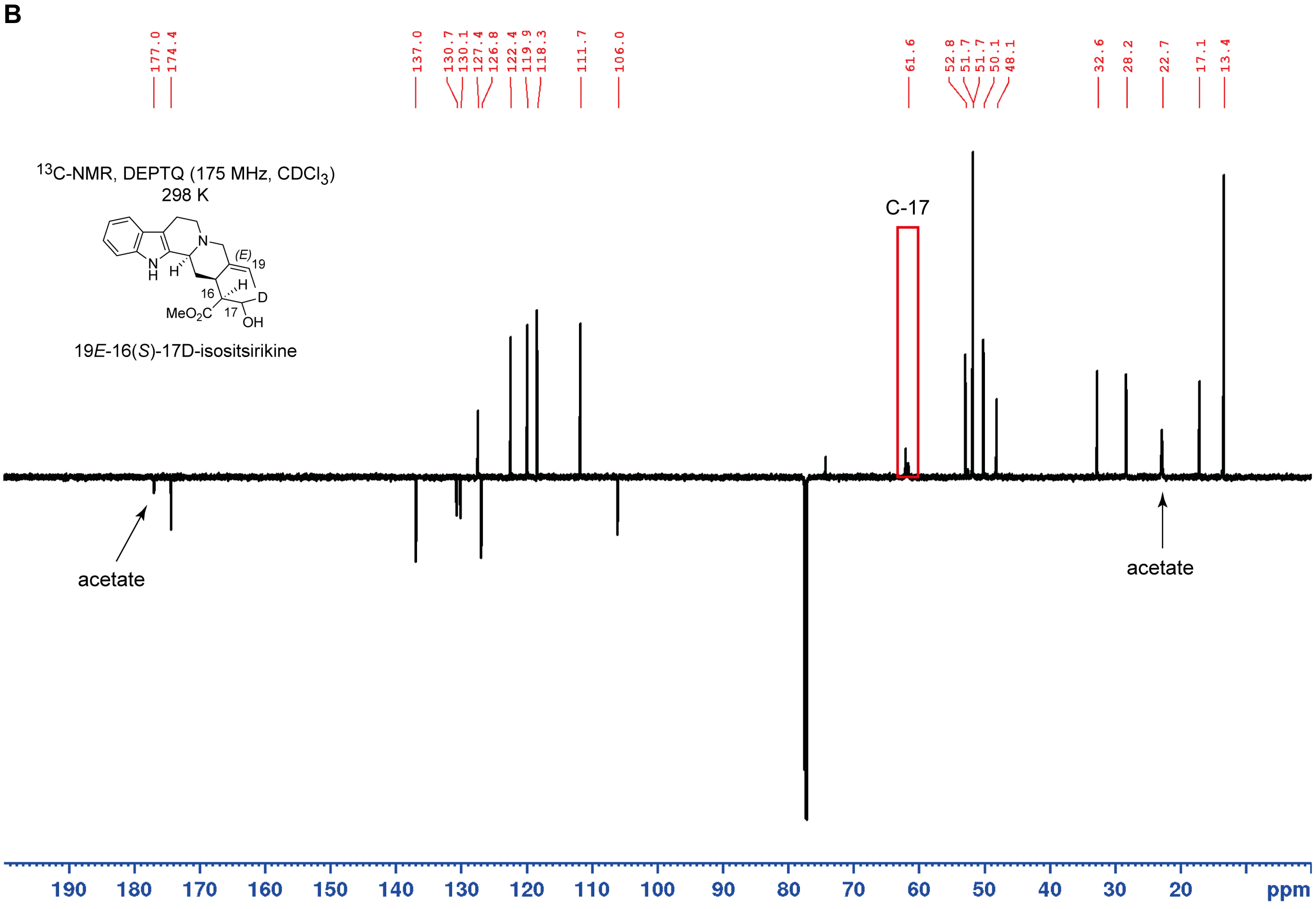


**Figure S19. NMR of 19*E*-16(*S*)-17D-isositsirikine.** (**A**) $^1$H NMR with water presaturation, full range in $CDCl_3$ of 19*E*-16(*S*)-17D-isositsirikine as acetate. (**B**) DEPTQ full range in $CDCl_3$ of 19*E*-16(*S*)-17D-isositsirikine as acetate.

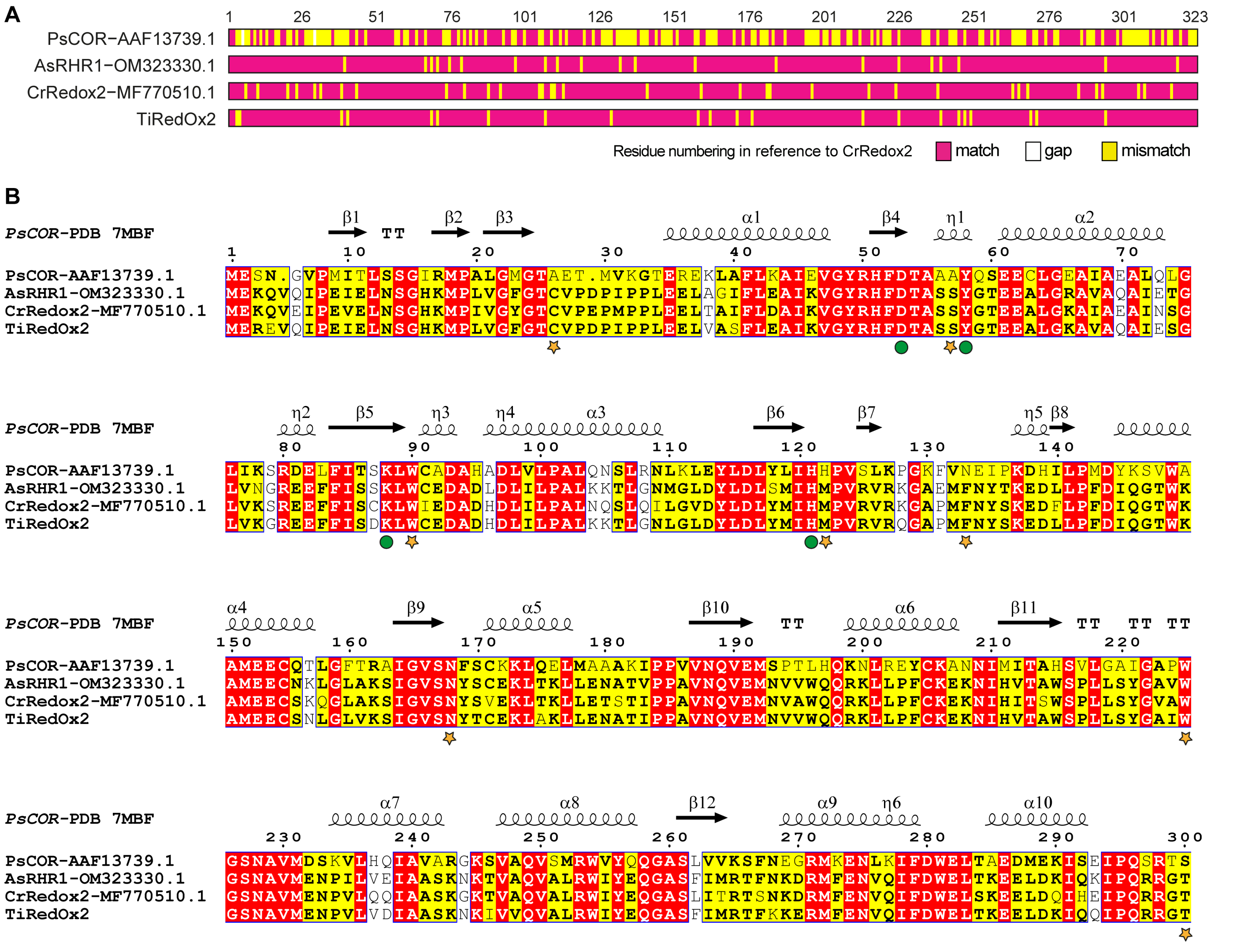


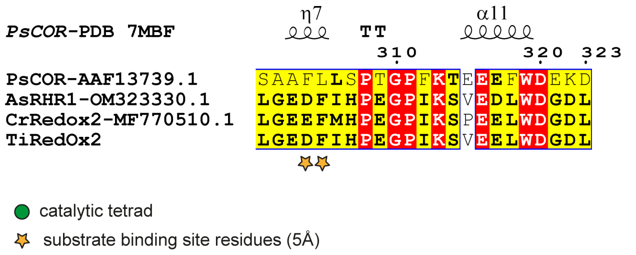


| | PsCOR−AAF13739.1 | AsRHR1−OM323330.1 | CrRedox2−MF770510.1 | TiRedOx2 |
|---|---|---|---|---|
| PsCOR−AAF13739.1 | | 48.61 | 49.23 | 49.23 |
| AsRHR1−OM323330.1 | 48.61 | | 82.66 | 90.71 |
| CrRedox2−MF770510.1 | 49.23 | 82.66 | | 82.97 |
| TiRedOx2 | 49.23 | 90.71 | 82.97 | |

**Figure S20. Multiple sequence alignment of aldo-keto reductase (AKR) enzymes involved in plant alkaloid biosynthesis.** (**A**) Schematic representation of amino acid sequence alignment of functionally characterised AKR enzymes, the enzyme family to which Redox2 belongs in the stemmadenine biosynthetic pathway. (**B**) Detailed alignment of *T. iboga* Redox2 with orthologous AKR enzymes involved in monoterpene indole alkaloid (MIA) biosynthesis (3, 6), annotated with structural features based on the closest structural homolog, *Papaver somniferum* codeinone reductase (PsCOR) (7). Key residues comprising the catalytic tetrad and those involved in substrate binding pocket interactions are highlighted (8). The percentage amino acid identity between TiRedox2 and the other plant AKR enzymes is also shown. GenBank accession numbers for all functionally characterized AKR enzymes are provided as appropriate.

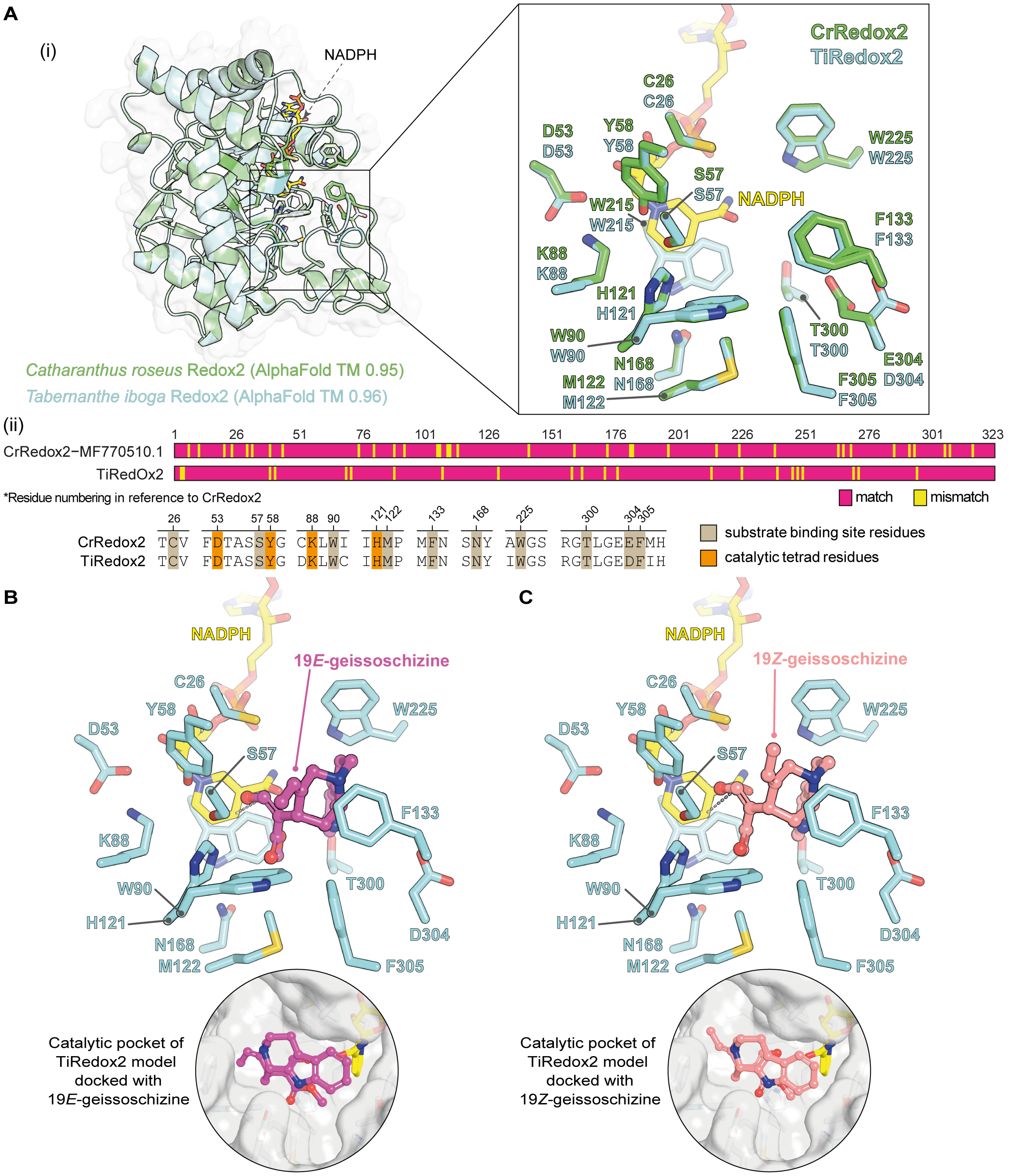


**Figure S21. Structural analysis of *Tabernanthe iboga* redox enzyme 2 (TiRedox2) model.** (**A**) (**i**) Superimposed AlphaFold models of TiRedox2 (TM-score = 0.9514) and *Catharanthus roseus* Redox2 (CrRedox2) (TM score = 0.9634), generated using AlphaFold v3 (9) with NADPH modelled in the cofactor pocket. The conserved catalytic tetrad characteristic of the aldo-keto reductase (AKR) family enzymes, along with residues located within 5 Å of the active site pocket, are visualized in complex with NADPH. (**ii**) Schematic representation of the amino acid sequence alignment between TiRedox2 and CrRedox2 (82.97% identity), highlighting conserved and divergent regions. The catalytic tetrad (Asp53, Tyr58, Lys88, and His121) (8) and putative substrate-binding residues are indicated. (**B**) Molecular docking of the substrate 19*E*-gesissoschizine into the TiRedox2 model, illustrating interactions between the substrate and active site residues. The carbonyl (aldehyde) moiety of 19*E*-geissoschizine (substrate) is positioned adjacent to the catalytic residues and NADPH, consistent with a reductive mechanism for AKRs. (**C**) Molecular docking of the substrate 19*Z*-gesissoschizine into the TiRedox2 model reveals a binding orientation comparable to the 19*E*-isomer. The *Z*-configuration of the ethyl moiety at C20 does not significantly alter the substrate orientation within the active site, suggesting a similar reduction mechanism. Template modelling (TM) scores reflect the overall confidence of the AlphaFold v3 (9) model. Molecular docking was performed using AutoDock Vina (10) using default settings; the best-ranked pose (lowest predicted binding energy and optimal orientation of interaction residues) was selected for analysis.

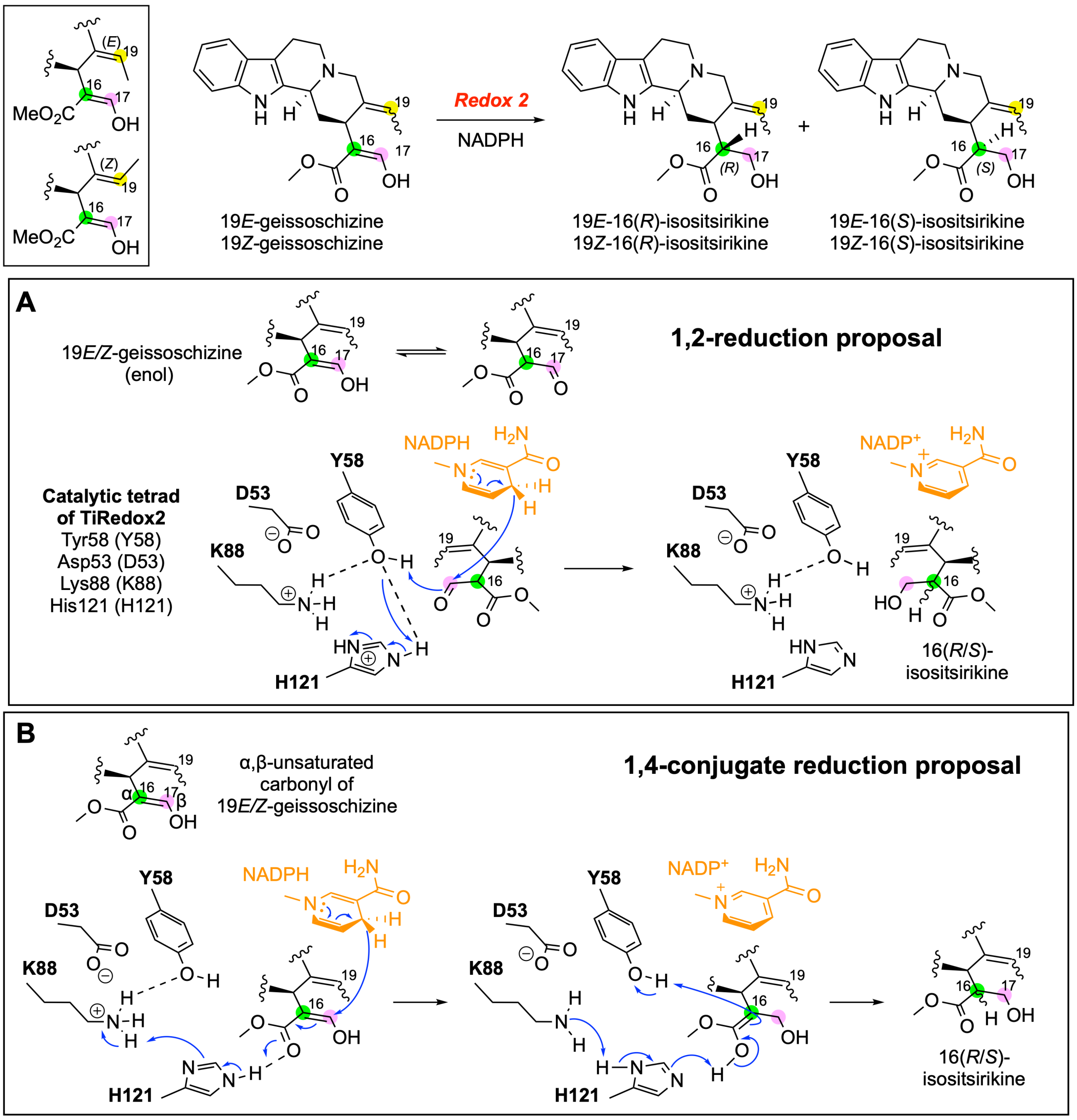


**Figure S22. Mechanistic proposal of 19*E/Z*-geissoschizine reduction catalyzed by Redox2.** *Tabernanthe iboga* aldo-keto reductase (AKR) enzyme Redox2 (TiRedox2) catalyze the NADPH-dependent reduction of 19*E/Z*-geissoschizine at C17, to yield the diastereomeric products 16(*R*)- and 16(*S*)- isositsirikine. (**A**) The reaction proceeds by the canonical "push-pull" reduction mechanism typical to AKRs (8, 11, 12). The conserved catalytic tetrad of TiRedox2, Tyr58, Asp53, Lys88, and His122 plays a crucial role in catalysis to facilitate the push-pull reaction mechanism. The NADPH cofactor binds to the enzyme, followed by a 4-pro-*R* hydride transfer to the substrate aldehyde and protonation of the oxygen by Tyr58 which acts as general acid by participating in the proton relay with His121, completing the 1,2-reduction to form a hydroxyl group. Lys88 hydrogen-bonds to Tyr58 to tune its *pKa* and enforce geometry, whereas Asp53 forms a salt bridge with Lys88 to modulate overall electrostatics and facilitate proton donation. (**B**) The spatial conservation of the active-site tetrad allows the possibility for an alternative 1,4 reduction mechanism, which cannot be ruled out based on the hydride transfer to C17. In this case, the substrate ester (enol) forms a hydrogen bond with His121, which in turn acts as a general acid and proton donor upon hydride transfer by NADPH. Dashed lines indicate hydrogen bonds.

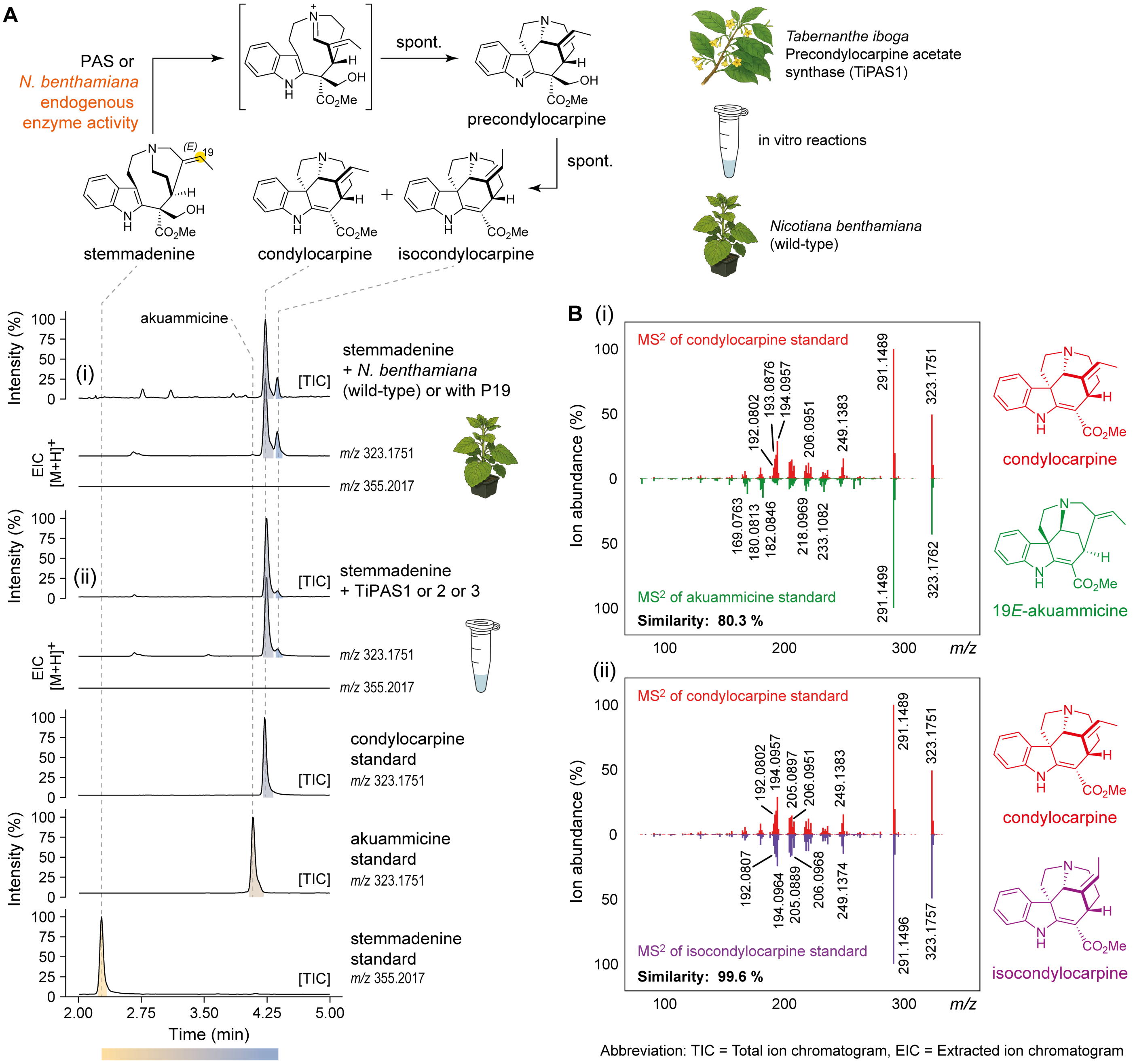


**Figure S23. Biosynthesis of condylocarpine and isocondylocarpine in *Nicotiana benthamiana*.** (**A**) (i) LC-MS chromatograms showing endogenous *N. benthamiana* metabolic activity resulting in the oxidation of stemmadenine to form condylocarpine and a putative isomer, isocondylocarpine. When stemmadenine is present in *N. benthamiana* either as a product of pathway reconstitution or following exogenous feeding, it is progressively oxidized to these products over time. (ii) LC-MS chromatograms of in vitro enzymatic assays with heterologously expressed *Tabernanthe iboga* precondylocarpine acetate synthase homologs (TiPAS1-3), incubated with stemmadenine. TiPAS homologs can catalyze the oxidation of stemmadenine to yield condylocarpine and isocondylocarpine. This suggests that the oxidation observed in *N. benthamiana* host may be mediated by a berberine bridge enzyme-like (BBL) oxidases, which belong to the same family as TiPAS (13). The structure of condylocarpine was confirmed in this study by purification and NMR analysis, while the minor product isocondylocarpine was tentatively annotated based on MS$^2$ fragmentation spectrum and retention time. (**B**) (i) MS$^2$ spectral comparison of condylocarpine and akuammicine, both metabolic sink products of the same *m/z* 323.1751 $(M+H)^+$ due to the absence of downstream biosynthetic enzymes. Although these compounds exhibit similar retention times and MS$^2$ fragment ions, close inspection of their MS$^2$ reveals unique diagnostic fragments for condylocarpine. (ii) MS$^2$ spectral comparison between condylocarpine and isocondylocarpine exhibits high similarity in fragmentation pattern, supporting the structural assignment of isocondylocarpine as a stereoisomer of condylocarpine.

**A**

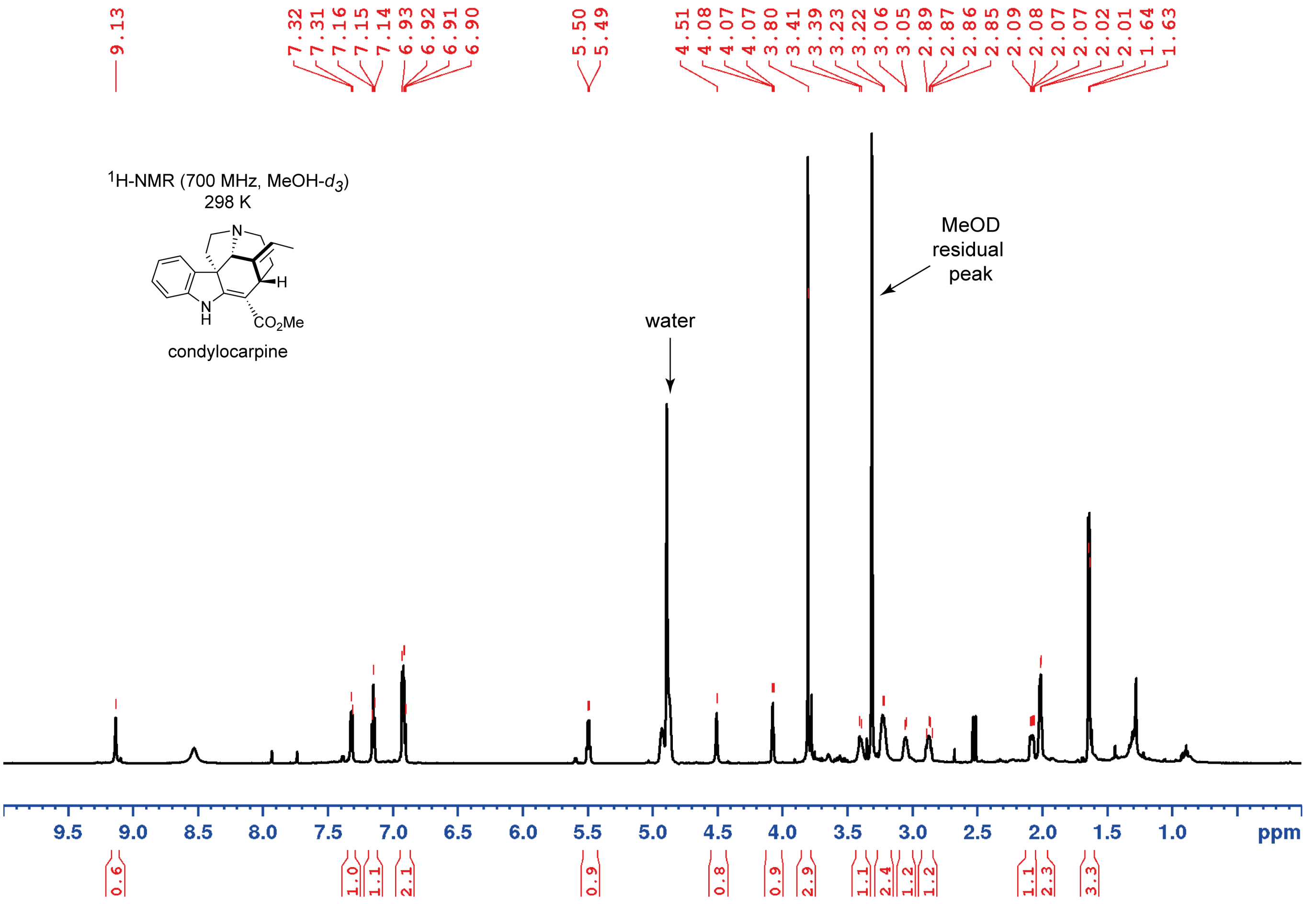


**B**

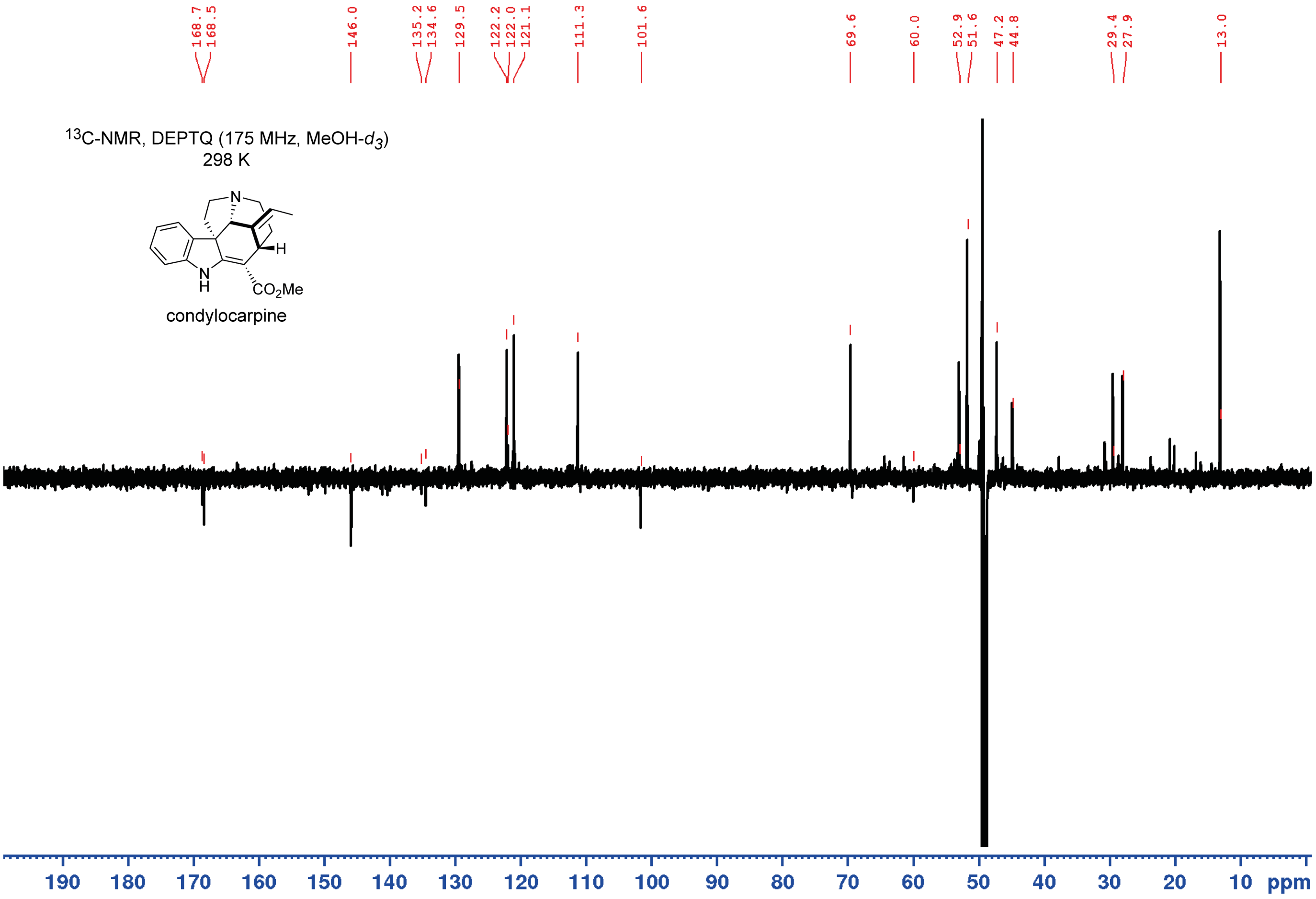


**Figure S24. NMR of condylocarpine.** (**A**) 1H NMR with water presaturation full range in MeOH-$d_3$ of condylocarpine as formate. (**B**) DEPTQ full range in MeOH-$d_3$ of condylocarpine as formate.

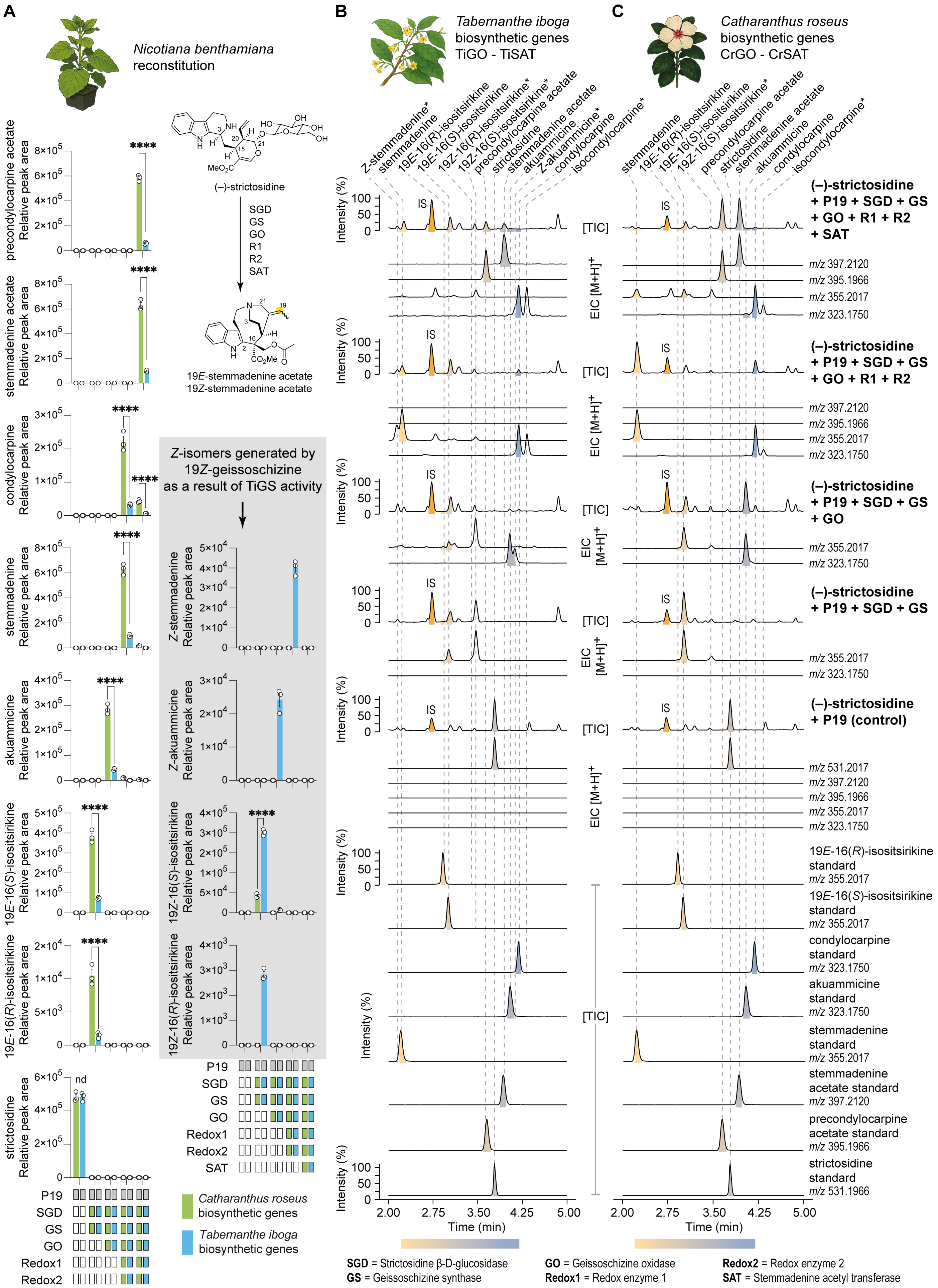


Abbreviation: TIC = Total ion chromatogram, EIC = Extracted ion chromatogram, IS = internal standard
* tentative assignment based on MS$^2$ fragmentation and retention time.

**Figure S25. Pathway reconstitution of stemmadenine acetate biosynthesis from strictosidine in *Nicotiana benthamiana*.** (**A**) Comparison of metabolite production from pathway reconstitution using previously characterized *Catharanthus roseus* stemmadenine acetate biosynthetic genes (green bars) and orthologous *Tabernanthe iboga* genes described in this study (blue bars) in *N. benthamiana*, with strictosidine as the common starting substrate. Stemmadenine acetate production was lower with *T. iboga* genes, primarily due to the activity of TiGS. Unlike CrGS, which exclusively generates the canonical 19*E*-isomer (19*E*-geissoschizine), TiGS activity also leads to the formation of 19Z-isomeric intermediates and products, highlighted in the grey shaded box. (**B**) Representative LC-MS chromatograms of the pathway intermediates and products generated by *T. iboga* biosynthetic genes expressed in *N. benthamiana*, compared against authentic standards (19*E*-isomers). Analytes were identified by comparison with authentic standards of the 19*E*-configuration. Compounds with a 19*Z*-configuration were annotated based on $MS^2$ fragmentation patterns and retention time offsets attributable to the *Z*-configuration at C19. (**C**) Representative LC-MS chromatograms of the corresponding pathway intermediates and products from reconstitution with *C. roseus* biosynthetic genes under identical conditions. For all intermediates, data are represented as the mean ± standard deviation (*n* = 3 biological replicates), based on extracted ion chromatogram peak area ($[M+H]^+$) normalized to the internal standard (IS). Statistical significance was evaluated using an unpaired t-test (*****P* < 0.0001). Bar graphs are ploted as the mean ± S.D.

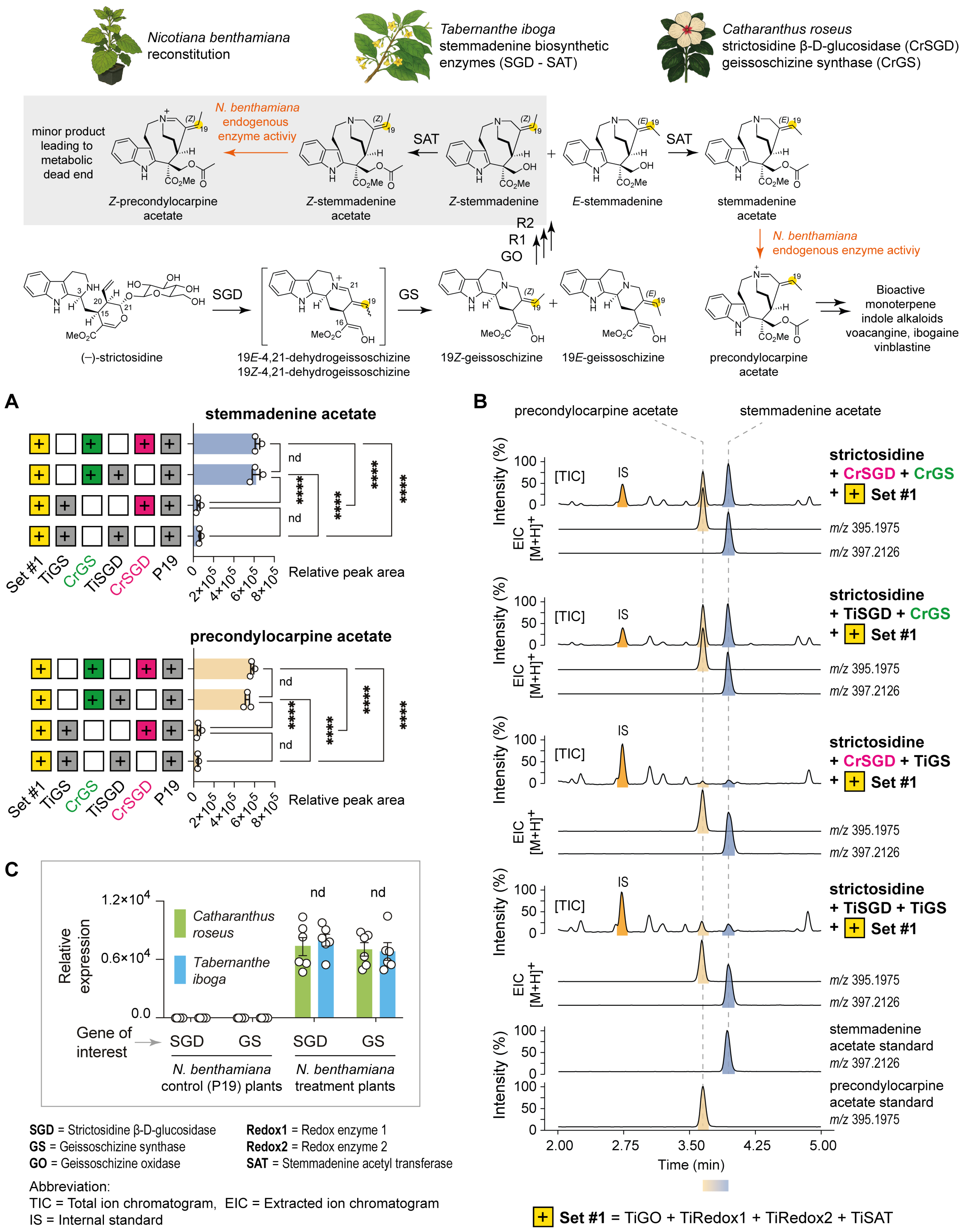


**Figure S26. Functional analysis of strictosidine β-D-glucosidase (SGD) and geissoschizine synthase (GS) in the reconstitution of the stemmadenine acetate biosynthetic pathway in *Nicotiana benthamiana*.** (**A**) Relative peak areas of stemmadenine acetate and precondylocarpine acetate produced in *N. benthamiana* leaf disks following combinatorial pathway reconstitution of *Tabernanthe iboga* multigene construct (pMGC2) with either *T. iboga* or *Catharanthus roseus* SGD and GS enzymes, using strictosidine as the substrate. Peak areas were normalized to an internal standard and are presented as mean ± standard deviation (*n* = 3 biological replicates). Statistical analysis was performed using one-way ANOVA (*****P* < 0.0001; nd. = not significant, *P* > 0.05, n = 3 biological replicates). (**B**) Representative LC-MS chromatograms corresponding to data in panel **A**, comparing product profiles resulting from different combinations of *T. iboga* or *C. roseus* SGD and GS enzymes in pathway reconstitution assays. (**C**) Relative gene expression of *T. iboga* and *C. roseus* SDG and GS genes transiently expressed in *N. benthamiana* leaves for pathway reconstitution. Gene expression was normalized to the reference gene *N. benthamiana* elongation factor 1α (EF1α) and compared to a P19-only control. Data are shown as mean ± standard deviation (*n* = 6 biological replicates). No significant differences in expression levels were observed between orthologous genes (unpaired *t-test*, nd = not significant *P* > 0.05). Bar graphs are ploted as the mean ± S.D.

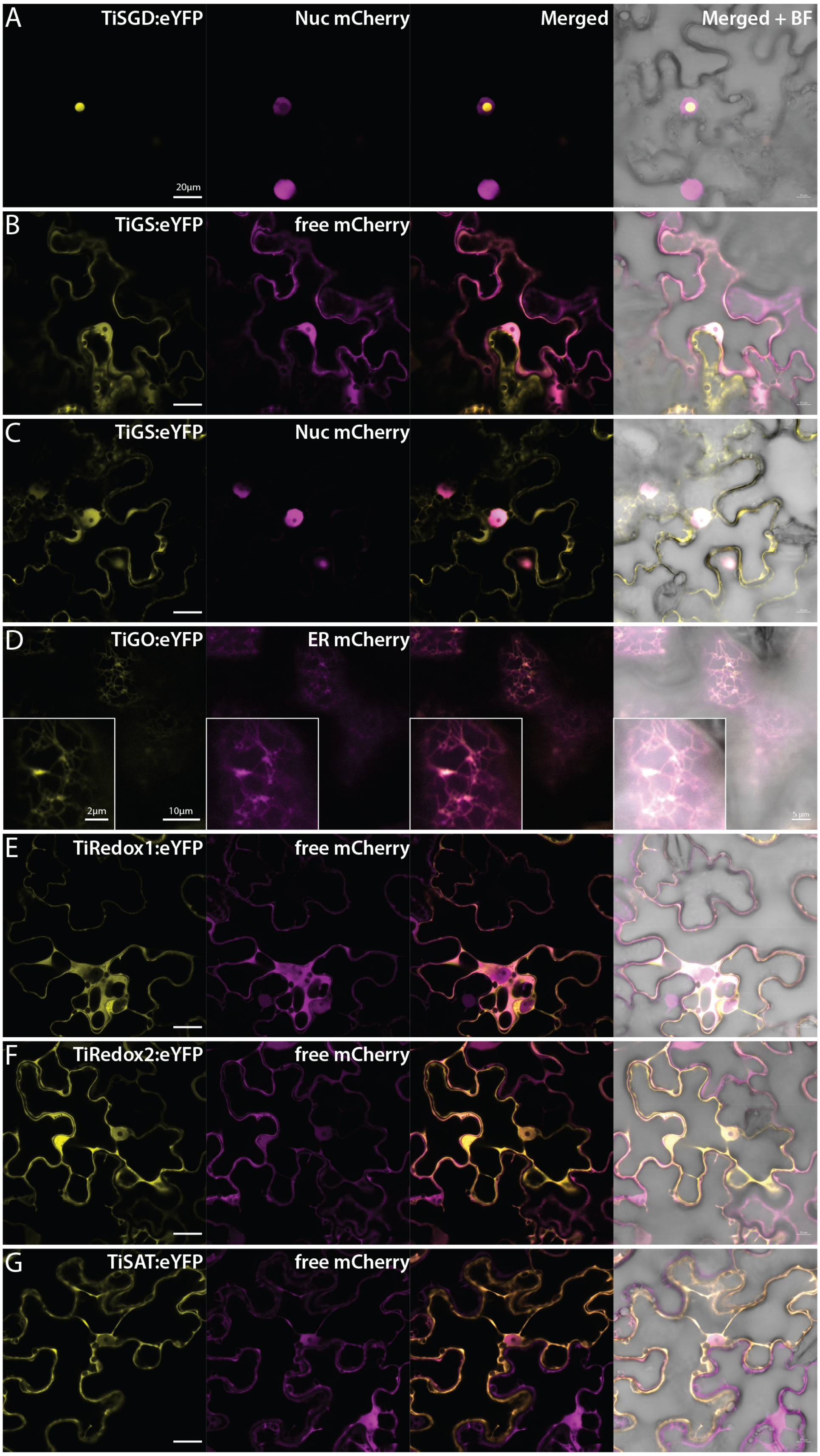


**Figure S27. Subcellular localization of *Tabernanthe iboga* stemmadenine acetate biosynthetic enzymes in *Nicotiana benthamiana* leaves.** *T. iboga* biosynthetic enzymes were C-terminally fused to enhanced yellow fluorescent protein (eYFP) and transiently co-expressed with subcellular localization marker in *N. benthamiana* leaves. Co-infiltration was performed with subcellular organelle markers to mCherry (red fluorescent protein) to determine localization (14). (**A**) *T. iboga* strictosidine β-D-glucosidase localizes to the nucleus. (**B, C**) Geissoschizine synthase (TiGS) localizes both to the nucleus and cytoplasm. (**D**) Geissoschizine oxidase (TiGO) localizes to the endoplasmic reticulum (ER). (**E**) Redox enzyme 1 (TiRedox1) localizes to the cytoplasm. (**F**) Redox enzyme 2 (TiRedox2) localizes to the cytoplasm. (**G**) Stemmadenine acetyl transferase (TiSAT) localizes to the cytoplasm.

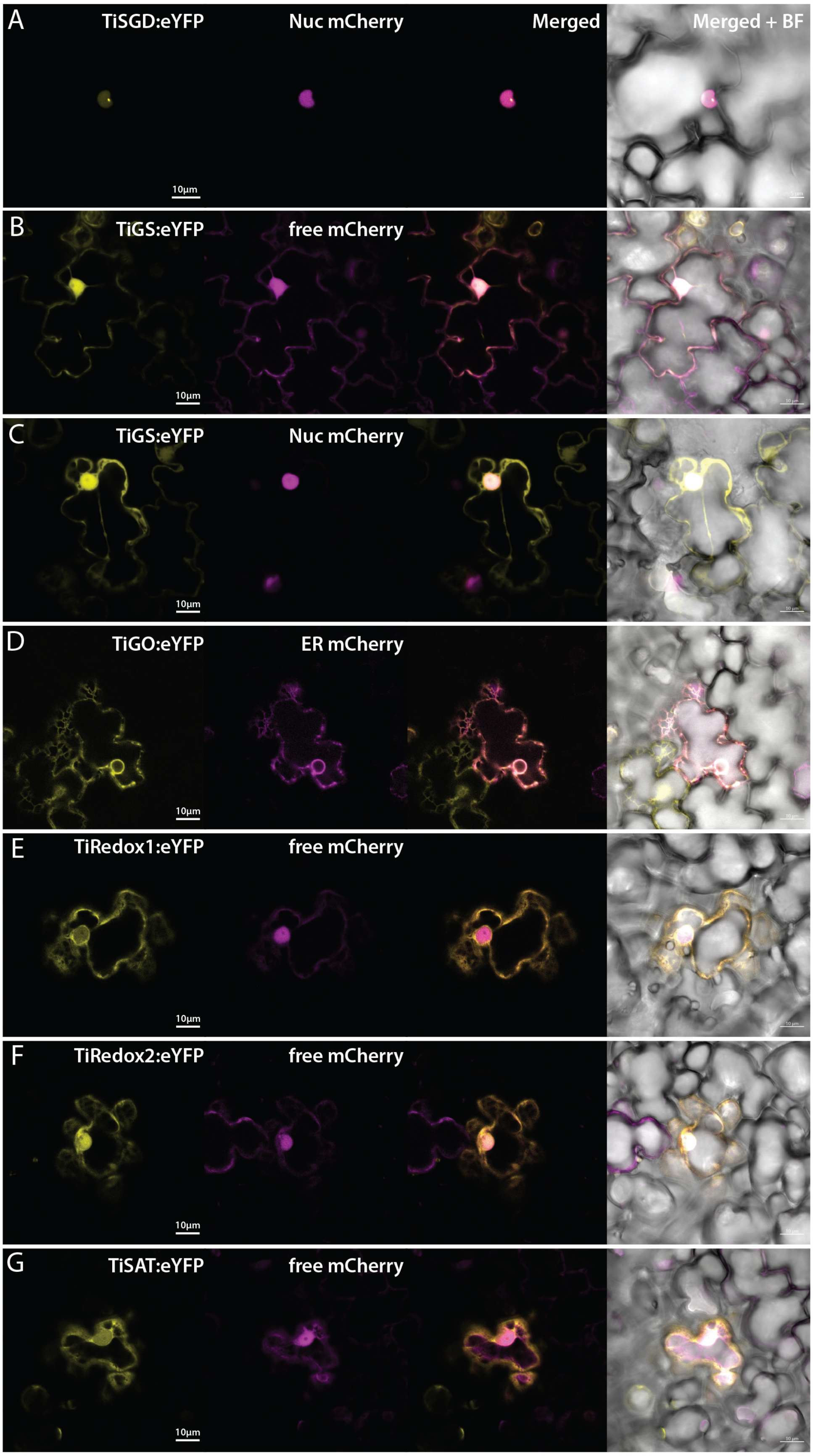


**Figure S28. Subcellular localization of *Tabernanthe iboga* stemmadenine acetate biosynthetic enzymes in *Catharanthus roseus* flower petals.** *T. iboga* biosynthetic enzymes were C-terminally fused to enhanced yellow fluorescent protein (eYFP) and transiently expressed with subcellular localization marker in *C. roseus* flower petals. Co-infiltration was performed with subcellular organelle markers to mCherry (red fluorescent protein) to determine localization (14). (**A**) *T. iboga* strictosidine β-D-glucosidase localizes to the nucleus. (**B, C**) Geissoschizine synthase (TiGS) localizes both to the nucleus and cytoplasm. (**D**) Geissoschizine oxidase (TiGO) localizes to the endoplasmic reticulum (ER). (**E**) Redox enzyme 1 (TiRedox1) localizes to the cytoplasm. (**F**) Redox enzyme 2 (TiRedox2) localizes to the cytoplasm. (**G**) Stemmadenine acetyl transferase (TiSAT) localizes to the cytoplasm.

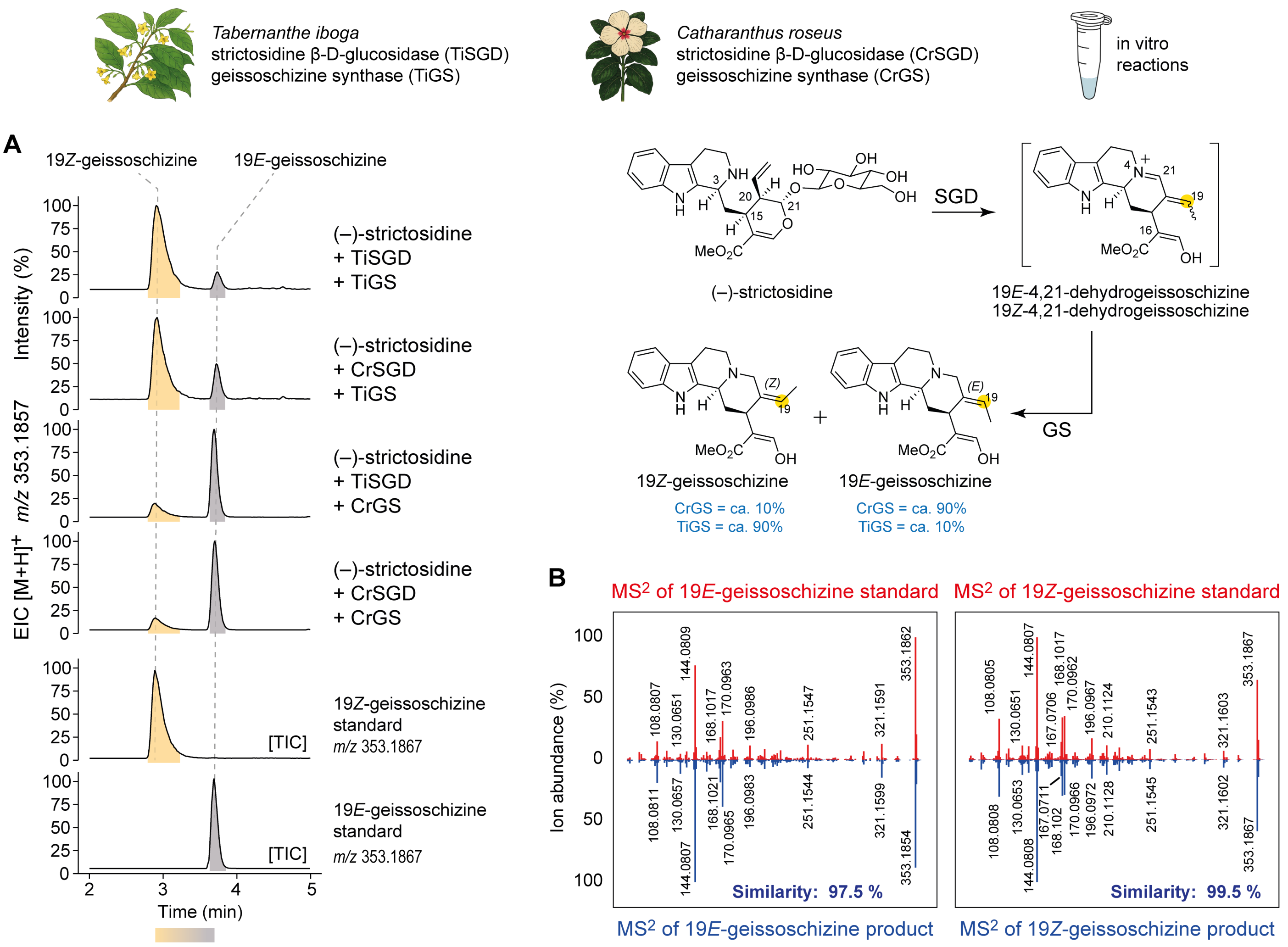


**Figure S29. Functional characterization of *Tabernanthe iboga* geissoschizine synthase (GS) enzyme activity.** (**A**) LC-MS chromatograms of in vitro enzymatic assays using heterologously expressed and purified GS and strictosidine β-D-glucosidase (SGD) enzymes from *T. iboga* and *C. roseus*, assayed with the substrate strictosidine. Coupled enzymatic assays revealed that *T. iboga* GS predominantly produces the 19*Z*-geissoschizine isomer, whereas *C. roseus* GS predominantly yields the 19*E*-geissoschizine isomer. (**B**) MS$^2$ spectra of the 19*E*- and 19*Z*-geissoschizine isomers generated by SGD-GS coupled reactions, shown alongside spectra of authentic synthetic standards.

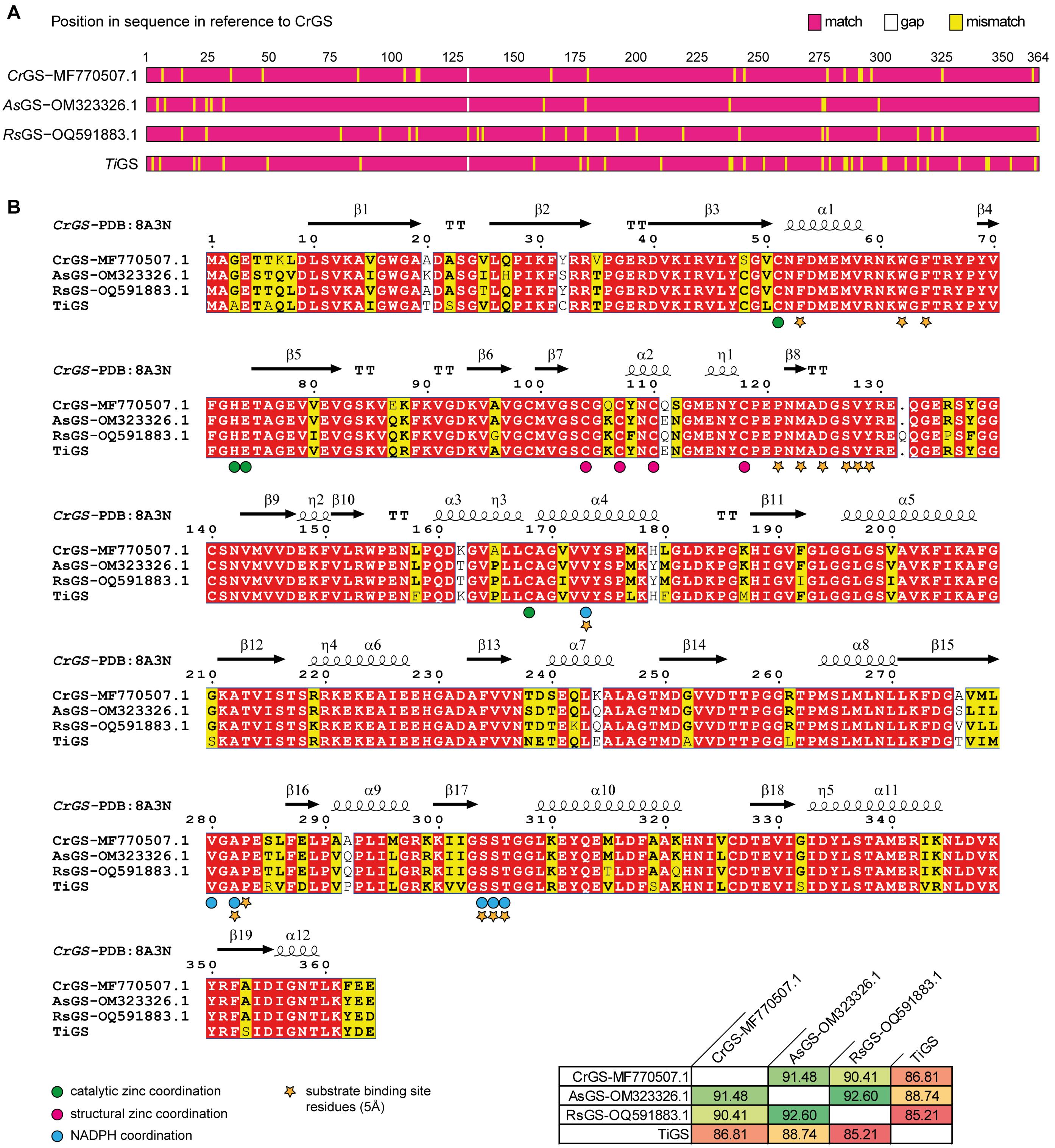


| | CrGS-MF770507.1 | AsGS-OM323326.1 | RsGS-OQ591883.1 | TiGS |
|---|---|---|---|---|
| CrGS-MF770507.1 | | 91.48 | 90.41 | 86.81 |
| AsGS-OM323326.1 | 91.48 | | 92.60 | 88.74 |
| RsGS-OQ591883.1 | 90.41 | 92.60 | | 85.21 |
| TiGS | 86.81 | 88.74 | 85.21 | |

**Figure S30. Multiple sequence alignment of geissoschizine synthase (GS) from monoterpene indole alkaloid (MIA) producing plant species.** (**A**) Schematic representation of amino acid sequence alignment of functionally characterised GS enzymes, members of the medium-chain alcohol dehydrogenase/reductase (MDR) family (15), from various MIA-producing plant species, including *T. iboga* GS. (**B**) Detailed alignment of GS enzyme sequences, annotated with structural features based on *C. roseus* GS crystal structure (16). Key residues involved in zinc coordination (structural and catalytic), NADPH binding, and substrate pocket binding interactions are highlighted (15, 16). The percentage amino acid identity between TiGS and GS enzymes from other plant species is also shown. GenBank accession numbers of the functionally characterized GS enzymes are indicated as appropriate.

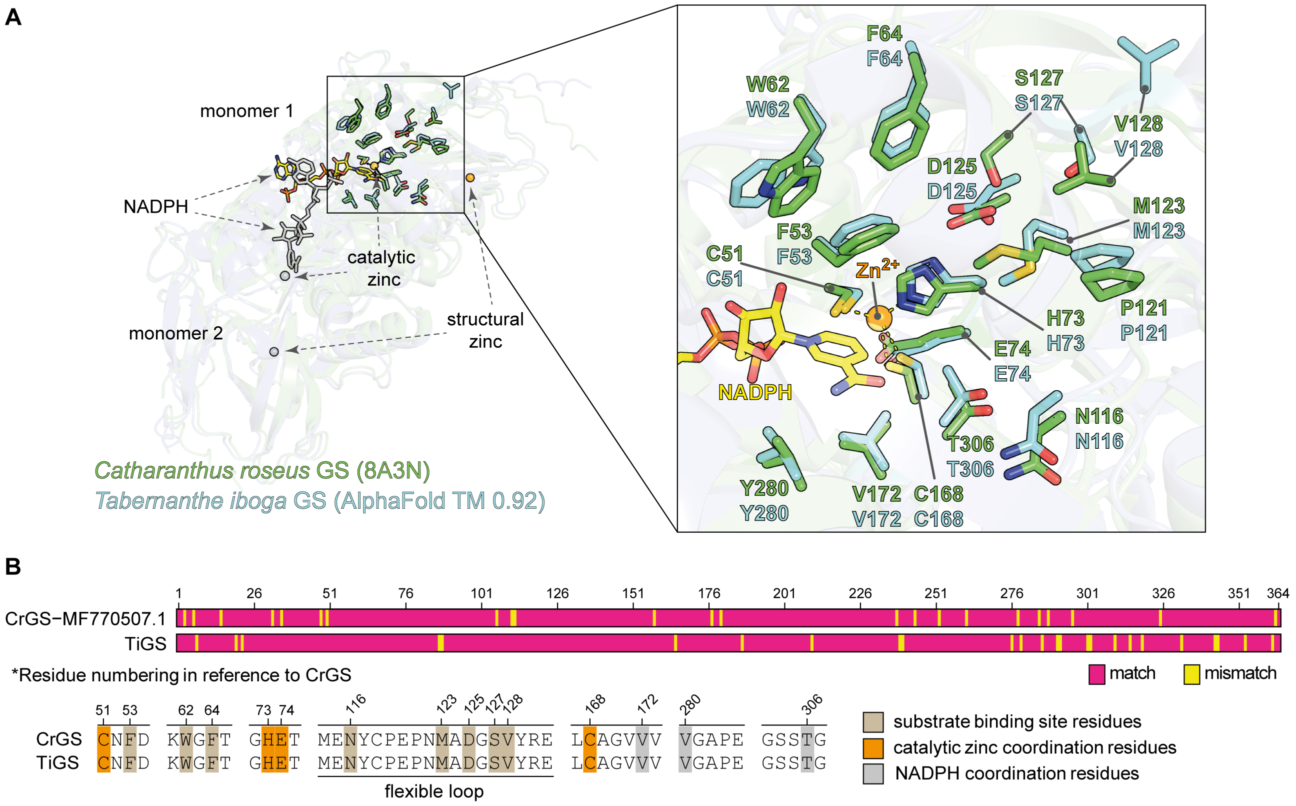


**Figure S31. Structural analysis of *Tabernanthe iboga* geissoschizine synthase (TiGS) model.** (**A**) Superimposed structures of the AlphaFold v3 model of TiGS (TM-score = 0.9261) and the experimentally determined crystal structure of *Catharanthus roseus* geissoschizine synthase (CrGS; PDB 8A3N) (16). The two medium-chain dehydrogenases/reductases (MDR) enzymes share a high structural similarity (RMSD 0.8), with the overall architecture of the dimeric protomers, the active site, catalytic and structural zinc binding sites, as well as NADPH binding being conserved. (**B**) Schematic alignment of TiGS and CrGS amino acid sequences (86.81% identity), highlighting conserved and divergent regions. Residues involved in catalytic zinc coordination (Cys51, His73, Glu74, and Cys168), NADPH coordination, and putative substrate-binding residues are indicated (16). No significant variation was observed in the catalytic pocket residues between TiGS and CrGS.

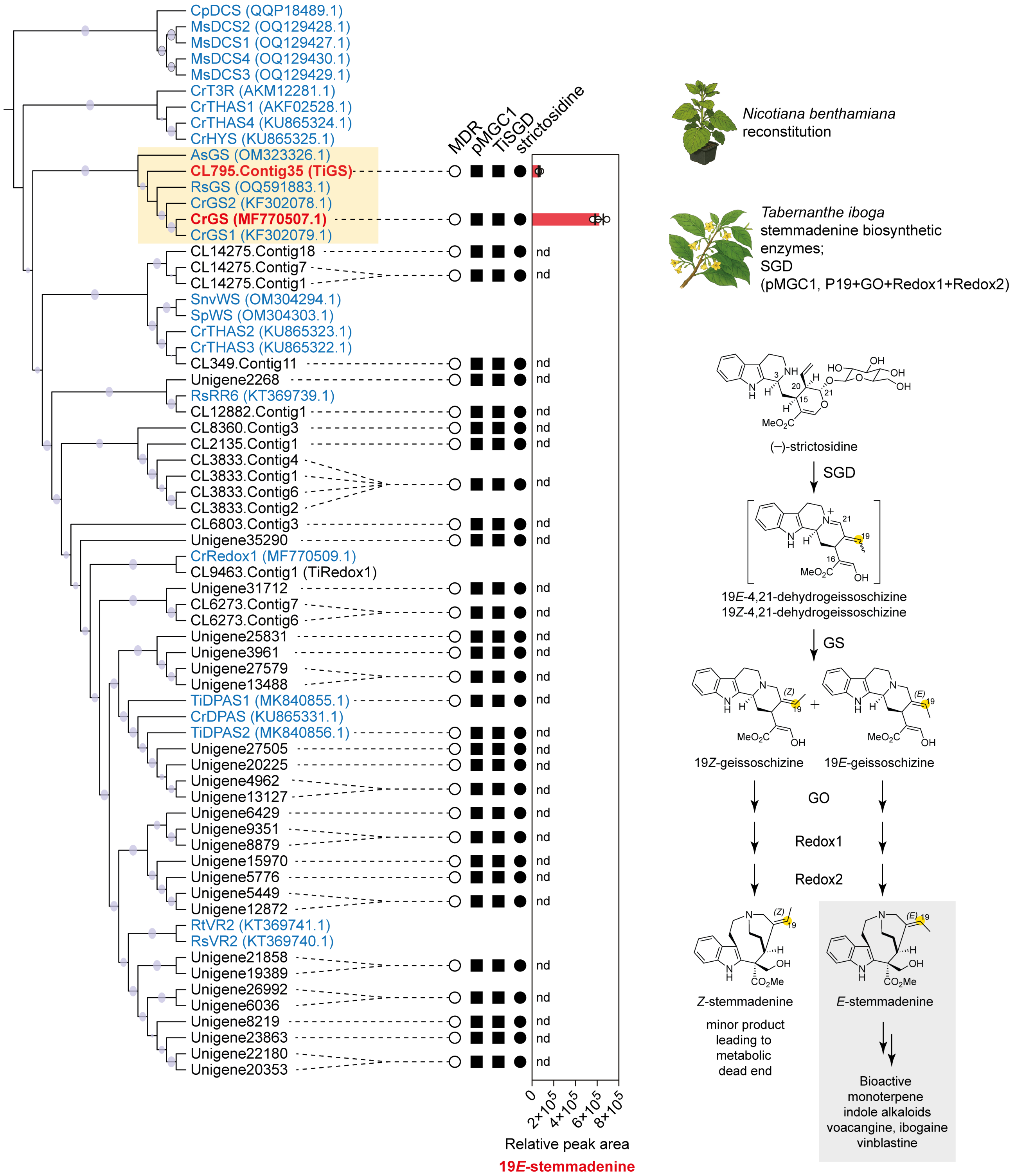


**Figure S32. Screening *Tabernanthe iboga* MDRs for geissoschizine synthase-like activity.** Medium chain dehydrogenase/reductase (MDR) candidates implicated in iminium reduction tested for geissoschizine synthase (GS) activity by transient expression in *N. benthamiana*. Co-expression of multigene construct pMGC1 harbouring p19, TiGO, TiRedox1, and TiRedox2, with TiSGD and a single MDR candidate, followed by strictosidine (substrate) feeding. GS activity assessed by LC-MS detection of stemmadenine, where CrGS served as a positive control. Bar graphs represent peak areas normalized to an internal standard (mean ± standard deviation, $n$ = 3 biological replicates).

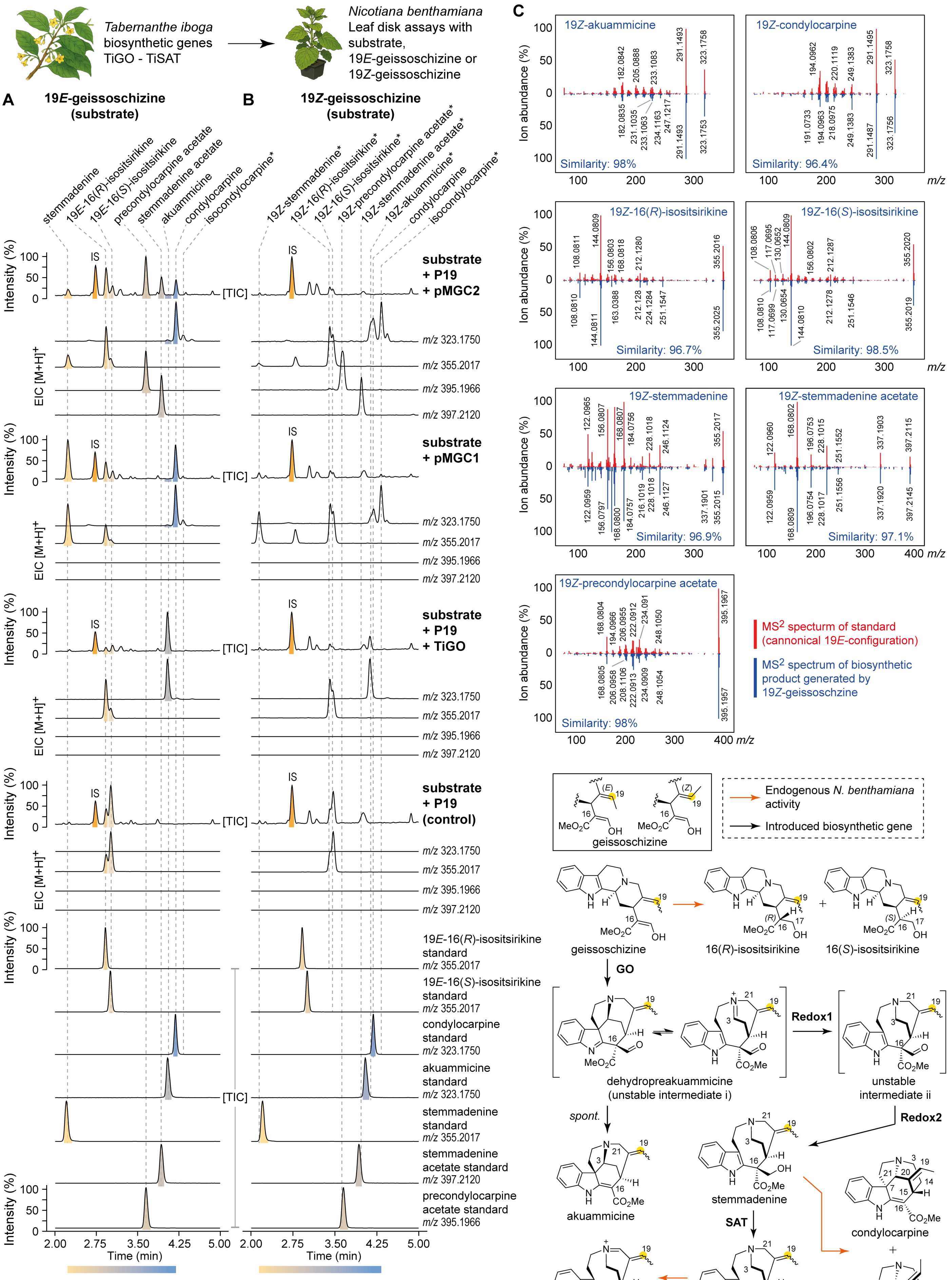


**pMGC2** = multigene consturct containing TiGO + TiRedox1 + TiRedox2 + TiSAT
**pMGC1** = multigene consturct containing TiGO + TiRedox1 + TiRedox2 + P19

Abbreviation: TIC = Total ion chromatogram, EIC = Extracted ion chromatogram, IS = internal standard
* tentative assignment based on MS$^2$ fragmentation and retention time.

**Figure S33. Pathway reconstitution of *Tabernanthe iboga* stemmadenine acetate biosynthetic genes in *Nicotiana benthamiana* and biosynthetic outcomes following 19*E*- and 19*Z*-geissoschizine substrate feeding.** Pathway reconstitution in *N. benthamiana* was performed by transient expression of multigene constructs encoding *T. iboga* stemmadenine acetate biosynthetic enzymes, followed by substrate feeding with 19*E*- or 19*Z*-geissoschizine. (**A**) LC-MS chromatograms of biosynthetic products obtained from 19*E*-geissoschizine feeding. Canonical products of the pathway, including stemmadenine acetate and its biosynthetic intermediates, are detected. (**B**) LC-MS chromatograms of products generated upon 19Z-geissoschizine feeding. 19Z-isomeric analogs of stemmadenine acetate pathway intermediates and products are detected, albeit at lower levels. (**C**) Comparison of $MS^2$ fragmentation spectra of representative 19*E*- and 19*Z*-isomeric products. $MS^2$ spectra of 19*Z*-isomers show high similarity to those of their 19*E* counterparts, supporting structural assignment. Reaction products were annotated by comparison to authentic 19*E*-isomer standards based on diagnostic $MS^2$ fragmentation and expected retention time offsets arising from the 19Z-configuration at C19.

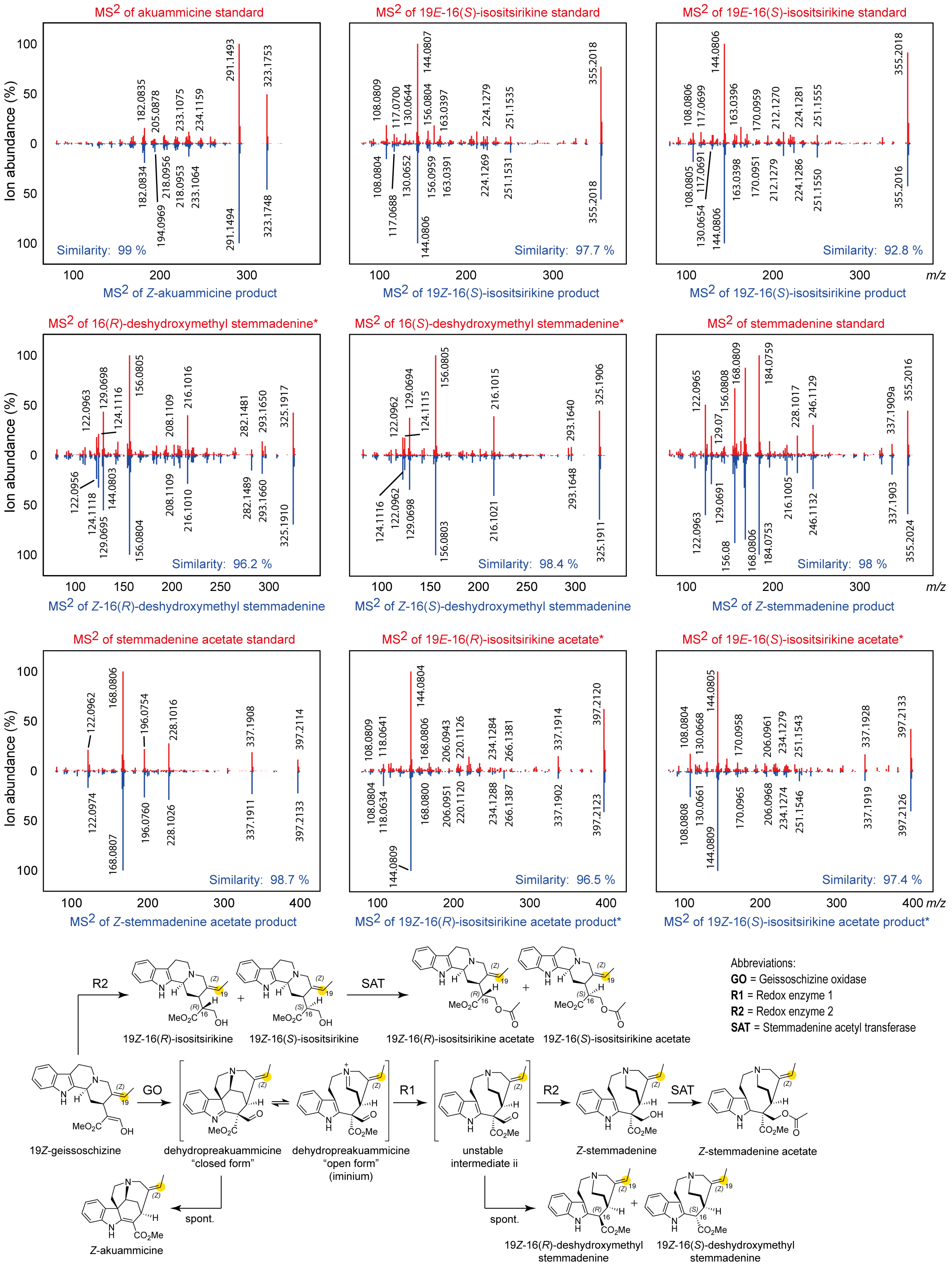


**Figure S34. MS² spectra of 19*Z*-isomers generated by the 19Z-geissoschizne.** MS² fragmentation spectra of 19*Z*-isomeric products generated from 19*Z*-geissoschzine following in vitro cascade reactions of *T. iboga* stemmadenine acetate biosynthetic genes, compared with MS² spectra of the corresponding 19*E*-isomer authentic standards produced by the canonical *Catharanthus roseus* MIA biosynthetic pathway.

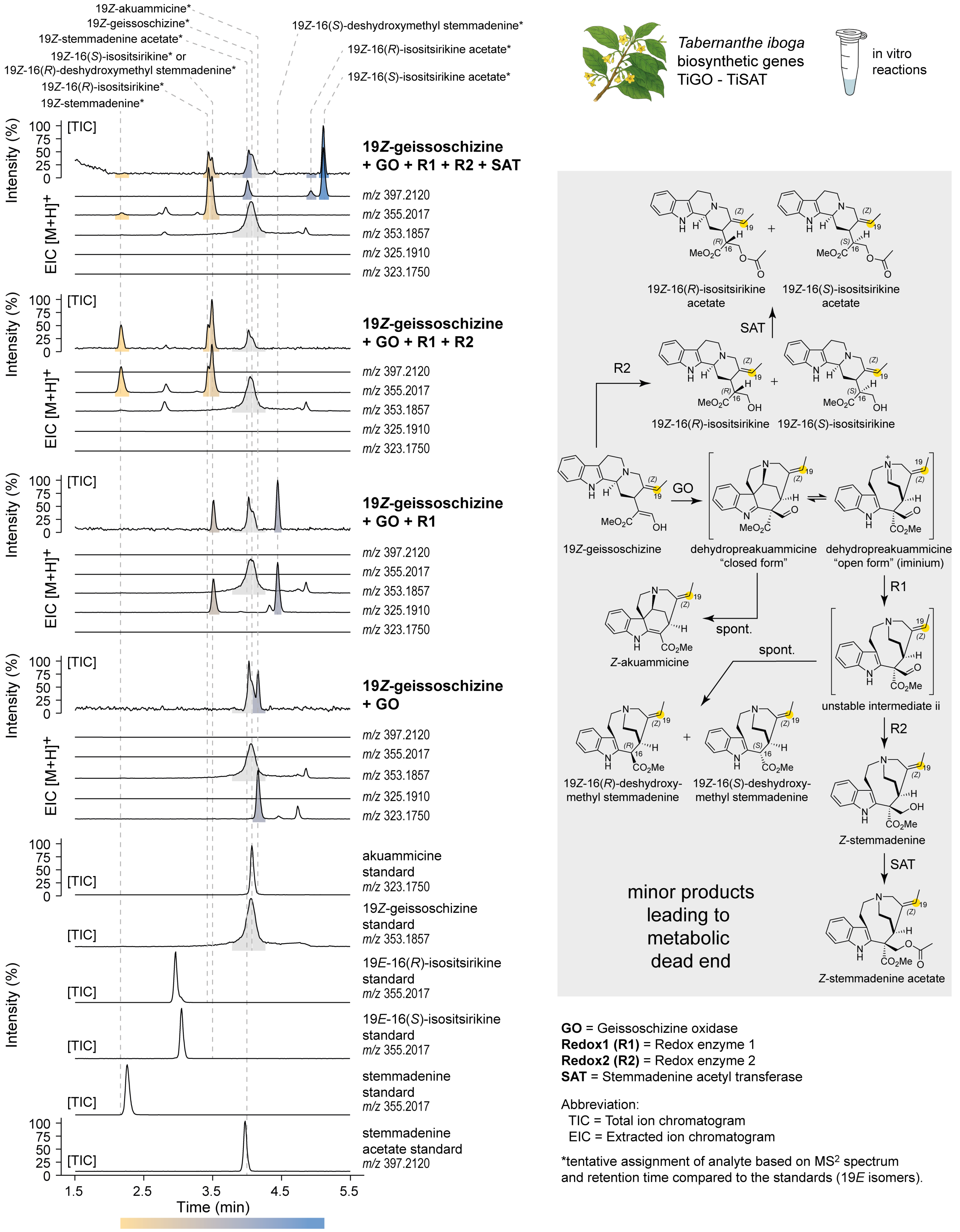


**Figure S35. Stepwise in vitro reconstitution of 19*Z*-isomeric stemmadenine acetate pathway intermediates from 19*Z*-geissoschizine using *Tabernanthe iboga* biosynthetic enzymes.** LC-MS chromatograms showing the sequential enzymatic conversion of 19*Z*-geissoschizine to 19*Z*-stemmadenine acetate via the action of *T. iboga* GO, Redox 1, Redox2, and SAT. The 19*Z*-isomeric products were produced at low levels, as evidenced by TIC and EIC traces, and are not biosynthetic precursors of bioactive compounds ibogaine and voacangine in *T. iboga*. Enzymes were incubated in combination to demonstrate stepwise pathway progression from 19*E*-geissoschizine substrate. Reaction products were annotated by comparison to authentic 19*E*-isomer standards based on $MS^2$ fragmentation and retention time offsets attributable to the 19Z-configuration at C19.

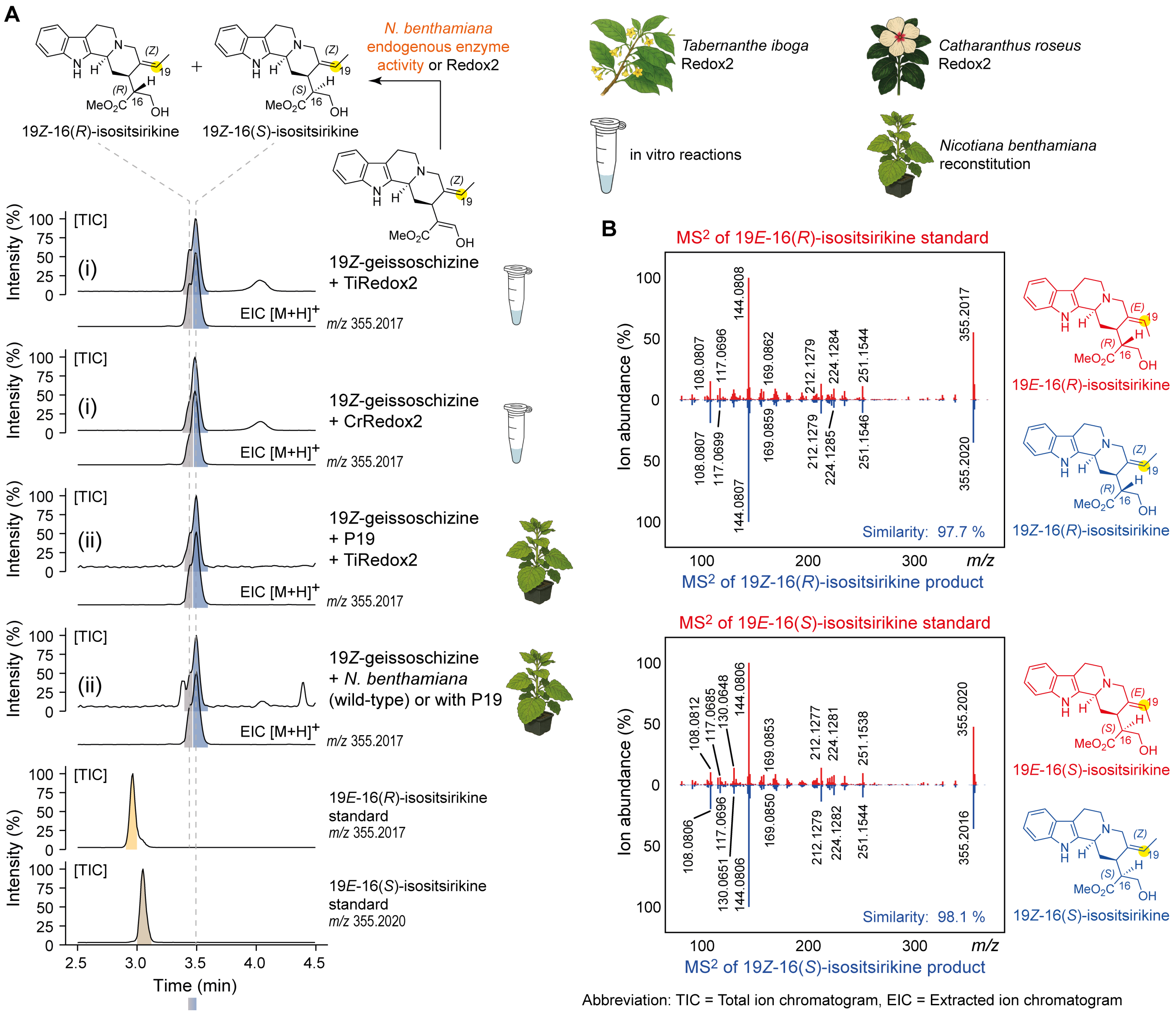


**Figure S36. Functional characterization of *Tabernanthe iboga* redox enzyme 2 (TiRedox2) activity using 19Z-geissoschizine**. (A) (i) LC-MS chromatograms of in vitro enzymatic assays using heterologously expressed TiRedox2 and *Catharanthus roseus* (CrRedox2), assayed with 19*Z*-geissoschizine as the substrate. Both enzymes catalyzed the reduction of 19*Z*-geissoschizine to the isomeric products 19*Z*-16(*R*)-isositsirikine and 19*Z*-16(*S*)-isositsirikine. (ii) LC-MS chromatograms showing that endogenous *Nicotiana benthamiana* metabolic activity also reduces 19*Z*-geissoschizine to the identical 19*Z*-isositsirikine isomers. (**B**) MS$^2$ spectra of the 19*Z*-16(*R*)-isositsirikine and 19*Z*-16(*S*)-isositsirikine isomers produced by TiRedox2 exhibited high spectral similarity to the authentic 19*E*-isomers standards, supporting the tentative structural assignment of the 19*Z*-isomers.

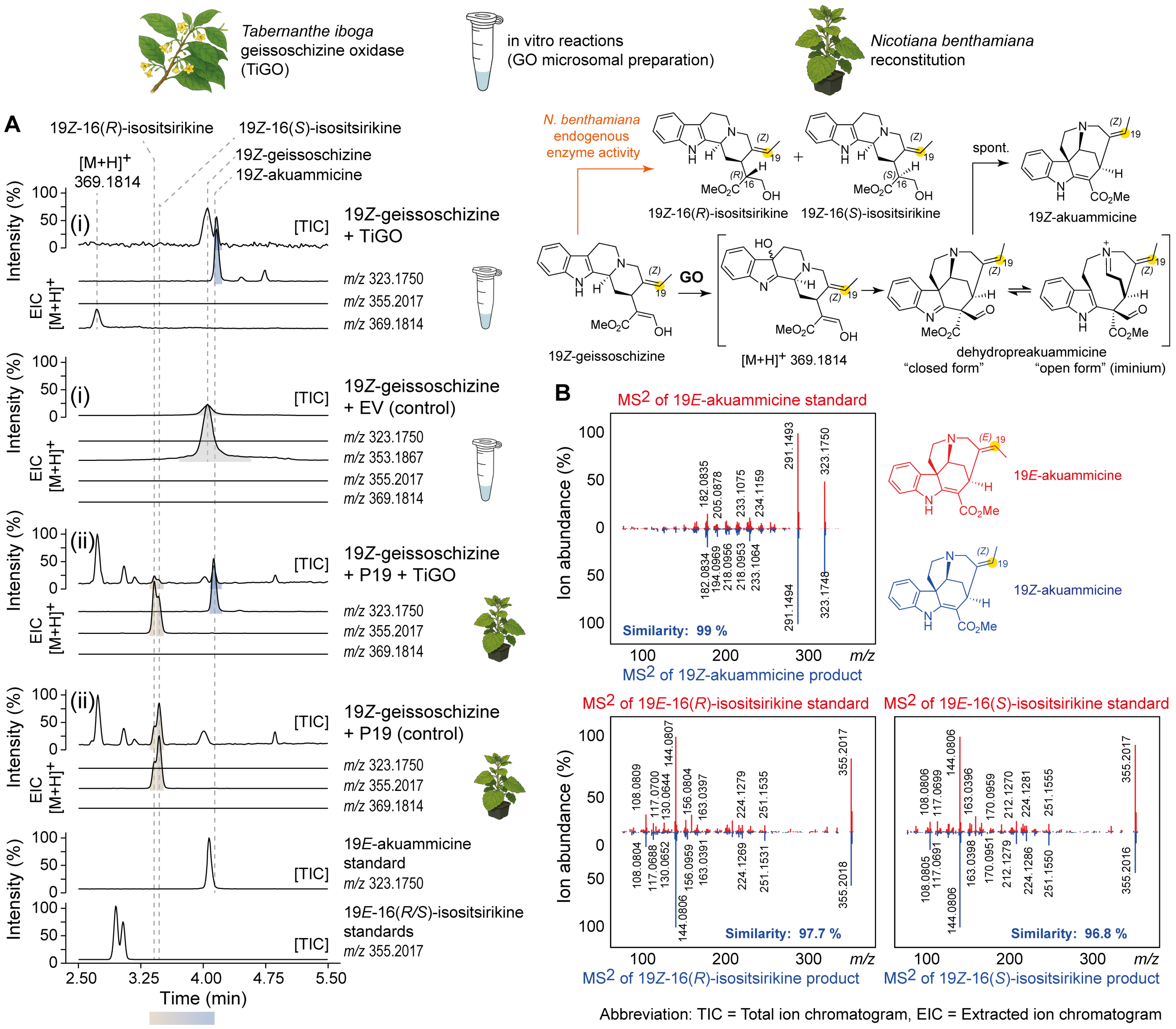


**Figure S37. Functional characterization of *Tabernanthe iboga* geissoschizine oxidase (TiGO) activity with 19*Z*-geissoschizine.** (**A**) (i) LC-MS chromatograms of in vitro enzymatic assays using yeast microsomes expressing TiGO, incubated with 19*Z*-geissoschizine as substrate. (ii) LC-MS chromatograms of TiGO transiently expressed in *Nicotiana benthamiana* with the P19 viral silencing suppressor, assayed in leaf discs supplied with 19*Z*-geissoschizine. In both systems, TiGO catalysed the oxidation of 19*Z*-geissoschizine to produce the thermodynamic product 19*Z*-akuammicine in the absence of downstream biosynthetic enzymes. The endogenous *N. benthamiana* metabolic activity resulted in the reduction of 19*Z*-geissoschizine to 19*Z*-isositsirikine isomers. (**B**) MS$^2$ spectrum of TiGO generated 19*Z*-configured products from both in vivo and in vitro assays, compared with spectra of authentic 19*E*-configured standards.

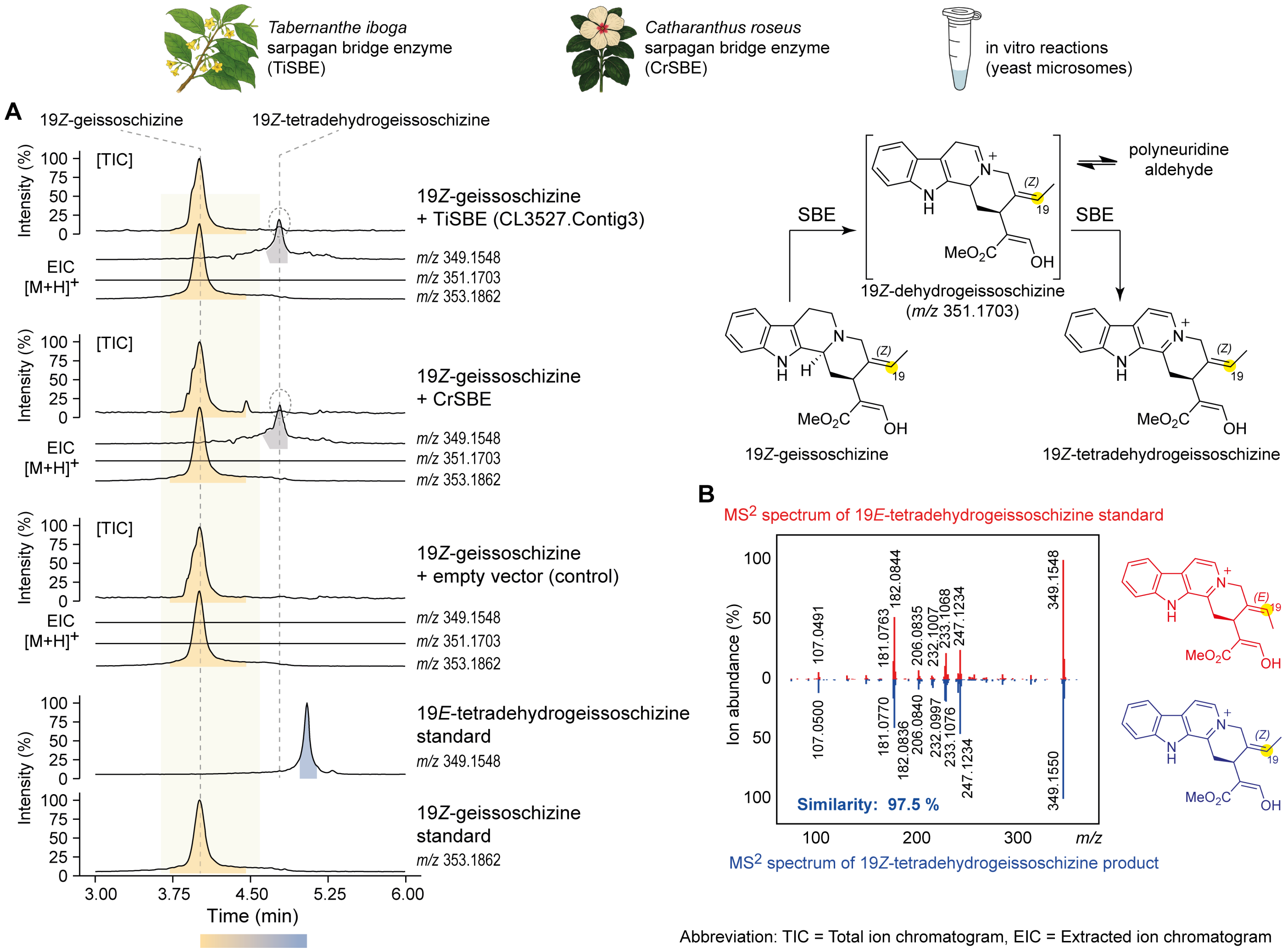


**Figure S38. Functional characterization of *Tabernanthe iboga* sarpagan bridge enzyme (TiSBE) activity in vitro using 19*Z*-geissoschizine.** (**A**) LC-MS chromatograms of in vitro enzymatic assays using yeast microsomes expressing TiSBE or *Catharanthus roseus* SBE (CrSBE). Microsomal fractions were assayed with 19*Z*-geissoschizine as substrate under identical conditions. Both enzymes catalyzed the formation of 19*Z*-tetradehydrogeissoschizine at very low amounts. (**B**) $MS^2$ spectrum of 19*Z*-tetradehydrogeissoschizine produced by TiSBE, compared with that of the authentic standard of 19*E*-variant.

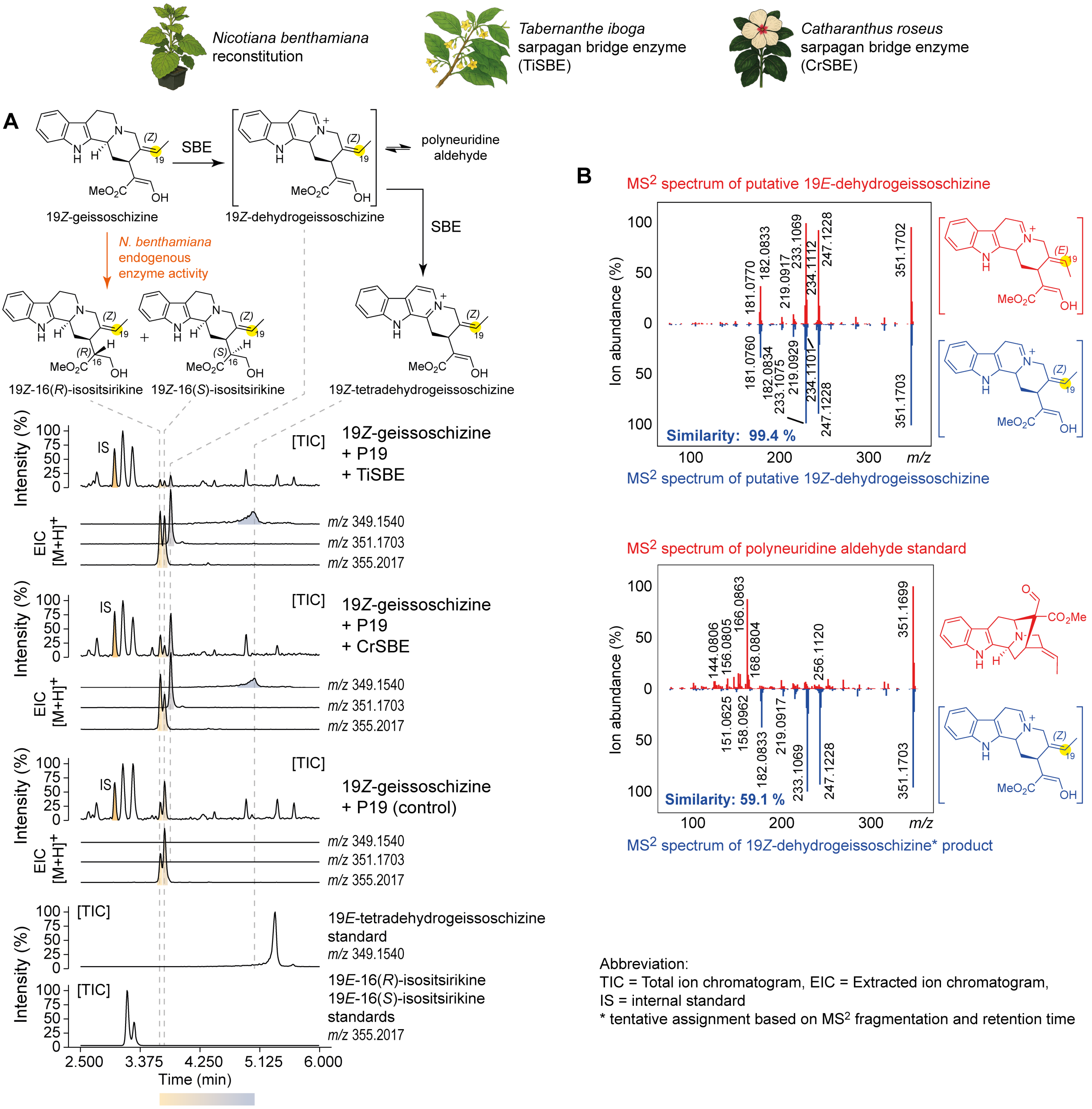


**Figure S39. Functional characterization of *Tabernanthe iboga* sarpagan bridge enzyme (TiSBE) activity in *Nicotiana benthamiana* using 19Z-geissoschizine as substrate.** (**A**) LC-MS chromatograms of TiSBE transiently expressed in *N. benthamiana* with the P19 viral silencing suppressor, assayed in leaf disks supplied with the substrate 19*Z*-geissoschizine. The previously characterised *Catharanthus roseus* SBE (CrSBE) (5) was included as a positive control under identical conditions. Both CrSBE and TiSBE catalyzed the formation of 19*Z*-tetradehydrogeissoschizine via the unstable intermediate 19*Z*-dehydrogeissoschizine, albeit at low abundance. Notably, the endogenous activity of *N. benthamiana* metabolic activity resulted in the reduction of 19*Z*-geissoschizine to 19*Z*-isositsirikine isomers. These 19Z-isomers were annotated based on MS$^2$ spectral similarity and retention times comparisons to the corresponding 19*E*-structural variants, for which authentic standards were available. (**B**) The MS$^2$ spectrum of the 19*Z*-dehydrogeissoschizine showed high similarity to that of 19*E*-dehydrogeissoschizine. This spectrum also differed from polyneuridine aldehyde, which could exist in equilibrium with 19*Z*-dehydrogeissoschizine, further supporting its structural annotation.
